# Gut bacteria degrade purines *via* the 2,8-dioxopurine pathway

**DOI:** 10.1101/2025.04.24.650524

**Authors:** Yuanyuan Liu, Zhiwei Zhou, J. Bryce Jarman, Haoqing Chen, Michelle Miranda-Velez, Robert Terkeltaub, Dylan Dodd

## Abstract

The loss of uricase during hominid evolution has predisposed humans to hyperuricemia and gout, conditions with high global prevalence. In healthy individuals, approximately one-third of urate is excreted into the intestinal tract^1^ where bacteria consume this purine aiding in its elimination^2,3^. However, the molecular details of purine metabolism in the gut microbiome remain unknown. Here we uncover the 2,8-dioxopurine pathway, a previously unrecognized anaerobic route for purine degradation in gut bacteria. This pathway uses a novel selenium-dependent enzyme, 2,8-dioxopurine dehydrogenase (DOPDH), and seven additional enzymes that together link purine metabolism to ATP generation and short chain fatty acid production. Competition experiments in gnotobiotic mice demonstrate that bacteria harboring this pathway exhibit a fitness advantage, with wild-type bacteria rapidly outcompeting a DOPDH-deficient strain. These findings highlight a host-microbe symbiosis, where host-secreted urate fosters a metabolic niche for bacteria that break it down. By elucidating this pathway, our study provides new avenues to modify and enhance intestinal elimination of urate, which has therapeutic implications for treating hyperuricemia and gout, conditions that affect millions of adults worldwide.

---

Uric acid (and its conjugate base at physiological pH, urate) is an intermediate in purine catabolism in organisms across all domains of life. In most mammals, the enzyme uricase converts urate to freely soluble allantoin which is excreted in the urine. However, during hominid evolution, mutations accumulated in the uricase gene, rendering it inactive. The loss of uricase may have conferred evolutionary advantages to early hominids^4,5^ but now poses a risk, as monosodium urate (MSU) has a low solubility limit (∼6.8 mg/dL)^6^, near the mean normal serum urate level of 5.5 mg/dL. Elevated serum urate (hyperuricemia) results in MSU crystal deposition, which causes the painful, and highly prevalent inflammatory arthritic disease gout^7^. The collective US and global disease burden is high; for example, ∼21% of US adults have hyperuricemia^8^, and ∼5% have gout^9^.

Radioisotope tracer studies in the 1960s showed that in healthy individuals, approximately two-thirds of urate is eliminated through urine, while the remaining one-third is excreted into the gastrointestinal tract and degraded by gut bacteria^1^. When the kidneys fail, as occurs with chronic kidney disease (CKD), the intestines assume the primary role in urate elimination^10^. Two recent studies identified human gut bacteria capable of consuming urate^2,3^. A metabolic gene cluster encoding enzymes of unknown function was found to be responsible for anaerobic conversion of uric acid to short-chain fatty acids (SCFAs). Loss-of-function and gain-of-function studies in mice demonstrated that bacteria that degrade uric acid and other purines contribute to uric acid homeostasis in the host^2,3^, indicating that these bacteria actively modulate urate elimination from the body. Furthermore, an association was identified between treatment with antibiotics targeting the anaerobic gut microbiota and the incidence of gout in humans^3^. Given the global burden of hyperuricemia and limited treatment options for CKD patients^11^, there is growing interest in leveraging gut bacteria to enhance urate elimination^10,12^. However, the molecular mechanisms underlying purine metabolism in the gut microbiome remain largely uncharacterized. Here we sought to uncover the molecular details of anaerobic purine metabolism in gut bacteria.

## Stable isotope tracing identifies pathway intermediates in anaerobic microbial uric acid metabolism

In previous work, we identified a uric acid-inducible gene cluster in *Clostridium sporogenes* that is required for converting uric acid to short-chain fatty acids (SCFAs)^3^ (**Extended Data Figure 1a-b**). When these genes were disrupted, SCFA production was blocked, yet a substantial amount of uric acid was still consumed (**Extended Data Figure 1c**). This suggested the possibility that pathway intermediates accumulate in the mutants and could be identified by untargeted liquid chromatography–mass spectrometry (LC-MS).

We cultured wild-type and mutant strains of *C. sporogenes* in the presence of either unlabeled uric acid or its [^13^C_5_]-labeled isotopolog and developed a metabolomic workflow to detect unknown metabolites produced from uric acid in the anaerobic mutant cultures (**Figure 1a**). After differential abundance analysis and feature filtering, we identified a series of characteristic molecular features which: (i) were elevated in mutant vs. wild-type cultures (**Extended Data Figure 2a**), (ii) increased in intensity over culture age (**Extended Data Figure 2b**), and (iii) contained retention-time matched isotopologs when mutants were cultured with [^13^C_5_]-labeled uric acid (**Figure 1b; Supplementary Tables 1-6**). For the six genes analyzed, we identified six unique intermediates (UA-2 through UA-7) that were enriched in mutant vs. wild-type *C. sporogenes* cultures (**Figure 1b**).

**Figure 1.**
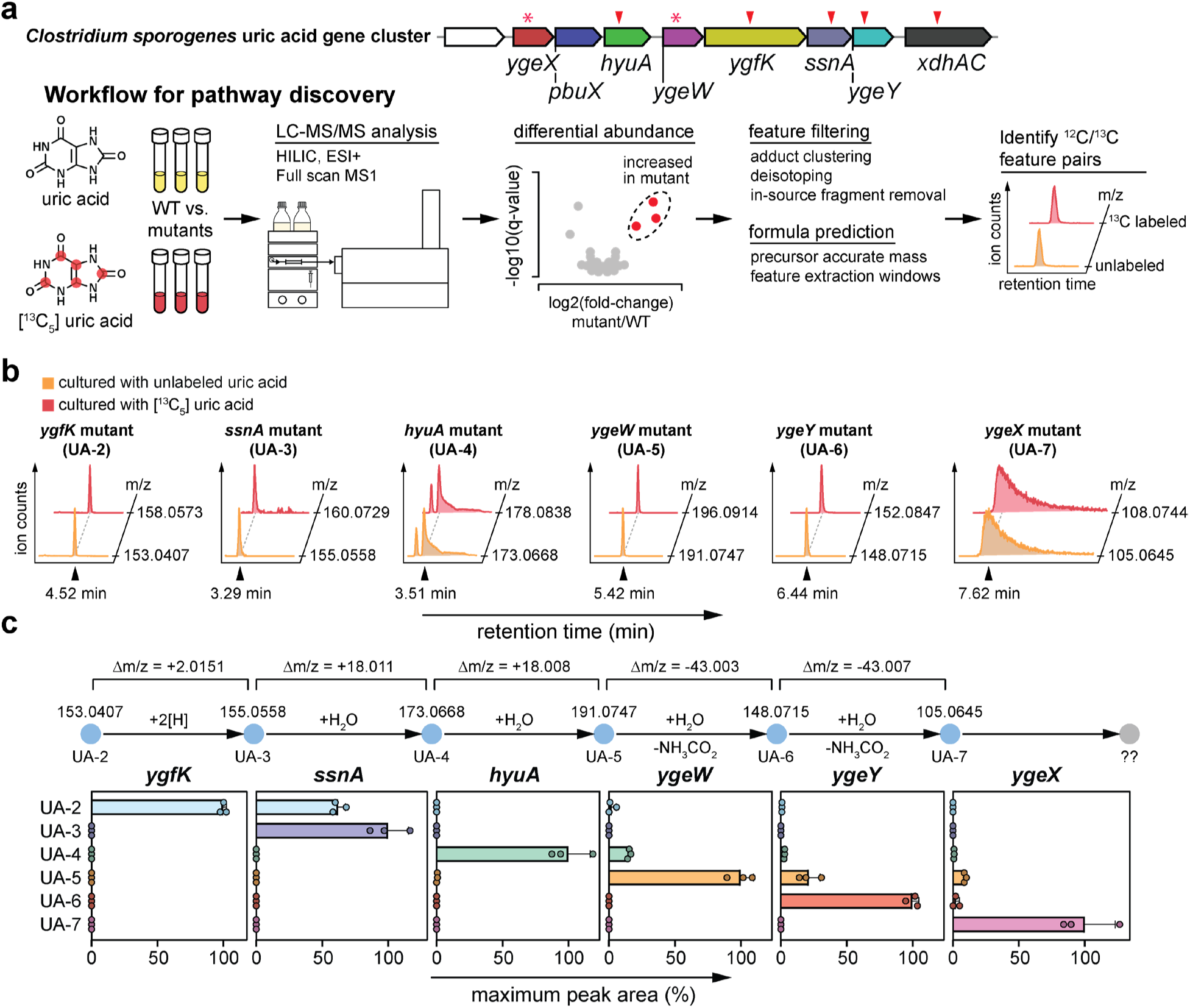
Stable isotope tracing and genetics identify pathway intermediates in anaerobic bacterial uric acid metabolism. (**A**) Schematic showing the gene cluster for *C. sporogenes,* and the workflow for stable isotope tracing and identification of labeled and ^13^C-labeled feature pairs. Arrows indicate ClosTron mutants and asterisks indicate genes that were deleted by CRISPR due to polar effects of ClosTron mutants. (**B**) Extracted ion chromatograms from HILIC LC-MS for the most abundant unlabeled and ^13^C-labeled feature pairs in each of the mutants. (**C**) Combining enzyme family level predictions of mass differences with patterns of intermediate accumulation (by HILIC LC-MS) to place intermediates and genes within the pathway. For **B**, three replicates were performed, and representative data are shown. For **C**, data represent means ± standard deviations from three replicates.

To verify these findings in a separate organism, we examined *Escherichia coli*, a well-established model organism with extensive genetic tools. Analysis of two laboratory adapted K-12 strains (MG1655 and W3110) along with eight “wild-type” strains, revealed presence of a highly conserved uric acid gene cluster (**Extended Data Figure 3a**). Despite this strong sequence conservation (>98% amino acid identity), there was notable variation in uric acid consumption rates. Some strains depleted all uric acid within 24 hours, while others required up to 48 hours for complete consumption (**Extended Data Figure 3b-c**). This variability suggests that differences in gene regulation, rather than intrinsic enzyme activity, contribute to strain-specific differences in uric acid metabolism. Given the genetic tractability of the K-12 MG1655 strain, we selected it for further genetic and metabolomic analyses. Using concentrated cell suspensions to enhance metabolic flux (**Extended Data Figure 4a-b**), we found *E. coli* mutants that corresponded to *C. sporogenes* mutants accumulated the same pathway intermediates (**Extended Data Figure 4c**), independently validating our findings in two phylogenetically distinct organisms.

A pathway for anaerobic purine metabolism was described in the 1950s in *Clostridium cylindrosporum*, an obligate purinolytic bacterium isolated from soil (**Extended Data Figure 1d**)^13–17^ and has recently been revisited^18^. We searched our metabolomic data from *C. sporogenes* and *E. coli* mutants and did not identify these previously proposed intermediates. The one exception was xanthine that accumulated in the *ygfK* mutants, and which we later found not to be xanthine, but instead the isomer 2,8-dioxopurine (*see below*). Coupled with our observation that the uric acid-inducible genes are not present in *C. cylindrosporum*^3^, our findings indicate that the pathway in gut bacteria is entirely distinct. We obtained clean tandem mass spectra for the intermediates in *C. sporogenes* mutants (**Extended Data Figure 5a**) and performed searches of the Mass Databank of North America. These searches returned only one precursor/mass spectrum hit (UA-7 for 2,3-diaminopropanoic acid; **Extended Data Figure 5b**), indicating that most of these intermediates are completely novel.

We reasoned that these six intermediates represent substrates or products for enzymes encoded by the gene cluster. By integrating protein family predictions of mass differences (e.g., peptidase, +H_2_O, Δm/z = +18.0106) and patterns of pathway intermediate accumulation in mutants, we established a framework for the pathway (**Figure 1c**). To verify these predictions, we cloned and over-expressed the corresponding *E. coli* K-12 MG1655 uric acid cluster genes (*ygfK*, *ssnA*, *hyuA*, *ygeW*, *ygeY*, and *ygeX*) in *E. coli* BL-21 (DE3) and purified the proteins by immobilized metal affinity chromatography (iMAC) (**Extended Data Figure 5c**). Larger quantities of pathway intermediates were produced by incubating cell suspensions of each *C. sporogenes* mutant with uric acid and harvesting the supernatant. Then purified proteins were incubated individually with each of the mutant supernatants and pathway intermediates were analyzed by LC-MS. After 3 h enzyme incubation, we observed either depletion of the predicted substrate or accumulation of the predicted product (**Extended Data Figure 5d**). Together, these genetic and biochemical data provide support for the arrangement of enzymes in a new pathway for uric acid metabolism.

## YgeX connects the uric acid pathway to central metabolism via pyruvate

*E. coli* YgeX is known to possess diaminopropionate ammonia-lyase activity^19–21^, yet *E. coli* does not grow with 2,3-diaminopropanoic acid (DPA) and the functional role for the *ygeX* gene is unknown^22^. Using stable isotope tracing (**Extended Data Figure 5e**), we found that both *C. sporogenes* and *E. coli ygeX* mutants accumulate a unique metabolic feature with m/z of 105.0645 (**Extended Data Figure 5f**) which matches authentic DPA in retention time, accurate mass (**Figure 2a**) and mass fragmentation pattern (**Figure 2b**). In the *C. sporogenes ygeX* mutant, DPA accounted for a significant portion of the consumed uric acid, whereas in the *E. coli ygeX* mutant, which had a greater block in uric acid consumption, DPA levels were minimal but detectable (**Figure 2c**). We confirmed results from previous studies^19–21^ showing that recombinant *E. coli* YgeX has diaminopropionate ammonia-lyase activity, converting DPA to pyruvate (**Figure 2d**). Furthermore, incubation of supernatants of *C. sporogenes* and *E. coli ygeX* mutants grown with either labeled or unlabeled uric acid (**Figure 2e**) with recombinant *E. coli* YgeX yielded labeled and unlabeled pyruvate, respectively (**Figure 2f**). These results define the cellular role for YgeX showing that it functions to link uric acid catabolism to pyruvate, a critical node in central metabolism. The observation that uric acid integrates into central metabolism through pyruvate clarifies why phylogenetically diverse bacteria generate distinct SCFA profiles from uric acid^3^. This also stands in sharp contrast to the anaerobic pathway for uric acid metabolism in obligate purinolytic bacteria like *C. cylindrosporum* where formiminoglycine, rather than pyruvate, is the end product of the pathway (**Extended Data Figure 1d**)^13,18^.

**Figure 2.**
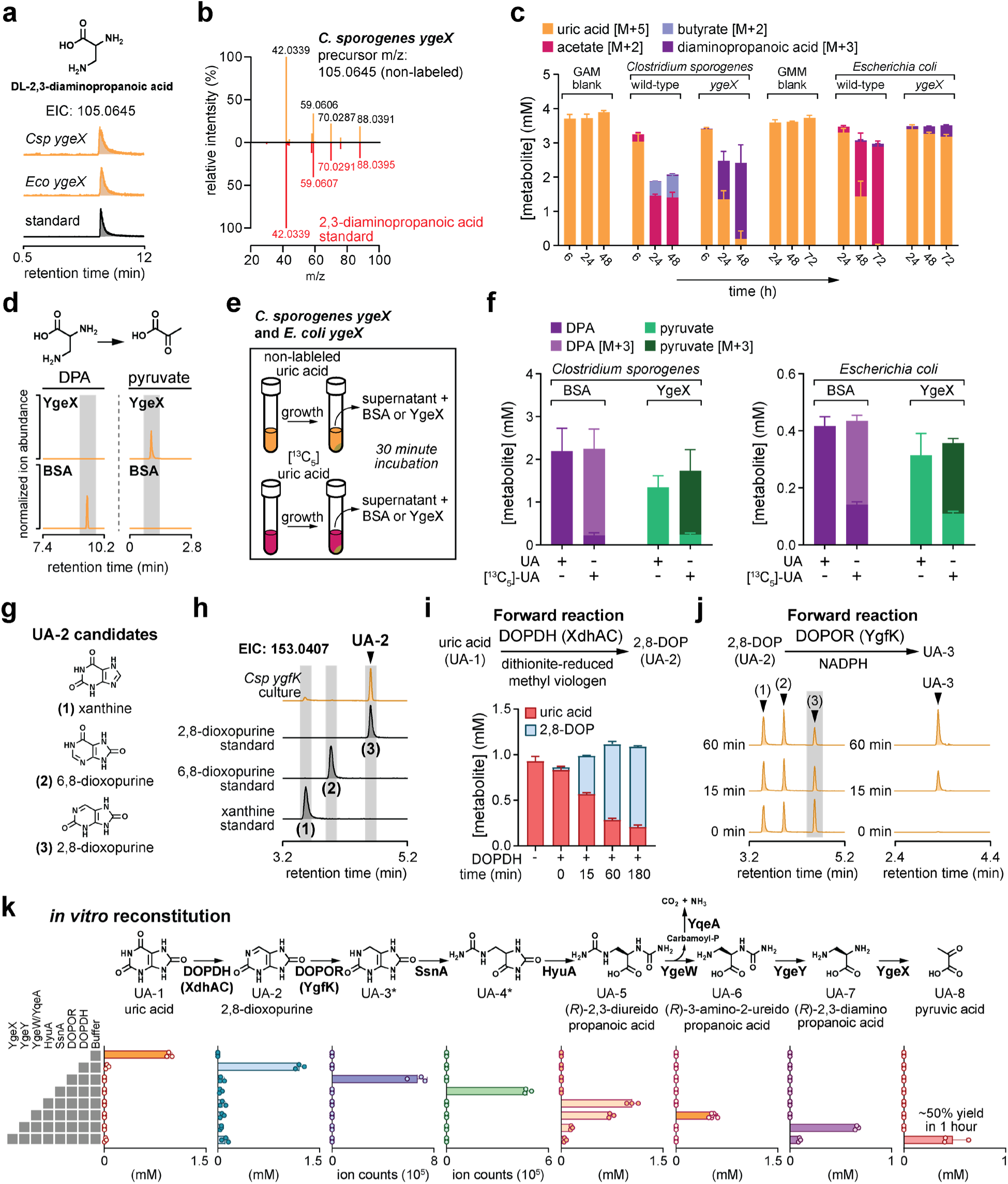
Complete *in vitro* reconstitution of the 2,8-dioxopurine pathway. (**A**) Extracted ion chromatograms from HILIC LC-MS for 2,3-diaminopropanoic acid standard and the molecular feature identified in *C. sporogenes* and *E. coli ygeX* supernatants. (**B**) Comparison of mass fragmentation spectra between *C. sporogenes ygeX* supernatants and an authentic 2,3-diaminopropanoic acid standard. (**C**) Labeled substrates and products detected in cell-free culture supernatants of *C. sporogenes* and *E. coli* wild-type and *ygeX* mutants using Poroshell positive (uric acid), DNS (DPA), or NPH (acetate and butyrate) LC-MS. (**D**) Extracted ion chromatograms for 2,3-diaminopropanoic acid and pyruvate after incubation of 2,3-diaminopropanoic acid with BSA (control) or recombinant YgeX. Compounds were analyzed by DNS (DPA) or C18 (pyruvic acid) LC-MS. (**E**) Overview of enzyme conversion assays in culture supernatants. (**F**) Concentrations of labeled or non-labeled 2,3-diaminopropanoic acid and pyruvate corresponding to experimental overview in **E**. Compounds were analyzed by DNS (DPA) or C18 negative (pyruvic acid) LC-MS. (**G**) Structures of candidates for the second intermediate in the pathway (UA-2) including xanthine and its two isomers. (**H**) Comparison of extracted ion chromatograms for *C. sporogenes ygfK* culture supernatants and authentic standards for the three xanthine isomers using HILIC LC-MS. (**I**) Conversion of uric acid to 2,8-dioxopurine using purified *C. sporogenes* DOPDH (XdhAC), analyzed by Poroshell positive LC-MS. (**J**) Extracted ion chromatograms for UA-2 and UA-3 after incubation of recombinant *E. coli* DOPOR (YgfK) with a xanthine isomer mixture using HILIC LC-MS. (**K**) *In vitro* reconstitution of the uric acid pathway. Each enzyme was added sequentially, and intermediates were quantified by Poroshell positive (uric acid and 2,8-dioxopurine), HILIC (UA-3, UA-4, UA-5, and UA-6), DNS (DPA), or Poroshell negative (pyruvic acid) LC-MS after 1 hour. Structures for the intermediates are supported by comparison of retention times, accurate m/z, and mass fragmentation patterns with authentic standards, except for UA-3 and UA-4 which are provisional (see **Supplemental Text** for details). For **A**, **D**, **H**, and **J**, three replicates were performed, and representative data are shown. For **C**, **F**, **I**, and **K**, data represent means ± standard deviations from three replicates.

## 2,8-dioxopurine, not xanthine, is the entry point into the uric acid pathway

Our stable isotope tracing studies enabled us to infer the position of each gene in the uric acid pathway (**Figure 1c**). Based on the pattern of intermediates that accumulated in the *C. sporogenes* and *E. coli* mutants, we predicted that *xdhAC* encodes the enzyme responsible for the first step in the pathway. *C. sporogenes* XdhAC possesses iron-sulfur cluster binding domains and molybdopterin co-factor (MoCo) binding domains, features characteristic of xanthine dehydrogenases (**Extended Data Figure 6a**). Given these similarities, we initially thought that XdhAC would convert uric acid to xanthine. To test this, we analyzed cultures of the *C. sporogenes ygfK* mutant, which is blocked in the next step of the pathway. Surprisingly, we detected only trace levels of xanthine but observed prominent accumulation of an unknown compound with the same monoisotopic mass as xanthine, but a different retention time (**Extended Data Figure 6b**). Suspecting this metabolite to be an isomer of xanthine, we synthesized 2,8-dioxopurine and 6,8-dioxopurine, as commercial standards were unavailable (**Figure 2g**). LC-MS analyses confirmed that the unknown compound in the *C. sporogenes ygfK* cultures was 2,8-dioxopurine (**Figure 2h** and **Extended Data Figure 6c**).

To test whether *C. sporogenes* XdhAC produces 2,8-dioxopurine, we sought to purify the native enzyme. Because molybdenum cofactor (MoCo)-dependent enzymes require complex cofactor biosynthesis and assembly, overexpressing all necessary components for MoCo synthesis would be challenging. Instead, we opted to purify XdhAC natively by using CRISPR-based genome editing to introduce a C-terminal His-tag in-frame with the *C. sporogenes xdhAC* coding sequence (**Extended Data Figure 6d**). Importantly, this modification did not block anaerobic uric acid consumption in the engineered strain (**Extended Data Figure 6e**). We then purified XdhAC from *C. sporogenes* cells grown with uric acid (**Extended Data Figure 6f**) and assessed its activity using methylviologen as an artificial electron donor/acceptor. XdhAC catalyzed the conversion of uric acid to 2,8-dioxopurine (forward reaction, **Figure 2i**) and also mediated the reverse reaction, converting 2,8-dioxopurine back to uric acid (**Extended Data Figure 6g**). Similar to other Clostridial molybdenum hydroxylases^23^, XdhAC did not use NADH or NADPH as electron donors (**Extended Data Figure 6h**). Although the physiological electron donor/acceptor remains unknown, these results establish XdhAC as a 2,8-<u>dio</u>xo<u>p</u>urine <u>d</u>e<u>h</u>ydrogenase (DOPDH) that catalyzes the 6-dehydroxylation of uric acid.

With 2,8-dioxopurine identified as the key intermediate, we next investigated the function of YgfK, the enzyme responsible for its downstream metabolism. YgfK contains a 4Fe-4S cluster binding domain, a pyridine nucleotide-disulfide oxidoreductase domain, and two large N- and C-terminal domains of unknown function (**Extended Data Figure 6i**). To obtain functional recombinant YgfK, we co-expressed it with the *E. coli* iron-sulfur repair system (*sufABCDSE*)^24^ and purified it under anaerobic conditions (**Extended Data Figure 6i**). From a single iMAC step, we obtained reasonably pure YgfK (**Extended Data Figure 6j**). Initial scouting experiments showed that NADPH, but not NADH, was required for YgfK activity (**Extended Data Figure 6k**). Despite its annotation as a selenate reductase^25^, selenate was neither required nor did it enhance the activity (**Extended Data Figure 6k-l**). When incubated with a mixture of the three xanthine isomers, YgfK selectively converted 2,8-dioxopurine to UA-3 (**Figure 2j**). Therefore, YgfK is an NADPH:2,8-<u>dio</u>xo<u>p</u>urine <u>o</u>xidoreductase which we name DOPOR.

Together, our findings reveal a previously unknown metabolic route for anaerobic uric acid degradation. The pathway begins with DOPDH (encoded by *xdhAC*) which catalyzes the conversion of uric acid to 2,8-dioxopurine, followed by DOPOR (encoded by *ygfK*) which reduces 2,8-dioxopurine to UA-3. These discoveries redefine the entry point into gut bacterial purine metabolism, demonstrating that 2,8-dioxopurine, rather than xanthine, is the key intermediate in gut bacterial uric acid degradation.

## *In vitro* reconstitution of the uric acid pathway

With knowledge of the beginnings and ends of the pathway, we next aimed to complete the pathway through stepwise addition of each enzyme. Using a combination of chemical synthesis, isotope tracing, mass spectrometry, and *in vitro* enzyme assays we assigned structures for intermediates in the pathway (**Figure 2k** and **Supplementary Text**). We note that structural assignments for UA-3, which is unstable, and UA-4, which we could not synthesize, are provisional (see **Supplementary Text**). Sequential addition of each enzyme yielded the next intermediate in the pathway and after all enzymes were added, uric acid was converted to pyruvate with a yield of ∼50% in 1 hour (**Figure 2k**). Our findings indicate that SsnA opens the six-membered ring, yielding a ureido-substituted hydantoin (UA-4). Next, HyuA, a cyclic amidohydrolase with activity towards hydantoins^26^, opens the five-membered ring yielding (*R*)-2,3-diureidopropanoic acid (UA-5). YgeW is a carbamoyltransferase^27^ which then cleaves the 3-ureido group, yielding (*R*)-3-amino-2-ureidopropanoic acid (UA-6). Next, YgeY, a putative amidase, hydrolyzes the 2-ureido group yielding 2,3-diaminopropanoic acid which is converted to pyruvate by YgeX. This pathway expands the scope of biochemical pathways for purine metabolism, introducing five intermediates (UA-2 through UA-6) not previously known in biology.

## Purine metabolism is linked to ATP formation through YgeW and YqeA

While reconstructing the uric acid pathway, we identified a potential link to ATP formation, which could provide a physiological advantage to bacteria that possess this pathway. YgeW is a carbamoyl transferase that cleaves UA-5 to UA-6, likely generating carbamoyl phosphate in the process. In our human gut bacteria strain library, we observed that many uric acid gene clusters also include a carbamate kinase (annotated as YqeA in *E. coli*), often located immediately downstream of YgeW (**Extended Data Figure 7a**). In most strains (33/39) carbamate kinase was found within the uric acid gene cluster (**Extended Data Figure 7b**). However, *C. sporogenes* ATCC 15579, *Collinsella aerofaciens* ATCC 25986, and *Collinsella sp.* 4_8_47FAA had carbamate kinase in a different region on the chromosome, while three additional strains lacked both carbamate kinase and YgeW (*Fusobacterium ulcerans* 12-1B, *Clostridioides difficile* ATCC BAA-1801 and *C. difficile* 630) (**Extended Data Figure 7b**). This suggested that in most organisms, coordinated action of YgeW and carbamate kinase could generate ATP (**Figure 3a**). To test this, we incubated recombinant *E. coli* YgeW and YqeA either individually or together with the substrate for YgeW (UA-5) and with magnesium and ADP, known requirements for carbamate kinases. Incubation of enzymatically synthesized UA-5 with YgeW and YqeA resulted in substantial ATP production, while only minimal ATP was generated when each enzyme was tested individually or in the buffer control (**Figure 3b**). ATP formation from YgeW and YqeA occurred concomitantly with conversion of UA-5 to UA-6 (**Figure 3c**). Therefore, our results show that uric acid degradation is linked to ATP formation through substrate level phosphorylation involving the concerted action of a carbamoyl transferase and carbamate kinase. This metabolic strategy is reminiscent of the arginine deiminase pathway where ornithine carbamoyltransferase and carbamate kinase generate _ATP28,29._

**Figure 3.**
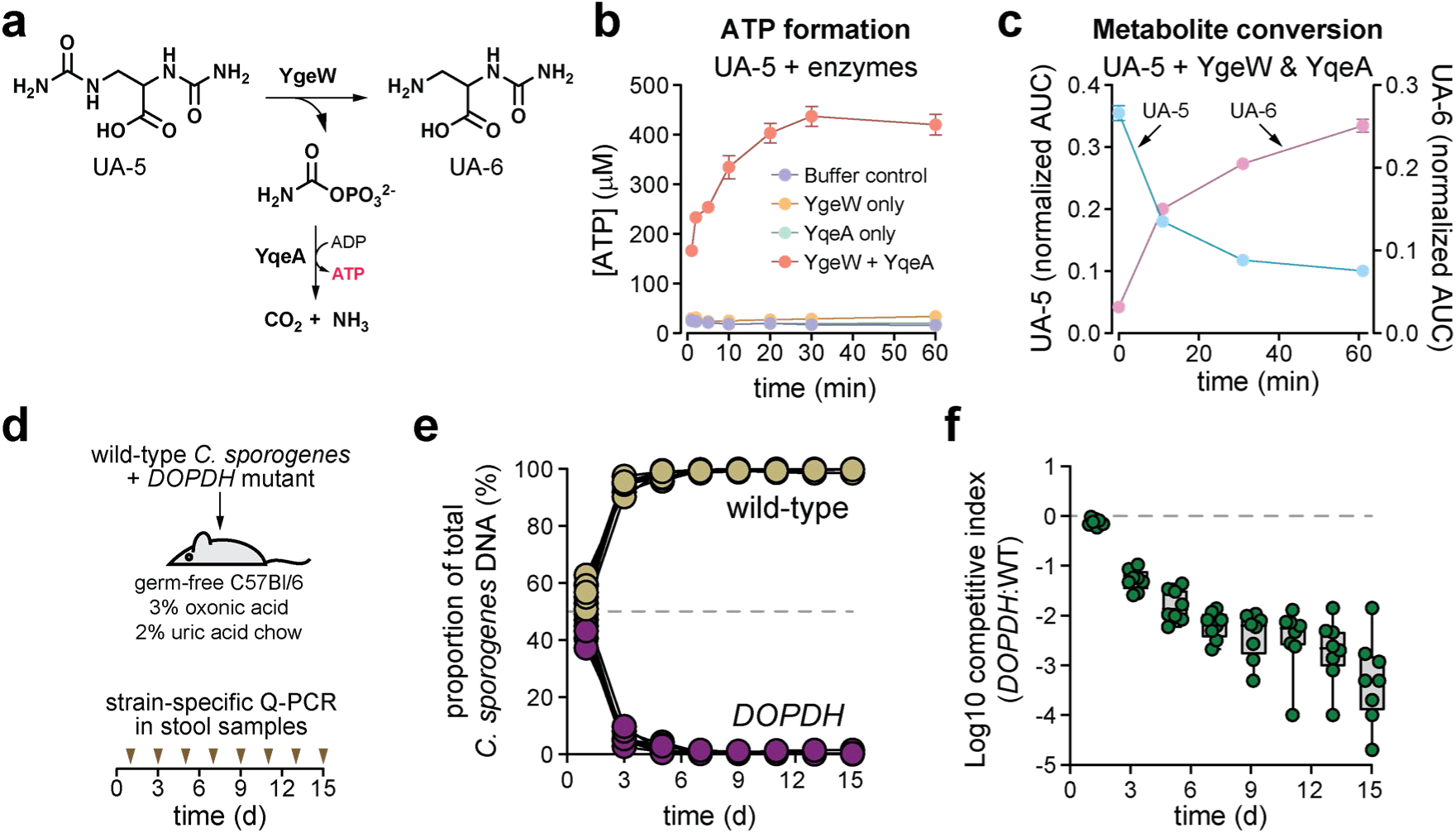
The 2,8-dioxopurine generates ATP and enhances microbial fitness *in vivo*. (**A**) Model for ATP formation during conversion of 2,3-diureidopropanoate (UA-5) to 3-amino-2-ureidopropanoate (UA-6) involving the concerted action of YgeW and carbamate kinase (YqeA). (**B**) ATP production in enzyme reactions using UA-5 as substrate. (**C**) Conversion of UA-5 to UA-6 in the YqeA + YqeW enzyme reaction from **B** analyzed by HILIC LC-MS. AUC, area under the curve. (**D**) Overview of *in vivo* competition experiment comparing wild-type and *DOPDH* (*xdhAC*) mutant strains of *C. sporogenes*. Equal amounts of both strains were administered to mice fed a hyperuricemia-inducing chow and strain abundance in stool was quantified by quantitative PCR. (**E**) Co-challenge data showing the proportion of wild-type and *DOPDH* strains in stool samples of mice after colonization. (**F**) Competitive index calculated from co-challenge data in **E**. For **B** and **C**, data represent means ± standard deviations from three replicates. For **E**, individual data points for n = 8 mice are shown. For **F**, boxes extend from the 25th to the 75th percentiles with lines at the median, and whiskers go down to the lowest value and up to the highest value for n = 8 mice per group; male germ-free mice (*M. musculus* (B6NTac)) aged 6–8 weeks were used for the experiments.

## The 2,8-dioxopurine pathway confers a bacterial fitness advantage in the gut

To directly test whether uric acid metabolism provides a competitive advantage to bacteria *in vivo*, we performed a competition experiment in hyperuricemic germ-free mice. Equal amounts of wild-type and *DOPDH* (*xdhAC*) mutant *C. sporogenes* were orally administered, and the relative abundances of the two strains were quantified in stool samples using quantitative PCR (**Figure 3d**). The *DOPDH* mutant was rapidly outcompeted by the wild-type strain, with a significant reduction in its relative abundance observed as early as 24 hours post-inoculation (**Figure 3e**). By day 7, the *DOPDH* mutant was barely detectable, while the wild-type strain remained stably colonized (**Figure 3e**). These findings demonstrate that the ability to degrade uric acid via the 2,8-dioxopurine pathway confers a substantial competitive advantage to gut bacteria (**Figure 3f**).

## Anaerobic bacterial purine metabolism requires selenium-dependent molybdenum hydroxylases

Selenium is essential for certain bacterial redox enzymes, functioning in selenocysteine incorporation, tRNA modification, and molybdenum hydroxylase activation. The latter Se-cofactor trait is thought to involve the accessory proteins YqeB and YqeC^30–33^ which may incorporate selenium into enzymes such nicotinic acid hydroxylase^34,35^, xanthine dehydrogenase^23,36^, and purine hydroxylase^23,37^. All three of these selenium utilization traits are thought to require the activity of selenophosphate synthase (SelD) (**Figure 4a**).

**Figure 4.**
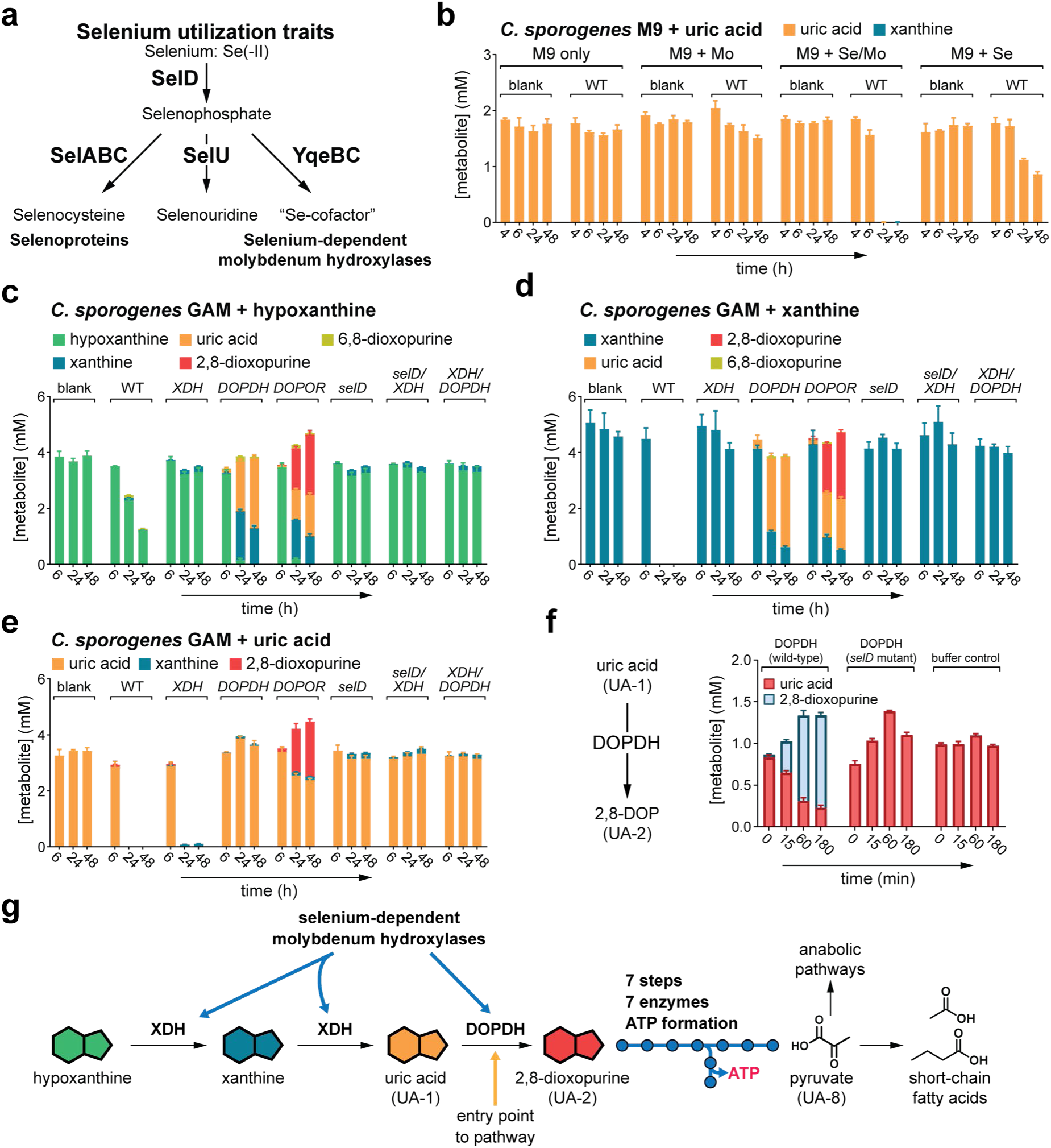
Anaerobic purine degradation represents a widely distributed third selenium utilization trait in bacteria. (**A**) Overview of selenium utilization traits in bacteria. (**B**) Uric acid consumption by *C. sporogenes* wild-type during growth in defined M9 media with or without the trace elements, molybdenum (Mo; sodium molybdate) and selenium (Se; sodium selenite). (**C-E**) Hypoxanthine, xanthine, and uric acid consumption by wild-type and mutant strains of *C. sporogenes*. (**F**) Enzyme activity of 2,8-dioxopurine dehydrogenase (DOPDH; encoded by the *xdhAC* gene) purified from either wild-type or *selD* mutant in the forward reaction, catalyzing the reduction of uric acid to 2,8-dioxopurine. (**G**) Overview of the purine metabolism pathway in gut bacteria. XDH and DOPDH are selenium-dependent molybdenum hydroxylases. The pathway generates ATP and culminates in pyruvate production, which can either be used for anabolic pathways or further catabolized to short-chain fatty acids. For **B-F**, metabolites were analyzed by Poroshell positive LC-MS and data represent means ± standard deviations from three replicates.

To explore the link between selenium metabolism and the 2,8-dioxopurine pathway, we analyzed selenium utilization traits in our gut bacterial strain library. All strains with the uric acid degradation gene cluster contained both YqeBC (Se-cofactor trait) and SelD (selenophosphate synthase), suggesting selenium dependence of the 2,8-dioxopurine pathway (**Extended Data Figure 8**). During growth in defined medium, both selenium and molybdenum were required for complete uric acid degradation by wild-type *C. sporogenes* (**Figure 4b**) and production of maximum levels of the SCFA, acetate (**Extended Data Figure 9a**). Residual uric acid degradation occurred with selenium alone, suggesting trace molybdenum contamination (**Figure 4b** and **Extended Data Figure 9a**).

The likely candidates for selenium-dependent enzymes include the molybdenum hydroxylases, DOPDH and xanthine dehydrogenase (XDH) which are both present in the *C. sporogenes* genome (**Extended Data Figure 9b**). We generated the single mutants, *XDH* and *selD*, and the double mutants *XDH/selD* and *XDH/DOPDH* in *C. sporogenes*. We then cultured these strains along with the wild-type, *DOPDH* and *DOPOR* single mutants in rich medium supplemented with either hypoxanthine, xanthine, or uric acid and quantified purines by LC-MS. Mutant analyses confirmed XDH as a xanthine dehydrogenase, its mutation blocking hypoxanthine and xanthine conversion to uric acid (**Figure 4c-d**), while DOPDH was blocked at the level of uric acid conversion to 2,8-dioxopurine (**Figure 4c-e**). Stable isotope tracing with [^13^C_5_]-labeled hypoxanthine extended these findings, showing that unlike wild-type, the *XDH* mutant did not produce labeled short-chain fatty acids (**Extended Data Figure 9c-d**).

The *selD* mutant displayed a metabolic profile resembling selenium-deficient conditions with disrupted Stickland metabolism (**Extended Data Figure 9e-f** and **Supplementary Table 7**) and defective purine degradation phenocopying the *XDH/DOPDH* double mutant (**Figure 4c-e**). This indicates that both molybdopterin-dependent enzymes (XDH and DOPDH) in *C. sporogenes* require selenophosphate synthase. To confirm that DOPDH requires selenophosphate synthase for its activity, we created a C-terminal his-tagged DOPDH strain in a *selD* mutant background and purified the enzyme natively from *C. sporogenes* (**Extended Data Figure 9g**). Western blots using an anti-His-tag antibody confirmed the presence of intact DOPDH in both wild-type and *selD* backgrounds (**Extended Data Figure 9h**). DOPDH purified from the wild-type background showed robust activity in the forward (**Figure 4f**) and reverse (**Extended Data Figure 9i**) directions. However, DOPDH purified from the *selD* mutant strain had no measurable activity (**Figure 4f** and **Extended Data Figure 9i**).

Our results demonstrate that the 2,8-dioxopurine pathway in *C. sporogenes* is selenium-dependent, requiring selenophosphate synthase (SelD) for XDH and DOPDH enzymes (**Figure 4g**). These findings are corroborated by a very recent paper by Johnstone and Self, which showed that purine-dependent growth in *Clostridioides difficile* requires SelD and the Se-cofactor marker protein, YqeB^30^.

## Functional differences between bacterial DOPDH and mammalian XOR

Unlike bovine XOR, which converts xanthine and 6,8-dioxopurine to uric acid, DOPDH selectively interconverts uric acid and 2,8-dioxopurine (**Extended Data Figure 10a-d**). DOPDH also differs from XOR by lacking NAD+/NADH dependence and functioning exclusively under anaerobic conditions (**Extended Data Figure 10a-e**). Phylogenetic analysis showed that DOPDH forms a unique cluster, most closely related to selenium-dependent nicotinate dehydrogenases (NDH; **Extended Data Figure 10f**). Despite low sequence identity, and the lack of a FAD binding domain in DOPDH, the molybdenum cofactor-binding domain of DOPDH shares strong structural conservation with that of bovine XOR (**Extended Data Figures 10g-h**). Amino acid sequence alignments revealed differences in active site residues (F1009V, A1078T/S) that may underlie its selective uric acid to 2,8-dioxopurine conversion (**Extended Data Figure 10h-j**). Similar to previous structural studies on selenium-dependent NDH^38^, we did not observe obvious active site residue differences that could explain the selenium vs. sulfur dependence of DOPDH vs. bovine XOR.

## Genes for uric acid degradation are widely distributed across host-associated microbiomes

To assess the global distribution of uric acid-degrading genes, we analyzed the Earth’s Microbiomes genomic catalog, comprising 52,515 metagenome-assembled genomes (MAGs) from 10,450 taxa (**Figure 5a**)^39^. We identified 2,102 MAGs from 1,350 bacterial taxa harboring the uric acid degradation gene cluster, with a strong enrichment in members of the Bacillota phylum (**Figure 5b**; **Supplementary Tables 8-9**).

**Figure 5.**
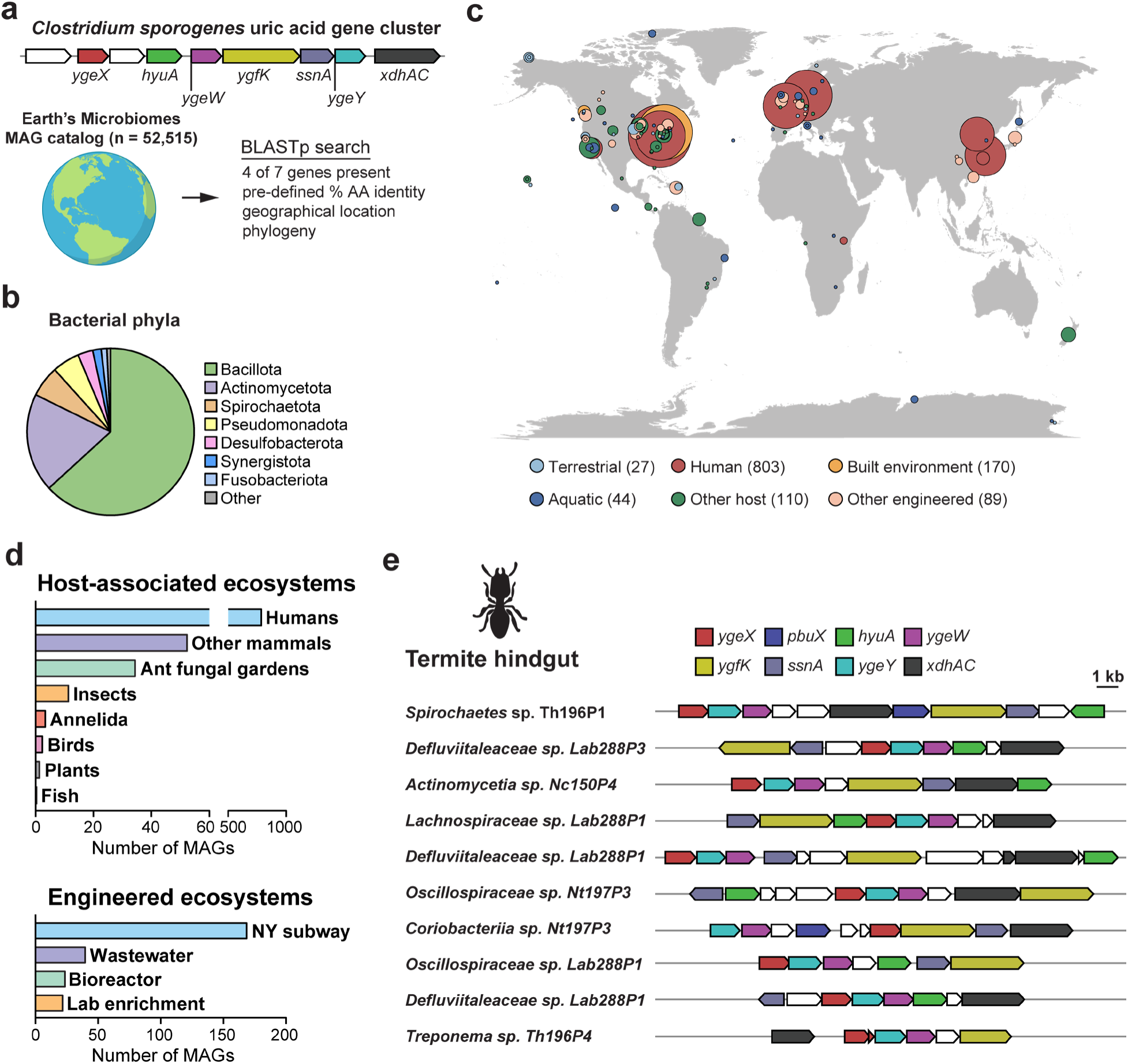
Distribution of uric acid genes across Earth’s microbiomes. (**A**) Uric acid genes from *C. sporogenes* were used as inputs for a BLASTp search of the Earth’s microbiomes metagenome assembled genome (MAG) catalog. (**B**) Phylum-level distribution of uric acid genes in Earth’s microbiomes. (**C**) Distribution of uric acid genes across the planet, color coded by habitat. (**D**) Number of MAGs containing the uric acid genes in different host-associated and engineered ecosystems. (**E**) The genomic context for uric acid genes from representative MAGs in termite metagenomes.

Uric acid-degrading genes were enriched in host-associated ecosystems (**Figure 5c-d**) including the gut of diverse mammals, birds, insects, and worms, including chickens and termites – two model organisms that excrete uric acid as a water-sparing nitrogenous waste (i.e., uricotelic organisms). A deeper analysis of MAG collections from chickens^40^ and termites^41^ revealed numerous bacterial taxa harboring the uric acid genes (**Supplementary Tables 10-11**) arranged in characteristic metabolic gene clusters (**Figure 5e**). These findings suggest that the 2,8-dioxopurine pathway has been selected for in the animal gut and may play a role in host-microbe symbioses.

## Discussion

Our discovery of the 2,8-dioxopurine pathway expands the known landscape of purine metabolism, revealing a previously unrecognized anaerobic route distinct from well-characterized pathways in aerobic bacteria^42,43^, yeast^44^, and mammals^5^. In humans and apes, purine catabolism progresses via xanthine oxidoreductase (XOR) to produce uric acid, which accumulates as the terminal product due to the loss of urate oxidase (uricase). In contrast, most other mammals express functional urate oxidase, which further converts uric acid to allantoin, a more soluble compound excreted in the urine.

The anaerobic 2,8-dioxopurine pathway identified in our study highlights a key contrast between aerobic and anaerobic purine metabolism. Unlike oxygen-dependent urate oxidase, anaerobes employ DOPDH, an unusual selenium-dependent molybdenum hydroxylase, to reduce uric acid to 2,8-dioxopurine – effectively running the XOR reaction in reverse. This reduction is further extended by DOPOR, the next enzyme in the pathway, resulting in a net four-electron reduction of uric acid. In anaerobic environments, where oxygen-dependent electron transport chains are absent, organisms must rely on alternative strategies to balance redox homeostasis. This reductive pathway likely serves as an electron sink, facilitating the regeneration of oxidized nicotinamides (NAD⁺/NADP⁺), which in turn helps drive energy-yielding oxidative reactions. By coupling purine metabolism to electron flow, anaerobes optimize both redox balance and ATP production, highlighting a key adaptation to life in oxygen-limited environments.

Why do anaerobes use 2,8-dioxopurine as an intermediate in the pathway instead of its isomer, xanthine? We propose two possible explanations: First, structural differences between 2,8-dioxopurine and xanthine may alter the resonance energy and ring stability, making 2,8-dioxopurine more reactive and thus more readily reduced and hydrolyzed by downstream reactions. This would be particularly important for anaerobes, who cannot harness the power of molecular oxygen for aromatic ring cleavage. Second, using 2,8-dioxopurine as an alternative intermediate may allow the anaerobic degradation pathway to remain metabolically insulated from purine salvage, preventing unwanted cross-talk between these pathways.

Our *in vivo* data in hyperuricemic gnotobiotic mice reveal an important fitness advantage conferred by this pathway. Wild-type bacteria harboring DOPDH rapidly outcompete a DOPDH-deficient strain demonstrating that the ability to metabolize uric acid plays a crucial role in bacterial colonization and persistence. The selective advantage conferred by purine metabolism underscores the broad impact this type of metabolism has on shaping microbial ecology within the gut.

This metabolic interaction represents a unique host-microbe symbiosis, wherein the host secretes uric acid into the intestines, providing a metabolic niche for bacteria capable of breaking it down. In return, bacteria enhance uric acid elimination, helping to regulate systemic uric acid levels in the host and mitigate hyperuricemia-associated pathology^2,3^. This symbiotic exchange highlights how microbial metabolism can directly influence host physiology, reinforcing the functional significance of gut bacteria in purine homeostasis^12^.

The wide distribution of the 2,8-dioxopurine pathway in bacteria colonizing the gastrointestinal tract of diverse animal species suggests that this host-microbe relationship has been conserved and refined over evolutionary time. The finding of these genes in the guts of insects, birds, and mammals implies that host-microbe purine interactions play an important role across a variety of different ecological contexts. In uricotelic organisms, such as birds and insects, the 2,8-dioxopurine pathway may improve the efficiency of water-sparing nitrogen excretion in two ways. First, the nitrogen may be held as a nutritional reserve in bacteria, being re-used in times of nutrient limitation through coprophagy or trophyllaxis (anal feeding)^45^. Second, bacterial production of SCFAs may provide the host with metabolic fuel from which the energetic cost of uric acid synthesis can be recovered^46^.

Beyond its ecological and evolutionary implications, our findings have therapeutic potential. Elevated serum urate (hyperuricemia) is the fundamental risk factor for all patients with gout, which has recently been estimated to affect ∼5% of US adults^9^. Strategies to enhance uric acid degradation by gut bacteria could provide a novel, safer alternative or complementary approach to currently prescribed urate-lowering therapies (ULTs)^10,12^. This could be especially important for individuals with severe chronic kidney disease who rely mainly on the gastrointestinal route for urate excretion and in whom ULT use is more complex, with some agents contraindicated^11^.

Our findings offer two significant translational insights: First, by establishing the biochemical basis for urate degradation, our results pave the way for engineering this pathway into probiotics that can be therapeutically administered. Second, since selenium and molybdenum are required for the 2,8-dioxopurine pathway, ensuring an adequate supply of these micronutrients to the gut microbiota may enhance intestinal urate elimination. Supporting this hypothesis, data from NHANES show that lower urinary molybdenum levels correlate with higher serum urate and increased gout incidence^47^. Additionally, a meta-analysis of 46 studies found that CKD patients on hemodialysis have significantly lower selenium levels^48^ and another study demonstrated a negative correlation between serum selenium and urate levels in CKD patients^49^.

Our discovery of the 2,8-dioxopurine pathway reveals a previously unknown anaerobic route of purine metabolism, with broad ecological, evolutionary, and clinical implications. This pathway not only defines a key metabolic distinction between aerobic and anaerobic life but also provides a potential target for microbiome-based interventions in managing hyperuricemia and gout.

## Methods

### Reagents used in this study

All chemicals and reagents used in this study were of the highest possible purity (**Supplementary Table 12**). Due to solubility issues, uric acid, [^13^C_5_]-uric acid, xanthine, 2,8-dioxopurine, hypoxanthine, and [^13^C_5_]-hypoxanthine were made fresh at 120 mM in 0.4 N sodium hydroxide (NaOH) prior to use. 6,8-dioxopurine was prepared at 40 mM in dimethyl sulfoxide (DMSO). The stock solution was used directly as LC-MS standard or sterile filtered and then diluted appropriately in anaerobic media for consumption assays.

### Bacterial strains and culture conditions

Bacterial strains used in this study are listed in **Supplementary Table 13**. All bacteria were cultured at 37 °C in a Coy type B anaerobic chamber using a gas mix containing 5% hydrogen, 10% carbon dioxide, and 85% nitrogen. An anaerobic gas infuser was used to maintain hydrogen levels of 3.3%. All media and plasticware were pre-reduced in the anaerobic chamber for at least 24 hours before use. For anaerobic uric acid consumption assays, *E. coli* was cultured in Gifu Anaerobic Medium, modified (GAM.M) and *C. sporogenes* was cultured in Gifu Anaerobic Medium (GAM) or M9 medium (**Supplementary Table 13**). Antibiotic concentrations in broth and agar media were used as follows: *E. coli*, chloramphenicol (25 μg/mL), carbenicillin (100 μg/mL); *C. sporogenes*, thiamphenicol (15 μg/mL); erythromycin (5 μg/mL).

### Culture conditions for in vitro growth consumption assays

All the cultures were performed under anaerobic conditions. *E. coli* strains were streaked on GAM.M plates and cultured in GAM.M broth, while *C. sporogenes* strains were streaked on Reinforced Clostridial Medium (RCM) plates and cultured in GAM broth. Individual colonies were picked and inoculated in rich media for 16-18 h. Subsequently, the cultures were diluted 50-fold in media containing 4 mM of purine and cultured at 37 °C for 48 to 72 h. At designated time points, samples were harvested by centrifugation (5,000 x *g*, room temperature, 5 min) and supernatants were collected and divided into two plates: one for SCFA measurement and the other for uric acid measurement. Both plates were stored at -80°C until analysis by LC-MS.

### Stable isotope tracing with [^13^C_5_]-uric acid or [^13^C_5_]-hypoxanthine

The culture conditions were the same as described above. To correct for natural isotope abundance, we cultured organisms with labeled and unlabeled purines (uric acid or hypoxanthine). After quantifying SCFA isotopologs, we subtracted concentrations of isotopologs in cultures with unlabeled purine from those cultured with labeled purine as previously described ^3^.

### Sample preparation for analysis of purines, pathway intermediates, pyruvate and SCFA by liquid chromatography – mass spectrometry (LC-MS)

Uric acid, xanthine, 2,8-dioxopurine, and hypoxanthine were freshly prepared at 120 mM in 0.4 N NaOH each time. 6,8-dioxopurine was dissolved at 40 mM in LC-MS grade DMSO and stored at -20°C. Pyruvate was freshly made at 100 mM in LC-MS grade water. These stock solutions were subsequently diluted in 100 mM potassium phosphate buffer, pH 7.5, to make calibration standards for LC-MS assays. To account for potential matrix effects, an equal portion of growth medium was added into the standard curves for *in vitro* culture samples. Calibrants were treated the same as samples during LC-MS preparation.

#### Metabolite extraction from samples

Culture supernatants or enzyme assay aliquots were first mixed with internal standard (ISTD) in a V-bottom, polypropylene 96-well plate, and then extracted by mixing with extraction solution (75% acetonitrile/25% methanol) at 1:3 ratio. The plate was covered with a lid and centrifuged at 5,000 x *g* for 10 min at room temperature to separate the supernatant.

The collected supernatant was preserved at -80 °C for LC-MS analysis. For measurement of UA intermediates (UA-3, UA-4, UA-5, UA-6, and 2,3-diaminopropanoic acid) and pyruvate, the extracted supernatants were used directly. To measure uric acid, xanthine isomers, and hypoxanthine, the supernatants were diluted in NH_4_OH (final ∼3 mM) before LC-MS analysis.

Additionally, the extracted supernatants were also used for 3-nitrophenylhydrazine derivatization (for SCFA measurement) or dansylation derivatization (for 2,3-diaminopropanoic acid measurement).

#### Dansylation (DNS) derivatization protocol

This derivatization method targets compounds containing a free amine. Extracted samples were mixed with dansyl chloride (50 mM in acetonitrile) and sodium carbonate/bicarbonate (100 mM, pH 9.8) at 1:1:1 ratio. The plate was covered with a sealing mat and incubated at 25 °C, 300 rpm in a ThermoMixer for 60 min. Ammonium hydroxide (final 0.1% v/v) was added to the plate and incubated at 25 °C for 5 min to quench the reaction. Then the mixture was diluted in 40% acetonitrile/0.01% formic acid before LC-MS.

#### 3-Nitrophenylhydrazine (NPH) derivatization protocol

This derivatization method targets compounds containing a free carboxylic acid. Extracted samples were diluted in 50% acetonitrile and then mixed with 3-nitrophenylhydrazine (200 mM in 80% acetonitrile) and *N*-(3-dimethylaminopropyl)-*N*’-ethylcarbodiimide (120 mM in 6% pyridine) at 1:1:1 ratio. The plate was sealed with a sealing mat and incubated at 40 °C, 300 rpm in a ThermoMixer for 60 min to derivatize the carboxylate containing compounds. The reaction mixture was quenched with 0.02% formic acid in 20% acetonitrile/water before LC-MS analysis.

#### Quantification of metabolites by liquid chromatography-mass spectrometry (LC-MS)

During this study, six different LC-MS conditions were used (C18 negative underivatized, Poroshell positive underivatized, Poroshell negative underivatized, HILIC positive underivatized, 3-nitrophenylhydrazine (NPH) derivatized C18 negative, and dansyl chloride (DNS) derivatized C18 positive). An overview of the general method is provided here and the specific instrument parameters for the different analytical methods are provided in **Supplementary Table 14**. Samples were injected via refrigerated autosampler into mobile phase and chromatographically separated by an Agilent 1290 Infinity II UPLC and detected using an Agilent 6545XT Q-TOF equipped with a dual jet stream electrospray ionization source operating under extended dynamic range (EDR 1700 m/z). MS1 spectra were collected in centroid mode, and peak assignments in samples were made based on comparisons of retention times and accurate masses from authentic standards (when available) using MassHunter Quantitative Analysis v.10.0 software from Agilent Technologies. Compounds were quantified from calibration curves constructed with authentic standards (when available) using isotope-dilution mass spectrometry with appropriate internal standards (**Supplementary Table 15**).

#### Comparison of mass fragmentation patterns for pathway intermediates and standards

Compounds were separated chromatographically using our standard HILIC positive method (**Supplementary Table 14**). For samples and authentic standards, we used targeted MS/MS, with pre-defined precursor m/z for each intermediate and with retention time window of 0.3 min, narrow isolation width, fixed collision energy of 20 eV, and acquisition time of 400 milliseconds per spectrum. Spectra were manually inspected using Agilent MassHunter Qualitative Analysis (v.10.0.10305.0) and the cleanest spectra centered closest to the maximum peak area for the precursor compound were exported as .txt files. After normalizing spectra to the maximum fragment ion intensity, spectra were used as queries for similarity searches in the Mass Databank of North America database.

#### CRISPR and ClosTron genetics in Clostridium sporogenes

Disruption of the *Clostridium sporogenes selD* gene was achieved using the Intron targeting and design tool available on the ClosTron website (http://www.clostron.com/clostron2.php) with the Perutka algorithm^50^. The intron within the pMTL007C-E2 plasmid was retargeted to the site listed in **Supplementary Table 16** and the targeting sequences were synthesized as gBlocks by IDT. Retargeted plasmids were made by digesting the pMTL007C-E2-CLOSPO_00316-736s^51^ plasmid with BsrGI and HindIII, followed by gel-purification of the plasmid backbone and Gibson assembly with retargeted intron gBlock fragments using NEBuilder HiFi DNA Assembly Master Mix.

The resulting Gibson assemblies were transformed into *E. coli* Stbl4 by electroporation, selected on LB-chloramphenicol plates and miniprepped plasmids were sequenced to confirm the correct retargeted sequence. Subsequently, intron retargeted plasmids were transformed into *E. coli* CA434 and then conjugated into *C. sporogenes* as described previously^52^. After picking and culturing transconjugants in antibiotic-free media, erythromycin resistant mutants were obtained and verified by PCR and sequencing.

Initial experiments with the *Clostridium sporogenes ygeW* or *ygeX* ClosTron mutants suggested polar effects on downstream genes, therefore new mutants were generated by CRISPR-deletion using the RiboCas system^53^. The CRISPR target site was designed by Geneious Prime, and the sgRNA was synthesized as a gBlock by IDT. Homologous DNA repair templates were amplified by PCR from wild-type *C. sporogenes* genomic DNA using primers listed in **Supplementary Table 16**.

The sgRNA and repair template fragments were cloned into pRECas1 plasmid by Gibson assembly (NEBuilder HiFi DNA Assembly Master Mix). The Gibson products were transformed into *E. coli* Stbl4 by electroporation and selected on LB-chloramphenicol plates and miniprepped plasmids were sequenced to confirm the correct retargeted sequence. The correct CRISPR plasmids were transformed into *E. coli* CA434 and subsequently conjugated into *C. sporogenes*. The expression of *cas9* was induced by 5 mM theophylline on thiamphenicol plates. Mutants were validated by PCR and sequencing.

To enable native purification of the *C. sporogenes* XdhAC protein by immobilized metal affinity chromatography, a thrombin site followed by a 10X His-tag fragment (using the same sequence as pET-52b (+)) was inserted to the *xdhAC* gene (CLOSPO_02131) at the 3′ end of the gene just preceding the stop codon, using the RiboCas system. Additionally, a silent mutation was introduced in the DNA repair template to eliminate the PAM sequence such that the plasmid would not be targeted by Cas9. The resulting mutants, designated as *Csp 02131xdhAC-CHis,* underwent verification through PCR and sequencing. Furthermore, LC-MS analysis was conducted to ensure that uric acid consumption remained unaffected by the His-tag insertion.

### ATP measurements in enzyme assays

UA-5 intermediates were generated anaerobically through enzyme synthesis under the following conditions: purified YgfK (final 0.2 mg/mL), SsnA (final 0.02 mg/mL) and HyuA (final 0.02 mg/mL) were combined with 1 mM of 2,8-dioxopurine, 4 mM of NADPH, 2 mM of sodium selenate, and 0.04 mM of MnCl_2_ in 100 mM potassium phosphate (KPi) buffer, pH 7.5. The reaction mixture was incubated at 37 °C in an anaerobic chamber for 3 h. After incubation, the reaction mixture was filtered through a 3 KDa filter to remove all enzymes, and the flow-through was stored at -80 °C. The depletion of 2,8-dioxopurine and the accumulation of UA-5 in the flow-through were confirmed by LC-MS analysis. The product was termed UA-5 substrate. Selenate was initially included in the reaction due to uncertainty regarding its necessity for YgfK. However, our subsequent results indicated that it is not required for the reaction.

The ATP assay was performed under aerobic conditions. During ATP assay, UA-5 substrate was combined with MgSO_4_ (final 2 mM) and ADP (final 1 mM), with or without YgeW (final 1 mg/mL) and/or YqeA (final 0.05 mg/mL) in a 96-well plate kept on ice. The reaction mixture was then incubated in a pre-warmed ThermoMixer at 37 °C for 1 h. At specified time intervals (0 min, 2 min, 5 min, 10 min, 20 min, 30 min, and 60 min), 4 μL aliquots were taken and immediately mixed with 180 μL of DMSO and 16 μL of 100 mM KPi buffer, pH 7.5 to quench the reaction, and saved for ATP measurement.

In addition, at time intervals (0 min, 11 min, 31 min and 61 min), 12 μL of reaction aliquots were withdrawn and mixed with 18 μL of ISTD and 90 μL of extraction solution (75% acetonitrile/25% methanol) for LC-MS analysis.

ATP levels were quantified using a luminescence-based ATP determination kit (Invitrogen, A22066) with a Synergy H1M multimode plate reader from BioTek (Winooski, VT). The ATP concentrations were determined by interpolation using GraphPad Prism based on a calibration curve constructed with known concentrations of ATP.

### Identification of pathway intermediates in C. sporogenes mutants using stable isotope tracing and shotgun metabolomics

Wild-type and mutant strains (*ygfK*, *ssnA*, *hyuA*, *ygeW*, *ygeY*, and *ygeX*) of *C. sporogenes* were cultured in triplicate in GAM broth supplemented with either unlabeled or [^13^C_5_]-labeled uric acid. Cell-free control samples (blanks) were also included. Aliquots of culture supernatants were collected at 7, 24, and 48 h and metabolites were extracted as described above. Extracted metabolites in culture supernatants were separated using a HILIC column and full spectrum MS1 data were collected using an Agilent 6545XT Q-TOF (conditions provided in **Supplementary Table 14**). Each mutant was compared with the wild-type and blanks in cultures with either unlabeled or [^13^C_5_]-labeled uric acid using MS-DIAL^54^. Data files were converted to .abf format using Reifycs Analysis Base File convertor (ver. 1.3.7815) and were imported into MS-DIAL (ver.4.9.221218) and two group comparisons were made (for example, wild-type vs. *ygfK* in unlabeled uric acid) including sample blanks for feature subtraction. Default MS-DIAL settings were used with the following exceptions: minimum peak height, 3000; remove features based on blank information. After peak processing the raw data matrix (area) was exported with filtering on blank samples. This was done for all wild-type vs. individual mutant comparisons with either unlabeled or labeled uric acid. We then developed a custom workflow for identification of isotope paired intermediates enriched in mutant vs. wild-type cultures (implementation available on GitHub; https://github.com/DoddLab/DoddLabTracer). Briefly, 1) the enriched features during growth with unlabeled uric acid (mutant vs. WT, marked as “enriched-unlabeled features” below) and [^13^C_5_]-labeled uric acid (mutant vs. WT, marked as “enriched-labeled features” below) were retrieved separately from the MS-DIAL exported peak tables. The Student T-test and fold-change were used to retrieve these enriched features (P-value ≤ 0.05, FDR corrected; fold-change ≥ 10); 2) Enriched-unlabeled features were then collapsed by detecting adducts, neutral loss features, and in-source fragments. The algorithm of related peak recognition follows our previous publication^55^; 3) The remaining enriched-unlabeled features were subjected to chemical formula generation to calculate the possible carbon atoms, where chemical formulas were generated and filtered by seven golden rules^56^; 4) The enriched-unlabeled features were finally paired with enriched-labeled features according to the number of carbon atoms, mass differences (m/z error ≤ 10 ppm), and retention time differences (RT error ≤ 3 s). The output from this pipeline is provided in **Supplementary Tables 1-6**.

### Expression and purification of recombinant proteins in E. coli

Genes were amplified by PCR from *E. coli* MG1655 genomic DNA using oligonucleotide primers containing homology arms to the pET-15b vector designed to generate N-terminal hexahistidine fusion proteins (primers listed in **Supplementary Table 16**). PCR products were assembled with BamHI-digested pET-15b using NEBuilder HiFi DNA Assembly Master Mix (New England Biolabs), transformed into *E. coli* Stbl4 by electroporation, and miniprepped plasmids were sequenced to verify the correct insert. For SsnA, HyuA, YgeW, YqeA, YgeY, and YgeX, plasmids were transformed into *E. coli* BL-21 (DE3) (New England Biolabs) by electroporation, selected on LB-carbenicillin plates, and inoculated into 10 mL LB-carbenicillin pre-cultures and incubated overnight at 37 °C with shaking. Pre-cultures were diluted into 1000 mL of LB-carbenicillin in 2.8 L baffled Fernbach flasks and grown aerobically at 37 °C with shaking until reaching an OD_600nm_ of 0.4 to 0.5. Cultures were then chilled in an ice-water bath until 16 °C, isopropyl-β-D-thiogalactopyranoside (IPTG) was added (0.1 mM final), and cultures were incubated with shaking overnight at 16 °C. After ∼16 h, cells were harvested by centrifugation (4,816 x *g*, 15 min, 4 °C), supernatant was decanted, and cell pellets were re-suspended in 30 mL of binding buffer (50 mM sodium phosphate, 300 mM NaCl, pH = 7.5). Cells placed in an ice-water bath were lysed by sonication using a Fisherbrand Model 120 Sonic Dismembrator with the following settings: power, 100%; on 4 seconds; off 12 seconds; 4 minutes total on time. Cell extracts were clarified by centrifugation (7,197 x *g*, 30 min, 4 °C) and supernatant was loaded onto a 5 mL HiTrap Talon Crude column (Cytvia) using an AKTA Start Fast Protein Liquid Chromatography instrument (GE Healthcare). The column was washed with 15 column volumes of binding buffer and proteins were eluted with 100% elution buffer (50 mM sodium phosphate, 300 mM sodium chloride, 150 mM imidazole, pH = 7.5). One milliliter fractions were collected using a Frac 30 (Ge Healthcare) and protein-containing fractions were pooled, concentrated to 2.5 mL using a 10 kDa molecular weight cut-off filter, and de-salted using a PD-10 Desalting Column (Cytvia) previously equilibrated with protein storage buffer (50 mM Tris, 150 mM NaCl, pH = 7.5). Proteins were concentrated to ∼500 μL, then glycerol was added to final concentration 20% and protein aliquots were frozen at -80 °C for subsequent enzyme assays. Protein concentrations were determined using a DC Protein Assay kit (BioRad) with bovine serum albumin (BSA) as the standard.

Because YgfK is predicted to have an iron-sulfur cluster we chose to co-express it with the *sufABCDSE* overexpression plasmid (pPH151) and purify it under anaerobic conditions. The pET-15b-*ygfK* plasmid was co-transformed with the pPH151 plasmid into *E. coli* BL-21 (DE3) by electroporation, colonies were selected on LB plates supplemented with carbenicillin and chloramphenicol, and colonies were inoculated into 10 mL LB-carbenicillin/chloramphenicol pre-cultures and incubated overnight at 37 °C with shaking. Pre-cultures were diluted into 1000 mL of LB-carbenicillin/chloramphenicol supplemented with iron(III) chloride (200 μM, final) in a 2.8 L baffled Fernbach flask and grown aerobically at 37 °C with shaking until reaching an OD_600nm_ of 0.4 to 0.5. Cultures were then chilled in an ice-water bath until 16 °C, IPTG was added (0.1 mM final), cysteine hydrochloride was added (600 μM, final) and cultures were incubated with shaking overnight at 16 °C. After ∼16 h, cells were harvested by centrifugation (4,816 x *g*, 15 min, 4 °C), supernatant was decanted, and cell pellets were brought into the anaerobic chamber on ice. Binding, elution, and protein storage buffers were made anaerobic by bringing to a boil in a microwave, placing in an ultrasonic bath set on “de-gas” setting for 30 minutes, then β-mercaptoethanol was added to a final concentration of 5 mM and buffers were brought into the anaerobic chamber at least 16 h prior to use. Cells were re-suspended in 30 mL of pre-chilled anaerobic binding buffer and lysed by sonication in the anaerobic chamber. Cell extracts were clarified by centrifugation (7,197 x *g*, 30 min, 4 °C) and supernatants were loaded onto 4 mL bed volume of TALON Metal Affinity Resin (Takara Bio) previously equilibrated twice with 30 mL anaerobic binding buffer. Protein binding to resin was achieved by mixing slowly on a rotisserie rotator for 30 minutes. Then resin was centrifuged (700 x *g*, 3 min, 4 °C), flow through was decanted, and resin was washed 4 times with 35 mL volumes of binding buffer with 5-minute mixing intervals. Protein was eluted with 10 mL of anaerobic elution buffer, filtered to remove residual resin, concentrated to ∼2.5 mL using a pre-reduced 30 kDa molecular weight cut-off centrifugal concentrator, and de-salted using a PD-10 Desalting Column (Cytvia) previously equilibrated with anaerobic protein storage buffer. Protein was concentrated to ∼500 μL, glycerol was added to final concentration 20%, protein was aliquoted anaerobically in air-tight silanized glass LC-MS vials with glass inserts sealed with caps with rubber septa, and frozen at -80 °C for subsequent enzyme assays. Protein concentration was determined using a DC Protein Assay kit (BioRad) with BSA as the standard.

To assess protein purity, proteins were diluted to a concentration of ∼0.22 mg/mL, then 13 μL diluted protein was combined with 5 μL of 4X LDS sample buffer and 2 μL reducing agent. Samples were heated in a PCR machine at 70 °C for 10 minutes, then cooled to room temperature. Five microliters of either perfect protein 15-150 kDa ladder or sample was then loaded onto a Bolt™ 4-12% Bis-Tris Plus 12-well gel and electrophoresed in 1X MES buffer for 22 min at 200 V. Gels were rinsed three times with milli-Q water, then fixed with protein fixative (7% acetic acid, 40% methanol in water) for 15 minutes, washed three more times with milli-Q water, fixed for ∼3 h with GelCode stain (Thermo), and destained overnight with milli-Q water.

### Native purification of C-terminal His-tagged XdhAC from C. sporogenes

*C. sporogenes* XdhAC cultures and protein purification steps were performed under anaerobic conditions with all the reagents and plasticware pre-reduced overnight. Strains *02131xdhAC-CHis* or *selD 02131xdhAC-CHis* were cultured as described above. Overnight cultures from individual colonies were diluted 40-fold in GAM broth supplemented with 4 mM uric acid and subcultured for ∼7 h until reaching an optical density (OD_600nm_) of 2 to 2.5. Cell pellets were washed with pre-reduced 100 mM KPi buffer, pH 7.5 and stored at -80 °C until purification.

During purification, cell pellets were suspended to ∼80 OD_600nm_ in pre-reduced and pre-chilled lysing buffer (300 mM NaCl, 50 mM Na_2_HPO_4_, pH 7.5, with 5 mM β-mercaptoethanol, protease inhibitor cocktail and 20 mM imidazole) within an anaerobic chamber. Cells were sonicated in an ice-water bath (4 seconds on, 12 seconds off, total 30 min sonication). Following sonication, cell lysates were centrifuged at 7,000 x *g* for 10 min at 4 °C. The clarified cell lysate was then incubated with Complete His-Tag Purification Resin, with slow end-over-end mixing (10 rpm) for 15 min at room temperature. The resin suspension was loaded into empty PD-10 Columns (Cytiva, 17-0435-01) equipped with a frit and washed with wash buffer (300 mM NaCl, 50 mM Na_2_HPO_4_, pH 7.5, with 5 mM β-mercaptoethanol, protease inhibitor cocktail and 50 mM imidazole) three times by gravity. Subsequently, protein was eluted from the resin twice with elution buffer (300 mM NaCl, 50 mM Na_2_HPO_4_, pH 7.5, with 5 mM β-mercaptoethanol, and 250 mM imidazole) by gravity. The eluted protein was desalted using PD-10 Desalting columns being exchanged into protein storage buffer (200 mM Tris-Cl, 150 mM NaCl, pH 7.5, with 5 mM β-mercaptoethanol).

Protein samples were concentrated using Microsep Advance Centrifugal Devices with a molecular weight cut-off of 10,000 and centrifuged at 7,000 x *g* for 12 min at 4 °C. Glycerol was then added to a final concentration of 20% v/v. An aliquot was saved for protein gel analysis, while the remaining proteins were portioned into small aliquots and stored in pre-reduced LC-MS glass vials with inert glass inserts. The vials were tightly capped and stored at -80 °C. Protein concentration was determined by measuring the absorbance at 280 nm using a nanodrop instrument. Overall yield of XdhAC-Chis was ∼15 μg of purified protein per liter of culture.

### Western Blot of C-terminal His-tagged XdhAC from C. sporogenes

Protein samples and ladders were electrophoresed on a NuPAGE 4-12% Bis-Tris Plus gel with 1X MES buffer at 200 V for 22 min. The gel was then transferred to a nitrocellulose membrane at 100 V for 1.5 h at 4 °C, using 1X Tris-Glycine Transfer Buffer (Cell Signaling, #12539) supplemented with 10% methanol. Following transfer, the membrane was blocked with 5% skim milk in 1X TBS (20 mM Tris-Cl, 140 mM NaCl, pH 7.5) for 45 min at room temperature, and subsequently washed with 1X TBS. Next, the membrane was incubated with 6x-His-Tag Antibody (Thermo Fisher, MA1-21315-HRP) diluted 500-fold in 1% skim milk in 1X TBST (1X TBS containing 0.1% tween 20) at room temperature for 2 h. Following the incubation, the nitrocellulose membrane underwent three washes with 1X TBST before immune detection using SuperSignal West Pico PLUS Chemiluminescent Substrate and imaging with an iBright imaging system (Thermo).

### In vitro enzyme assays

All the procedures were conducted within the anaerobic chamber unless otherwise specified.

#### Generation of pathway intermediates from UA mutant supernatants

To obtain pathway intermediates, which were commercially unavailable, we utilized *C. sporogenes* uric acid mutants as a reliable source. The mutants were cultured in GAM broth supplemented with 4 mM uric acid until an OD_600nm_ of ∼2.5, following the procedure described earlier. The cultures were then harvested and washed once with 1X M9 salts. Cell pellets, adjusted to a final OD_600nm_ of 25, were resuspended in 1X M9 salt containing 2 mM uric acid. These suspensions were incubated at 37 °C within an anaerobic incubator for 24 h. Subsequently, the supernatants were collected and filtered through 10 kDa MWCO filter at 7,000 x *g* for 45 min at room temperature. The resulting flow-through was transferred to pre-reduced LC-MS silanized glass vials, tightly capped, and stored at -80 °C. After confirmation of the presence of specific intermediates by LC-MS, these supernatants were utilized as a source of pathway intermediates and were designated as UA pathway intermediate supernatants. We also analyzed pathway intermediates in *E. coli* concentrated cell suspensions using a similar method to *C. sporogenes*, but with minor differences. Specifically, cells were grown in GAM.M broth supplemented with uric acid for 45 h prior to harvesting, washing, and resuspending at a final OD_600nm_ of ∼20.

#### Verification of purified enzyme activity and determination of order of enzymes in pathway

To assess the activity of purified *E. coli* enzymes (YgfK, SsnA, HyuA, YgeW, YgeY and YgeX), UA intermediate supernatants were combined with each purified enzyme (final 0.2 mg/mL) at 4:1 ratio. The mixtures were incubated at 37 °C in an anaerobic chamber for 3 h. In the YgfK assays, final 2 mM each of NAD^+^, NADH, NADP^+^ and NADPH were included in the reactions. Aliquots of reactions were taken at 1 h and 3 h and preserved for LC-MS measurement.

#### Purified YgeX assay

Culture supernatants from *C. sporogenes ygeX* (grown in GAM supplemented with 4 mM UA or [^13^C_5_]-UA for 48 h) and *E. coli ygeX* (grown in GAM.M supplemented with 4 mM UA or [^13^C_5_]-UA for 72 h) were harvested and filtered through 3 kDa Nanosep filters at 20,000 x *g* for 30 min. Following filtration, BSA (final 0.2 mg/mL) or purified *E. coli* YgeX protein (final 0.2 mg/mL) was added to the culture supernatants. The mixtures were then incubated aerobically in a ThermoMixer at 37 °C for 30 min. The reaction mixtures were snap frozen and stored at -80 °C until LC-MS analysis.

#### Identification of cofactor requirements and the substrate of YgfK

To ascertain which nicotinamides are necessary for YgfK function, separate stocks of 50 mM NAD^+^, NADH, NADP^+^ and NADPH were freshly prepared in pre-reduced 100 mM KPi buffer at pH 7.5. Each of these stocks (final 4 mM) was then combined with UA-2 intermediate supernatant and purified YgfK (final 0.2 mg/mL). The reaction mixtures were incubated anaerobically at 37 °C for 1 h, and aliquots were collected for LC-MS analysis at designated time points.

To determine the authentic substrate for YgfK, a combination of xanthine, 2,8-dioxopurine, and 6,8-dioxopurine was diluted in pre-reduced 100 mM KPi buffer, pH 7.5. Xanthine isomers (final 1 mM each) were then mixed 4 mM NADPH and 0.2 mg/mL YgfK, with or without sodium selenate (final 2 mM). The reaction mixtures were incubated anaerobically at 37 °C for 1 h, and aliquots were saved for LC-MS analysis at designated time points.

#### Determination of C. sporogenes XdhAC activity

For the reverse reaction, a mixture of xanthine isomers (final 1 mM each) was combined with methyl viologen (oxidized form, final 10 mM) in 100 mM KPi buffer, pH 7.5. For the forward reaction, uric acid (final 1 mM) was combined with a mixture of methyl viologen/dithionite (final 1 mM/10 mM) in the same buffer. Following the addition of purified *C. sporogenes* XdhAC (final 0.001 mg/mL), the reaction mixtures were incubated anaerobically at 37 °C for 3 h. Aliquots were collected at designated time points for LC-MS analysis.

#### In vitro reconstitution

All steps were performed in an anaerobic chamber with pre-reduced buffers and supplies. A fresh uric acid stock was diluted in pre-reduced 100 mM KPi buffer, pH 7.5, supplemented with methyl viologen (final 0.2 mM), dithionite (final 4 mM), and NADPH (final 4 mM). Enzyme working stocks were created by combining them with MnCl_2_ (final 16 μM) in 100 mM KPi buffer, pH 7.5, containing 1 mM dithionite, prior to addition of the uric acid substrate. The final enzyme concentrations in the reactions were as follows: 0.001 mg/mL of *C. sporogenes* XdhAC, 0.2 mg/mL of *E. coli* YgfK, 0.02 mg/mL of *E. coli* SsnA, 0.02 mg/mL of *E. coli* HyuA, 0.2 mg/mL of *E. coli* YgeW, 0.2 mg/mL of *E. coli* YgeY and 0.02 mg/mL of *E. coli* YgeX. When necessary, 0.04 mg/mL of *E. coli* YqeA, along with 4 μM MgSO_4_ and 2 mM ADP, were added in the reaction mixtures. YqeA was added along with YgeW because we found that it stimulated YgeW activity, likely by consuming carbamoyl phosphate and relieving end-product inhibition of YgeW. The reactions were initially set up on ice before incubating at 37 °C for 1 h. At designated time points, aliquots of the reaction mixtures were mixed with pre-reduced ISTD in extraction solution to quench the reaction. Then they were centrifuged under anaerobic conditions, and the extraction supernatant was used for LC-MS analysis.

#### Comparison of bovine XOR and C. sporogenes XdhAC activity

To investigate the cofactor requirements and substrate specificity of *C. sporogenes* XdhAC and bovine milk XOR (Sigma), enzyme assays were performed to assess their ability to catalyze the oxidation of xanthine isomers and the reduction of uric acid.

### Anaerobic Assays

Fresh stocks of the following compounds were prepared in pre-reduced 100 mM KPi buffer, pH 7.5: dithionite (400 mM), methyl viologen (200 mM), NAD^+^ (50 mM), NADH (50 mM), NADPH (50 mM) and NADP^+^ (50 mM).

Oxidation of xanthine isomers. Reaction mixtures contained individual xanthine isomers (final 1 mM), *C. sporogenes* XdhAC (final 0.001 mg/mL) or bovine milk XOR (final 1 U/mL), and one of the following electron acceptors: methyl viologen (oxidized form, final 10 mM), or NAD+ (final 10 mM), or NADP+ (final 10 mM).

Reduction of uric acid. Reaction mixtures contained uric acid (final 1 mM), *C. sporogenes* XdhAC (final 0.001 mg/mL) or bovine milk XOR (final 1 U/mL), and one of the following electron donors: methyl viologen/dithionite (final 1mM/10 mM), or NADH (final 10 mM), or NADPH (final 10 mM).

Reactions were incubated anaerobically at room temperature for 3 hours. Aliquots were collected at designated time points for LC-MS analysis.

### Aerobic Assays

To assess enzyme activity under aerobic conditions, assays were performed by incubating individual xanthine isomers (final 1 mM) with *C. sporogenes* XdhAC (final 0.001 mg/mL) or bovine milk XOR (final 0.2 U/mL) on the bench top (exposed to air) for 1 hour at room temperature. Aliquots were collected at designated time points for LC-MS analysis.

#### Competition experiments for wild-type and DOPDH (xdhAC) mutant C. sporogenes in gnotobiotic mice

C57BL/6 germ-free mice (Taconic Biosciences, *Mus musculus*, Tac:B6) were bred and maintained in-house within aseptic flexible film isolators (CBC, Madison, WI). Mice were housed under a 12-hour light/dark cycle at a controlled temperature (20-22 °C) and humidity (40-60%), with ad libitum access to standard chow (LabDiet Cat. # 5K67) and sterile water. Germ-free status was confirmed before each experiment by anaerobic culture of fecal pellets from each mouse in GAM medium for 24 h. All animal procedures were approved by the Stanford University Administrative Panel on Laboratory Animal Care.

In this study, eight male C57BL/6 germ-free mice (6–8 weeks old) were placed on a 3% oxonic acid/2% uric acid diet (Inoviv, TD.220052, **Supplementary Data File 3**) one day prior to bacterial colonization. Mice were colonized with a mixture of wild-type and *xdhAC* mutant *C. sporogenes* by oral gavage (200 μL, ∼1 × 10^10^ CFU). Fresh fecal pellets were collected and stored at -80 °C until DNA extraction was performed.

To generate the inoculum, wild-type and *xdhAC* mutant *C. sporogenes* strains were individually cultured anaerobically overnight in GAM broth. Equal volumes of overnight cultures of the two strains were combined, concentrated 7-fold, and administered to individual mice via intragastric gavage. A sample of the inoculum was retained for DNA extraction and subsequent qPCR analysis to determine the initial strain ratio which showed the actual ratio at time of gavage was 1:2.4 (wild-type:*xdhAC*).

Total genomic DNA (gDNA) was extracted from fecal samples (3-20 mg) and the gavage mixture using QIAamp PowerFecal Pro DNA Kit (QIAGEN, 51804) according to manufacturer’s guidelines. DNA concentration was measured by Qubit™ dsDNA Quantification Assay Kit (ThermoFisher Scientific, Q32853). Two primer sets were designed for *C. sporogenes* strains (**Supplementary Table 16**): one specific to the ClosTron-disrupted *xdhAC* gene (*xdhAC*_CT) and the other targeting the wild type *xdhAC* gene (*xdhAC*_WT). Q-PCR was performed using PowerUp SYBR Green Master Mix (ThermoFisher Scientific, A25742) with four replicates and an Applied Biosystems QuantStudio 5 real-time PCR instrument (ThermoFisher Scientific). Absolute quantification was used to determine the amount of *xdhAC*_CT and *xdhAC*_WT in each sample. Serially diluted standard curves (2 fg/μL - 2 ng/μL) of purified gDNA from wild type *C. sporogenes* (xdhAC_WT) and *xdhAC C. sporogenes* (xdhAC_CT) were included in each qPCR run. The concentration of each strain was determined by interpolating sample Cq value onto the corresponding standard curve. The competitive index (CI) was calculated according to equation 1 below:

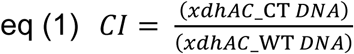

### Identification of uric acid degrading genes in metagenome assembled genomes

We used BLASTp searches to identify uric acid genes in metagenome assembled genomes (MAGs) from the Earth’s Microbiomes^39^, chicken gut^40^, and termite gut^41^ datasets. The protein fasta files were downloaded from the respective databases and used to generate individual protein BLAST libraries using Geneious Prime (ver. 2024.0.2). Each of the seven *C. sporogenes* uric acid proteins (XdhAC, YgfK, SsnA, HyuA, YgeW, YgeY, and YgeX) were used as queries for a BLASTp search of the databases. Results were filtered based minimum sequence coverage of 80% and predefined % amino acid cutoffs derived from phenotypic analyses of uric acid degrading bacteria^3^. These cutoffs were as follows: XdhAC, 46%; YgfK, 41%; SsnA, 33%; HyuA, 36%; YgeW, 63%; YgeY, 53%; YgeX, 50%. MAGs were considered positive for the uric acid genes if at least 4/7 proteins were present. Metadata for positive MAGs were retrieved from the Earth’s Microbiomes study and MAGs containing uric acid genes were grouped into ecosystem categories and placed on the world map using the ggplot package in R studio.

### Identification of selenium utilization traits in human gut bacterial strain library

We previously generated a strain library of human gut bacteria, most of which had publicly available genome sequences^3^. To identify selenium utilization traits in these bacteria, we downloaded the protein fasta sequences from these genomes, concatenated the files, and generated a protein BLAST database using Geneious Prime (ver. 2024.0.2). We then performed BLASTp searches of this database using the following *E. coli* MG1655 proteins: SelD (NCBI accession AAC74834.1) as a marker for selenium utilization; SelA (NCBI accession AAC76615.1) as a marker for the selenocysteine decoding trait; SelU (NCBI accession AAC73605.1) as a marker for selenouridine incorporation; YqeB (NCBI accession AAC75913.1) and YqeC (NCBI accession AAC75914.2) as markers for selenium co-factor trait. We also performed BLASTp searches with the *C. sporogenes* uric acid proteins XdhAC (NCBI accession EDU35963.1), YgfK (NCBI accession EDU35960.1), SsnA (NCBI accession EDU35961.1), HyuA (NCBI accession EDU35958.1), YgeW (NCBI accession EDU35959.1), YgeY (NCBI accession EDU35962.1), and YgeX (NCBI accession EDU35956.1) and xanthine dehydrogenase XdhA (NCBI accession EDU37525.1). Results were filtered based minimum sequence coverage of 80% and the following amino acid sequence identities: EcSelD, 34%; EcSelA, 37%; EcSelU, 26%; EcYqeB, 41%; EcYqeC, 22%; CsXdhA, 31%. Criteria to assign genomes as positive for uric acid genes included a minimum of 4/7 of the following proteins with corresponding amino acid sequence identities: XdhAC, 46%, YgfK, 41%, SsnA, 33%, HyuA, 36%, YgeW, 63%, YgeY, 53%, and YgeX, 50%.To generate the phylogenetic tree, RpoB protein sequences were retrieved from 187 organisms in our library, uploaded into Geneious Prime, and a Muscle alignment was performed with default parameters. Gaps were then manually trimmed, and sequences were re-aligned with Muscle. A neighbor joining unrooted phylogenetic tree was generated from the alignment and exported as a newick file, then uploaded into the interactive tree of life (iTOL). The presence or absence of SelD, SelA, SelU, YqeB/C, UA genes, and XdhABC was then mapped onto the tree using the iTOL annotation editor.

### Phylogenetic analysis of molybdenum binding domains using CLANS

To compare the phylogenetic distribution of molybdenum hydroxylases, we chose to focus on the molybdenum-binding domains which form the substrate binding active sites of these enzymes. First, we searched the Protein Data Bank and experimentally annotated Swiss-Prot databases to identify biochemically or structurally characterized enzymes. These searches revealed nine key enzymes, bovine xanthine oxidoreductase (XOR), *Rhodobacter capsulatus* xanthine dehydrogenase (XDH), *Clostridium sporogenes* 2,8-dioxopurine dehydrogenase (DOPDH), *C. sporogenes* xanthine dehydrogenase representative of gut bacteria (gbXDH), *Megalodesulfovibrio gigas* aldehyde dehydrogenase (AOR), *Eubacterium barkeri* nicotinate dehydrogenase (NDH), *Paenarthrobacter nicotinovorans* ketone dehydrogenase (KDH), *Pseudomonas putida* quinoline 2-oxidoreductase (QorL), and *Hydrogenophaga pseudoflava* carbon monoxide dehydrogenase (CODH). For each of these proteins, we expanded the group by performing a BLASTp search of the NCBI database and retrieving 9-30 hits for subsequent analyses (**Supplementary Table 17**). Each group was aligned to the molybdenum binding domains of *C. sporogenes* DOPDH (amino acids 307 – 839) and non-aligning residues were trimmed. Note that for *E. bacterium* NDH and homologs, the two molybdenum-binding domains are often separated in two polypeptides, so individual alignments were made with DOPDH for each polypeptide, and the aligned sequences were concatenated into once sequence. These sequences were then used as a query in the CLuster ANalysis of Sequences (CLANS) software^57^ implemented on the MPI Bioinformatics Toolkit web-server^58^. The resulting file was opened with CLANS software (v.29-05-2012) and ∼50,000 iterations were performed using the following settings: complex attraction, cluster in 2D, antialiasing, show connections. The force-directed graph was exported as a .pdf and nodes were colored according to protein family in Adobe Illustrator.

### Synthesis of UA-5 and UA-6 by carbamation of 2,3-diaminopropanoic acid

To synthesize UA-5 and UA-6 from diaminopropanoic acid, we adapted a previously published approach using urea, amines, and heat under aqueous conditions ^59^. Urea (final 2 M) and 2,3-diaminopropanoic acid hydrochloride (final 0.5 M) were mixed in LC-MS grade water and incubated at 80 °C for 48 h. This reaction produced a mixture containing UA-6 and UA-5 intermediates (as determined by LC-MS). UA-6 reached its peak concentration at 12 h, while UA-5 achieved its highest level at 48 h. Aliquots of the reaction mixture were withdrawn at specified intervals for subsequent LC-MS analysis.

### LC-MS data supporting figures

All targeted LC-MS data corresponding to main and extended data figures are provided in **Supplementary Tables 18-33.**

## Data availability

All data are available in the main text or the supplementary materials. Code used for stable isotope tracing is available on GitHub (https://github.com/DoddLab/DoddLabTracer). Strains and plasmids generated in this study are available from the corresponding author without restriction.

## Acknowledgements

We thank Michael Fischbach for valuable discussions. We also thank Linh Thuy Nguyen for assistance with western blots and Cherelle Soriano and Steven Higginbottom for help with gnotobiotic experiments. We thank Tyler Grove for sharing the pPH151 plasmid. This work was funded in part by National Institutes of Health grants K08-DK110335 (D.D.), R35-GM142873 (D.D.), R01-AT011396 (D.D.), the Stanford Microbiome Therapies Initiative (D.D.), an OHF-ASN Foundation for Kidney Research Career Development Award (D.D.), a Stanford Medicine Dean’s Postdoctoral Fellowship (Z.Z.), a Stanford Medicine Children’s Health Center for IBD and Celiac Disease Postdoctoral and Early Career Support Award (Z.Z.), and the Stanford PROPEL Scholars Program (M.M-V.).

## Author contributions

Y.L., J.B.J., and D.D. conceived and designed the project; J.B.J. and H.C. performed initial pathway characterization which was expanded upon by Y.L., Z.Z., and D.D.; Y.L. conducted and analyzed all growth and *in vitro* enzyme assays and LC-MS sample preparation and targeted LC-MS data analysis; H.C., Z.Z., and D.D. developed LC-MS assays; Z.Z. and D.D. performed LC-MS data acquisition; Z.Z. developed code for untargeted data analysis with help from D.D.; D.D. cloned genes, produced and purified recombinant proteins; Y.L. performed CRISPR gene editing and produced and purified native protein; D.D. analyzed genes in metagenomic datasets and the strain library with help from Z.Z.; M.M. coordinated synthesis of authentic standards; R.T. provided intellectual contributions into urate and purine biology and translation; Y.L. performed gnotobiotic mouse experiments with help from D.D.; Y.L., Z.Z., and D.D. prepared figures and wrote the manuscript with input from all co-authors.

