## Supplementary Information and Extended Data Figures for "Gut bacteria degrade purines *via* the 2,8-dioxopurine pathway"

Supplementary Text
Extended Data Figures 1-10
Supplementary Figures 1-4
Supplementary Data files 1-3

**Supplementary text:**

**Data supporting structural assignment of intermediates in the pathway.** Our current proposal for the pathway for uric acid metabolism in gut bacteria is presented in **Figure 2k**. There are eight intermediates in the pathway which we have designated UA-1 through UA-8. Below is a summary of the data supporting the assignment of each compound in the pathway.

*UA-1, uric acid, 7,9-dihydro-1*H*-purine-2,6,8(3*H*)-trione.* Authentic standards of unlabeled uric acid and $[^{13}C_5]$-labeled uric acid were obtained from commercial vendors. When used as substrates, both labeled and unlabeled uric acid were consumed by wild-type *C. sporogenes* and $[^{13}C_5]$-labeled uric acid was converted to labeled SCFAs, acetate and butyrate (**Extended Data Figure 1c** and **Figure 2c**). In addition, unlabeled uric acid was used as a substrate by *C. sporogenes* DOPDH (XdhAC) (**Figure 2i**) and was converted to other pathway intermediates during *in vitro* reconstitution of the pathway (**Figure 2k**).

*UA-*2*, 2,8-dioxopurine, 7,9-dihydro-2*H*-purine-2,8(3*H*)-dione*. An authentic standard of 2,8-dioxopurine matched UA-2 that accumulated in the supernatants of the *C. sporogenes ygfK* mutant by comparison of retention time, accurate mass-to-charge ratio, and MS/MS spectrum (**Figure 2h** and **Extended Data Figure 6c**). In addition, 2,8-dioxopurine was used as a substrate for *C. sporogenes* DOPDH in the reverse direction (**Extended Data Figure 6g**), for *C. sporogenes* DOPOR (YgfK) in the forward direction (**Figure 2j**), and accumulated during *in vitro* reconstitution of the pathway when DOPDH was incubated with uric acid and disappeared when DOPOR was also included (**Figure 2k**).

*UA-3,* ***provisional assignment*** *1,6,7,9-tetrahydro-2*H*-purine-2,8(3*H*)-dione*. A metabolic feature with m/z of 155.0558 ($[M+H]^+$) and retention time by HILIC of 3.29 minutes accumulated in the supernatants of the *C. sporogenes* and *E. coli ssnA* mutants (**Figure 1b**, **Extended Data Figure 2a-b and Extended Data Figure 4c**). The same compound was produced when DOPOR was incubated with UA-2 intermediate supernatants (**Extended Data Figure 5d** and **Extended Data Figure 6k**) and authentic 2,8-dioxopurine (UA-2) (**Figure 2j** and **Extended Data Figure 6i**). The compound disappeared when UA-3 intermediate supernatants were incubated with SsnA (**Extended Data Figure 5d**). In addition, this compound accumulated during *in vitro* reconstitution of the pathway when DOPDH and DOPOR were incubated with uric acid and disappeared when SsnA was also included (**Figure 2k**). The compound was unstable in sample matrix, completely disappearing after storage of culture supernatants stored in 96-well plates with a lid in the freezer (-80 °C) for 2 weeks. To consistently detect UA-3, we made all attempts to limit the time between sample preparation and LC-MS analysis.

Stable isotope tracing demonstrated that this compound contains 5 carbon atoms (**Figure 1b**, **Extended Data Figure 2a-b** and **Extended Data Figure 4c**). Combining this information with the compound's accurate mass, and assuming only carbon, hydrogen, nitrogen, and oxygen atoms, we assigned its molecular formula as $C_5H_6N_4O_2$. DOPOR is an oxidoreductase that uses NADPH as a co-factor and the mass difference between 2,8-dioxopurine (UA-2) and UA-3 corresponds to the addition of two hydrogen atoms. Therefore, we predict that DOPOR reduces either the $N_1$-$C_6$ or $C_4$-$C_5$ double bonds present in 2,8-dioxopurine (UA-2) using a hydride from NADPH and a proton from solution. Although we have no direct data to support which double bond is reduced, we favor that it is the $N_1$-$C_6$ because the resulting intermediate could then be attacked by the next enzyme in the reaction sequence (SsnA) to open the six-membered ring in a reaction similar to that previously demonstrated during catalytic hydrogenation of 2,8-dioxopurine in aqueous acid[1]. Confirmation of the identity of this intermediate will likely require its purification from reactions of DOPOR with 2,8-dioxopurine (UA-2). However, given the instability of this compound, purification and structural determination is beyond our lab's current capabilities.

*UA-4, **provisional assignment** 1-((2,5-dioxoimidazolidin-4-yl)methyl)urea.* Two metabolic features with m/z of 173.0668 ($[M+H]^+$) accumulated in the supernatants of the *C. sporogenes hyuA* mutant (**Figure 1b** and **Extended Data Figure 2a-b**). The first feature showed a sharp peak centered at retention time of 3.36 min; the second peak had a broad tail and was centered at a retention time of 3.51 min (**Figure 1b**). When UA-4 intermediate supernatants were incubated with HyuA, only the second broader peak disappeared, being converted to UA-5 (**Supplementary Figure 1a**). Therefore, the second peak centered at 3.51 minutes is likely the true pathway intermediate which we designate UA-4. This compound also accumulated in the supernatants of the *E. coli ssnA* mutant (**Extended Data Figure 4c**). Stable isotope tracing demonstrated that this compound contains 5 carbon atoms (**Figure 1b**, **Extended Data Figure 2a-b** and **Extended Data Figure 4c**). Combining this information with the compound's accurate mass, and assuming only carbon, hydrogen, nitrogen, and oxygen atoms, we assigned its molecular formula as $C_5H_8N_4O_3$. We initially thought that UA-4 might arise from cleavage of the 5-membered ring of UA-3 by SsnA and purchased 1-(2,4-

dioxohexahydropyrimidin-5-yl)urea. However, this compound did not match UA-4 in retention time (**Supplementary Figure 1b**) or mass fragmentation pattern (**Supplementary Figure 1c**). Instead, the 1-(2,4-dioxohexahydropyrimidin-5-yl)urea standard matched the first peak at 3.36 minutes (**Supplementary Figure 1b-d**). Given these findings, we propose that UA-4 represents a cleavage product of the 6-membered ring of UA-3 by SsnA (**Figure 2k**).

*UA-5, (R)-2,3-diureidopropanoic acid.* A metabolic feature with m/z of 191.0747 ($[M+H]^+$) and retention time by HILIC of 5.42 minutes accumulated in the supernatants of the *C. sporogenes* and *E. coli ygeW* mutants (**Figure 1b**, **Extended Data Figure 2a-b** and **Extended Data Figure 4c**). The same compound was produced when HyuA was incubated with UA-4 intermediate supernatants (**Extended Data Figure 5d**). In addition, this compound accumulated during *in vitro* reconstitution of the pathway when DOPDH, DOPOR, SsnA, and HyuA were incubated with uric acid and was diminished when YgeW was also included (**Figure 2k**). Stable isotope tracing demonstrated that this compound contains 5 carbon atoms (**Figure 1b**, **Extended Data Figure 2a-b** and **Extended Data Figure 4c**). Combining this information with the compound's accurate mass, and assuming only carbon, hydrogen, nitrogen, and oxygen atoms, we assigned its molecular formula as $C_5H_{10}N_4O_4$. We used [$^{13}C_5$]-uric acid stable isotope tracing in combination with NPH derivatization (for free carboxylates) and identified a single derivatized peak present in the *C. sporogenes ygeW* mutant cultures that was not present in the wild-type cultures (**Supplementary Figure 1e-g**). This data supports the presence of a free carboxylic acid in UA-5. We then used a previously published synthetic method for carbamation of free amines[2] to produce UA-5 using DL-2,3-

diaminopropanoic acid and urea as starting reagents (**Supplementary Figure 1h**). After 24 h of incubation, a compound with the same retention time and accurate mass-to-charge ratio appeared, becoming the dominant peak by 48 hours (**Supplementary Figure 1i**). Tandem mass spectrometry showed that this compound matches the MS/MS spectrum of UA-5 in *C. sporogenes ygeW* mutant culture supernatants (**Supplementary Figure 1j**). To provide insight into the stereochemistry of this compound, we repeated this synthesis using either (*R*)- or (*S*)-2,3-diaminopropanoic acid (DPA) as the starting material. We then used the synthesized products as the substrate for their corresponding enzymes in the pathway. Carbamate kinase (YqeA) was added to YgeW incubations because we found that it stimulated YgeW activity, likely by consuming carbamoyl phosphate and relieving end-product inhibition of YgeW. While YgeW and YqeA readily converted UA-5 synthesized from (*R*)-DPA to UA-6 (**Supplementary Figure 2a**), no activity was seen toward UA-5 synthesized from (*S*)-DPA (**Supplementary Figure 2b**). Furthermore, when YgeY was also added, UA-7 appeared only in reactions where UA-5 was synthesized from (*R*)-DPA (**Supplementary Figure 2a-b**). Subsequently, both the (*R*) and (*S*) enantiomers of 2,3-diureidopropanoic acid were synthesized by KareBay Biochem (**Supplementary Figure 2c** and **Supplementary Data File 2**). Both enantiomers had retention time and MS/MS spectrum matches with UA-5 in *C. sporogenes ygeW* mutant culture supernatants (**Supplementary Figure 2d-e**). However, only (*R*)-2,3-diureidopropanoic acid was a substrate for YgeW, being converted to 2,3-diaminopropanoic acid when incubated with both YgeW and YgeY (**Supplementary Figure 2f**).

*UA-6, (R)-3-amino-2-ureidopropanoic acid.* A metabolic feature with m/z of 148.0715 ($[M+H]^+$) and retention time by HILIC of 6.44 minutes accumulated in the supernatants of the *C. sporogenes* and *E. coli ygeY* mutants (**Figure 1b**, **Extended Data Figure 2a-b** and **Extended Data Figure 4c**). The same compound was produced when YgeW was incubated with UA-5 intermediate supernatants (**Extended Data Figure 5d**). In addition, this compound accumulated during *in vitro* reconstitution of the pathway when DOPDH, DOPOR, SsnA, HyuA, and YgeW were incubated with uric acid and disappeared when YgeY was also included (**Figure 2k**). Stable isotope tracing demonstrated that this compound contains 4 carbon atoms (**Figure 1b**, **Extended Data Figure 2a-b** and **Extended Data Figure 4c**). Combining this information with the compound's accurate mass, and assuming only carbon, hydrogen, nitrogen, and oxygen atoms, we assigned its molecular formula as $C_4H_9N_3O_3$. Derivatization with NPH (for free carboxylates) and DNS (for free amines) both yielded a single derivative of this compound (**Supplementary Figure 3a-f**; *note that a second NPH peak was found to be an in-source fragment of UA-5*), indicating the presence of a carboxylic acid and a free amine. We used a previously published synthetic method for carbamation of free amines to produce this compound using 2,3-diaminopropanoic acid and urea as starting reagents (**Supplementary Figure 1h**). Within 6 h of incubation, two overlapping peaks with the same accurate mass-to-charge ratio as UA-6 appeared, becoming most abundant by 12 h then decreasing in intensity over time as the reaction proceeded (**Supplementary Figure 1i**). Tandem mass spectrometry showed that this doublet peak matches the MS/MS spectrum of UA-6 in *C. sporogenes ygeY* mutant culture supernatants (**Supplementary Figure 1k**). We reasoned that the two peaks likely result from carbamation of 2,3-diaminopropanoic acid at either the 2- or 3- position, forming a mixture of 3-amino-2-ureidopropanoic acid and 2-amino-3-ureidopropanoic acid. It follows that only one of these compounds is likely to be a substrate for YgeY. To test this, we incubated 12 h synthetic reaction mixtures with YgeY and monitored UA-6 and UA-7 levels by LC-MS. Over time, the second (right-most) peak decreased in intensity and UA-7 was produced (**Supplementary Figure 3g-h**). To distinguish whether UA-6 has the ureido group at the 2- or 3-position, we performed stable isotope tracing with [1,3-$^{15}N_2$]-uric acid in cell suspensions of the *C. sporogenes ygeY* mutant. If YgeW cleaves the 2-ureido group of UA-5, the M+2 isotopolog of UA-6 will be detected in the *ygeY* mutant; however, if YgeW cleaves the 3-ureido group, the M+1 isotopolog of UA-6 will appear (**Supplementary Figure 3i**). Our data showed that only the M+1 isotopolog of UA-6 appeared in the *ygeY* mutant (**Supplementary Figure 3j**), demonstrating that YgeW cleaves the 3-ureido group and supporting the assignment of UA-6 as 3-amino-2-ureidopropanoic acid. As shown in **Supplementary Figure 3g-h**, YgeY has activity with both enantiomers of UA-6. However, given the stereoselectivity of YgeW on (*R*)-2,3-diureidopropanoic acid, it is likely that UA-6 is also in the (*R*) configuration. Both the (*R*) and (*S*) enantiomers of 3-amino-2-ureidopropanoic acid were synthesized by KareBay Biochem (**Supplementary Figure 4a** and **Supplementary Data File 3**). Both enantiomers displayed retention time and MS/MS spectra that matched UA-5 in *C. sporogenes ygeY* mutant culture supernatants (**Supplementary Figure 4b-c**). Both enantiomers of UA-6 served as substrates for YgeY (**Supplementary Figure 4d**), however only the *R*-enantiomer was converted to UA-5 by YgeW (reverse reaction) when incubated with carbamoyl phosphate (**Supplementary Figure 4d**). Therefore, our data provide evidence on UA-6 stereochemistry, indicating that this intermediate is (*R*)-3-amino-2-ureidopropanoic acid.

*UA-7, 2,3-diaminopropanoic acid.* An authentic standard of 2,3-diaminopropanoic acid was obtained from Sigma (Cat. # 219630). This compound matched UA-7 that accumulated in the supernatants of the *C. sporogenes* and *E. coli ygeX* mutant by comparison of retention time, accurate mass-to-charge ratio, and MS/MS spectrum (**Figure 2a-b**). In addition, 2,3-diaminopropanoic acid was used as a substrate for *E. coli* YgeX (**Figure 2d**) and accumulated during *in vitro* reconstitution of the pathway when DOPDH, DOPOR, SsnA, HyuA, YgeW, and YgeY were incubated with uric acid and disappeared when YgeX was also included (**Figure 2k**).

*UA-8, pyruvic acid, 2-oxopropanoic acid.* An authentic standard of sodium pyruvate was obtained from Sigma (Cat. # P2256). This compound matched UA-8 that was produced during incubation of *E. coli* YgeX with 2,3-diaminopropanoic acid by comparison of retention time and accurate mass-to-charge ratio (**Figure 2d**). In addition, pyruvic acid accumulated during *in vitro* reconstitution of the pathway when all enzymes were incubated with uric acid (**Figure 2k**).

#### Extended Data Figures

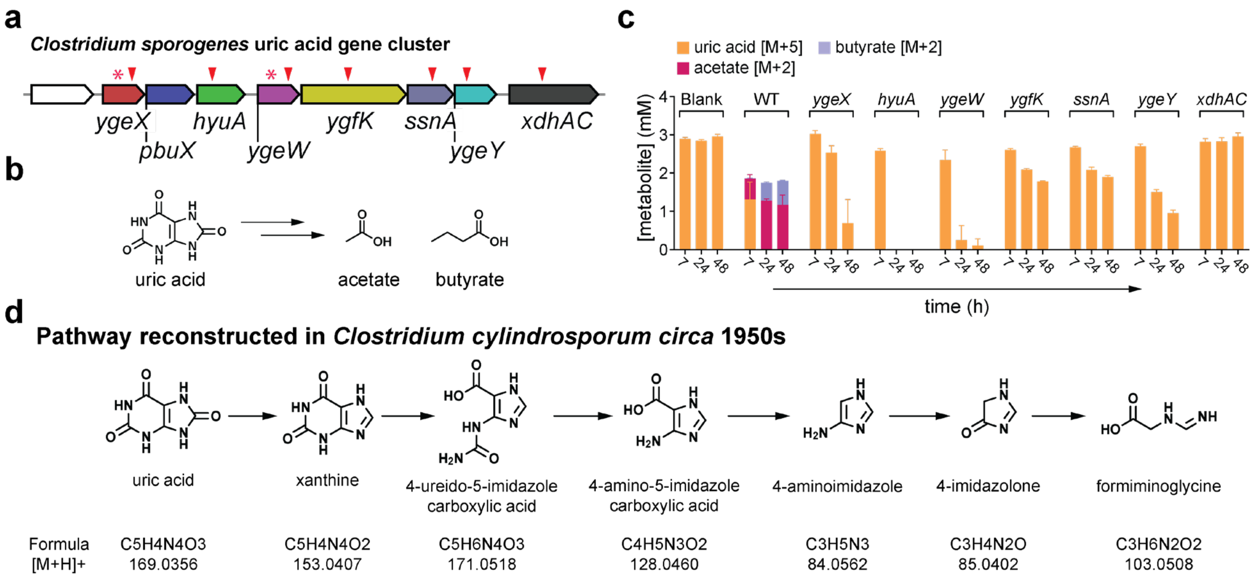

**Extended Data Figure 1. Uric acid metabolism by *C. sporogenes* mutants and overview of previously published anaerobic pathway for uric acid metabolism.** (**A**) Schematic showing the gene cluster for *C. sporogenes,* and the mutants tested in this study. Arrows indicate ClosTron mutants and asterisks indicate genes that were also deleted by CRISPR due to polar effects of ClosTron mutants. (**B**) The uric acid genes are required for synthesis of short-chain fatty acids from uric acid by gut bacteria, but the pathway is not known. (**C**) Stable isotope tracing in wild-type and mutant strains of *C. sporogenes* using Poroshell positive (uric acid) and NPH (acetate and butyrate) LC-MS. (**D**) Pathway for anaerobic uric acid reconstructed from cell extracts of the obligate purine degrading bacterium *C. cylindrosporum*, in previous studies. The predicted hydrogen adducts of each compound were used to search through untargeted metabolomics data we collected from both *C. sporogenes* and *E. coli* mutants. Although uric acid and xanthine were identified, the remaining intermediates were not detected in the *C. sporogenes* or *E. coli* mutants.

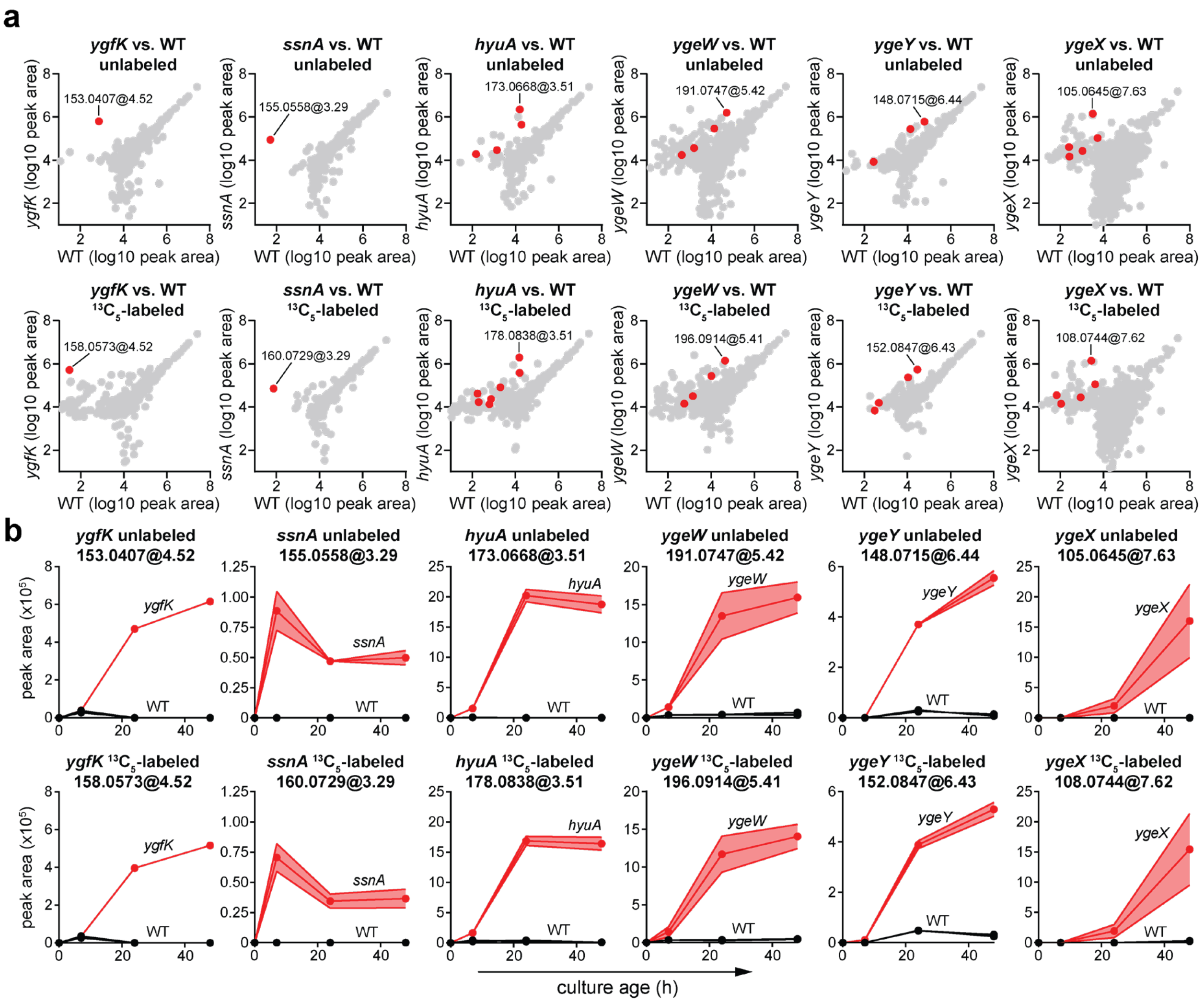

**Extended Data Figure 2. Isotope tracing in *C. sporogenes* mutants.** (**A**) Scatterplots comparing abundances of metabolites identified by HILIC LC-MS in wild-type and mutant *C. sporogenes* cultures. The top panels are for cultures with unlabeled uric acid and the bottom panels are for cultures with $^{13}C$-labeled uric acid. The 7-hour timepoint is shown for the *ssnA* mutant, whereas for all other mutants, the 48-hour timepoint is shown. Red dots indicate metabolite feature pairs that were filtered through our isotope tracing analysis. The most abundant feature in the mutants is labeled. (**B**) Peak areas for most abundant features from (**A**) over time during growth of *C. sporogenes* wild-type and mutant in rich medium supplemented with unlabeled uric acid (top panels) or [$^{13}C_5$]-labeled uric acid (bottom panels). For **B**, data represent means ± standard deviations from three replicates.

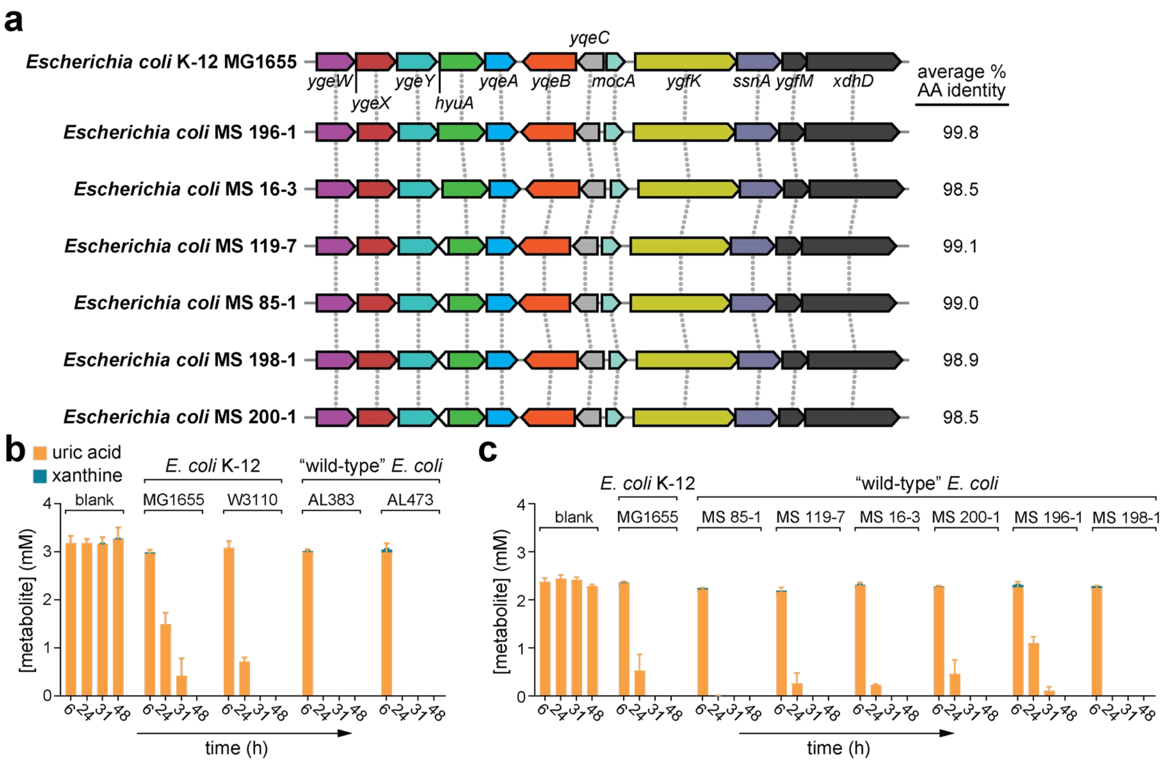

**Extended Data Figure 3. Uric acid gene clusters and in vitro uric acid consumption in K-12 and "wild-type" strains of *E. coli*.** (**A**) Genomic context of uric acid gene clusters from different strains of *E. coli*. The average percentage amino acid identity of the protein encoding genes relative to the *E. coli* K-12 MG1655 strain are shown on the right. (**B-C**) Uric acid consumption in modified Gifu Anaerobic Medium by two K-12 derivative strains and eight "wild-type" *E. coli* strains. For **A**, NCBI GenBank Assembly IDs for the different strains are as follows: *E. coli* MS 196-1, GCA_000164555.1; *E. coli* MS 16-3, GCA_000164495.1; *E. coli* MS 119-7, GCA_000179155.1; *E. coli* MS 85-1, GCA_000179075.1; *E. coli* MS 198-1, GCA_000164195.1; *E. coli* MS 200-1, GCA_000164535.1. For **B-C**, data represent means ± standard deviations from three replicates. The assays in **B** and **C** were performed on different days with different media formulations.

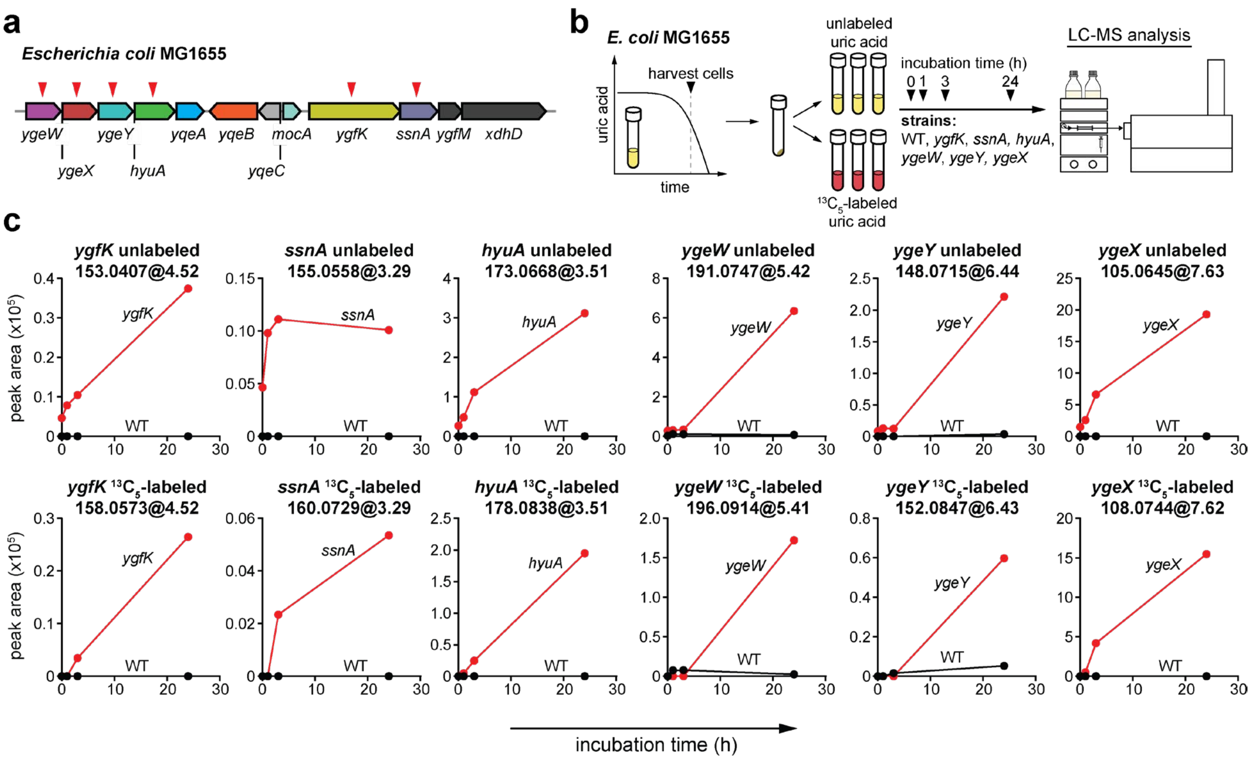

**Extended Data Figure 4. Isotope tracing in *E. coli* mutants.** (**A**) Gene cluster in *E. coli* showing mutants tested. (**B**) Overview of stable isotope tracing experiment. Because *E. coli* does not robustly consume uric acid during growth in rich media, we incubated concentrated cell suspensions with either labeled or unlabeled uric acid and monitored intermediates in supernatants over time by HILIC LC-MS. (**C**) Peak areas for *E. coli* mutant intermediates identified through our isotope tracing analysis of *C. sporogenes* mutants. Top panels indicate unlabeled cell suspensions and bottom panels are for $^{13}C$-labeled cell suspensions. For **C**, data are from a single replicate.

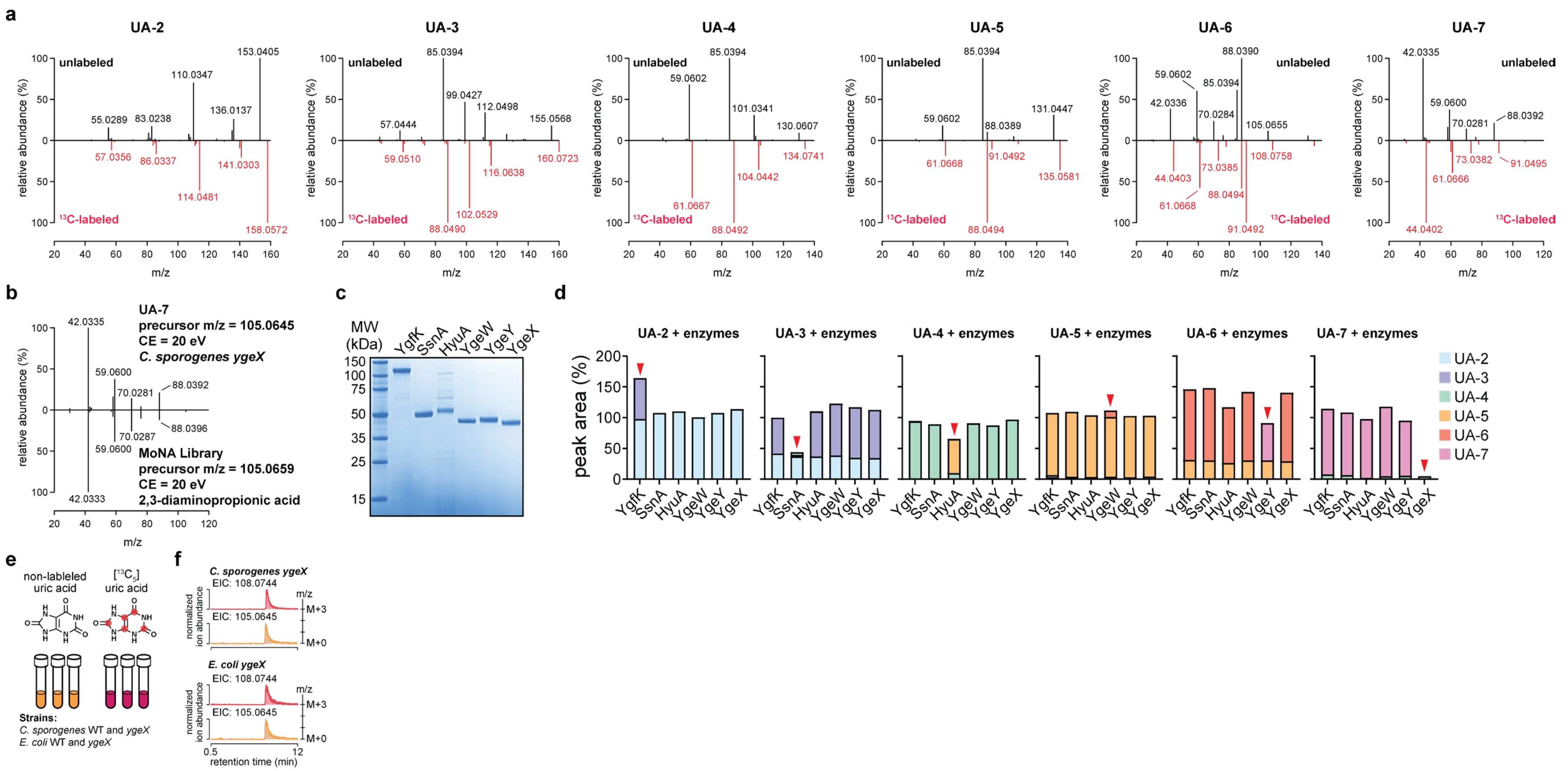

**Extended Data Figure 5. Mass fragmentation patterns for pathway intermediates and recombinant expression of enzymes.** (**A**) Targeted MS/MS was performed on intermediates using an Agilent LC-Q-TOF with collision energy of 20 eV. Top spectra are fragmentation patterns of unlabeled intermediates and bottom spectra are for labeled intermediates. (**B**) A precursor m/z and mass fragmentation pattern hit was found for UA-7 in the MoNA database. The top spectrum is from the unlabeled intermediate in *C. sporogenes* culture supernatants and the bottom spectrum is for 2,3-diaminopropanoic acid from the MoNA database. (**C**) SDS-PAGE gel showing purification of recombinant *E. coli* uric acid proteins. (**D**) Incubation of pathway intermediates (UA-2 through UA-7) with each of the recombinant *E. coli* proteins and analysis by HILIC LC-MS. Pathway intermediates were obtained by incubating concentrated cell suspensions of each *C. sporogenes* mutant with uric acid and filtering the supernatants (e.g., UA-2, *ygfK* mutant; UA-3, *ssnA* mutant; UA-4, *hyuA* mutant; UA-5, *ygeW* mutant; UA-6, *ygeY* mutant; UA-7, *ygeX* mutant). (**E**) Overview of stable isotope tracing for *C.*

*sporogenes* and *E. coli* wild-type (WT) and *ygeX* mutants. (**F**) Extracted ion chromatograms from HILIC LC-MS comparing m/z values for the molecular feature in *ygeX* culture supernatants with either [$^{13}C_5$]-labeled or non-labeled uric acid. For **D**, experiments were repeated twice, and representative data are shown.

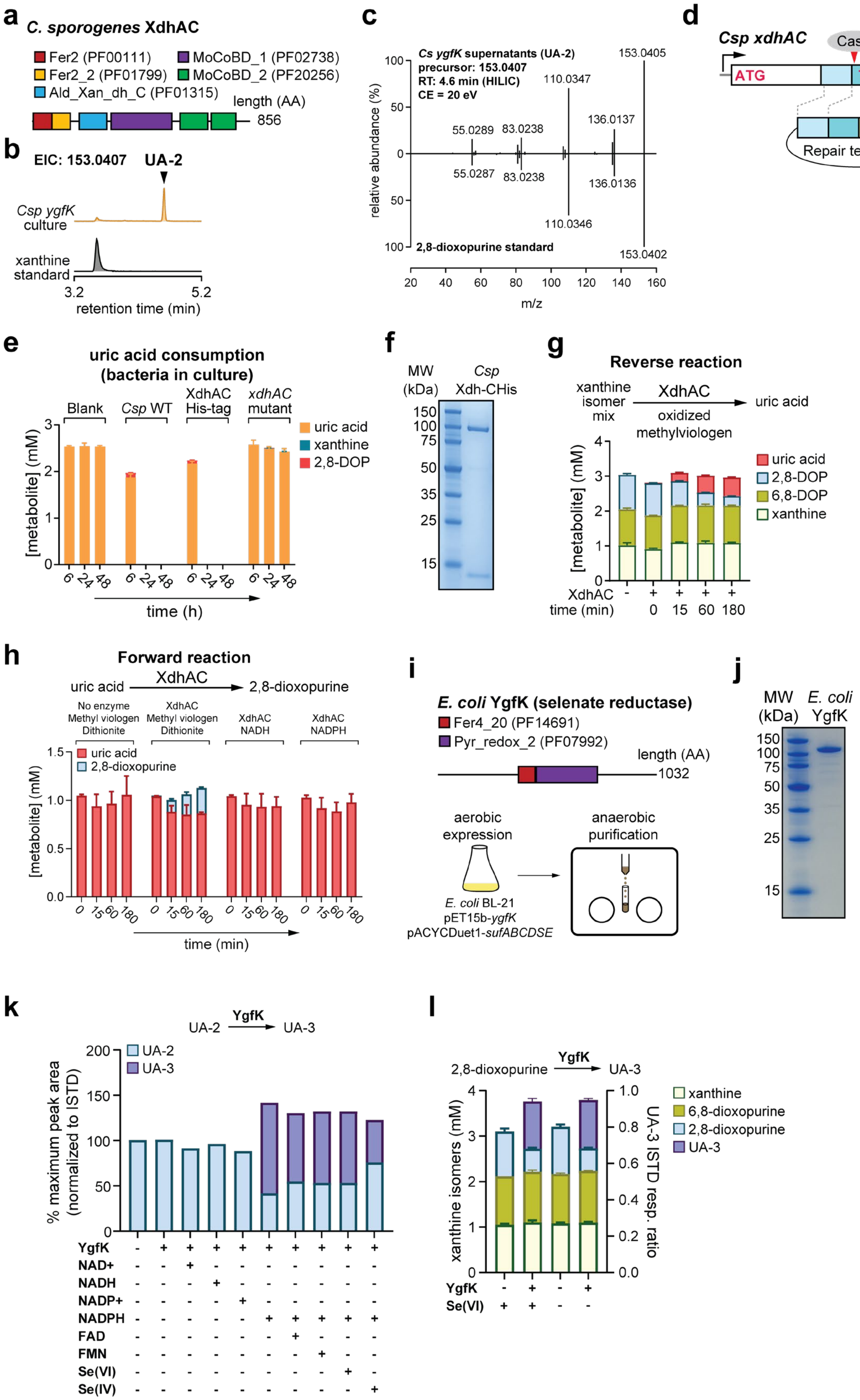
a
C. sporogenes XdhAC
Fer2 (PF00111)
MoCoBD_1 (PF02738)
Fer2_2 (PF01799)
MoCoBD_2 (PF20256)
Ald_Xan_dh_C (PF01315)
length (AA)
856
b
EIC: 153.0407
UA-2
Csp ygfK culture
xanthine standard
3.2
5.2
retention time (min)
c
Cs ygfK supernatants (UA-2)
precursor: 153.0407
RT: 4.6 min (HILIC)
CE = 20 eV
153.0405
110.0347
136.0137
55.0289
83.0238
55.0287
83.0238
136.0136
110.0346
153.0402
2,8-dioxopurine standard
relative abundance (%)
m/z
d
Csp xdhAC
Cas9
ATG
TAA
10X His
Repair template
e
uric acid consumption (bacteria in culture)
Blank
Csp WT
XdhAC His-tag
xdhAC mutant
uric acid
xanthine
2,8-DOP
[metabolite] (mM)
time (h)
f
MW (kDa)
Csp Xdh-CHis
g
Reverse reaction
xanthine isomer mix
XdhAC
oxidized methylviologen
uric acid
uric acid
2,8-DOP
6,8-DOP
xanthine
[metabolite] (mM)
XdhAC
time (min)
h
Forward reaction
uric acid
XdhAC
2,8-dioxopurine
No enzyme Methyl viologen Dithionite
XdhAC Methyl viologen Dithionite
XdhAC NADH
XdhAC NADPH
uric acid
2,8-dioxopurine
[metabolite] (mM)
time (min)
i
E. coli YgfK (selenate reductase)
Fer4_20 (PF14691)
Pyr_redox_2 (PF07992)
length (AA)
1032
aerobic expression
anaerobic purification
E. coli BL-21
pET15b-ygfK
pACYCDuet1-sufABCDSE
j
MW (kDa)
E. coli YgfK
k
UA-2
YgfK
UA-3
UA-2
UA-3
% maximum peak area (normalized to ISTD)
YgfK
NAD+
NADH
NADP+
NADPH
FAD
FMN
Se(VI)
Se(IV)
l
2,8-dioxopurine
YgfK
UA-3
xanthine
6,8-dioxopurine
2,8-dioxopurine
UA-3
xanthine isomers (mM)
UA-3 ISTD resp. ratio
YgfK
Se(VI)

**Extended Data Figure 6. Experimental support for the biochemical functions of YgfK (DOPOR) and XdhAC (DOPDH).** (**A**) Domain organization of *C. sporogenes* XdhAC. (**B**) Extracted ion chromatograms from for *C. sporogenes ygfK* culture supernatants and xanthine standard using HILIC LC-MS. (**C**) Targeted MS/MS was performed on UA-2 from *C. sporogenes ygfK* mutant culture supernatants and authentic 2,8-dioxopurine standard using an Agilent LC-Q-TOF with collision energy of 20 eV. (**D**) Schematic for introducing a C-terminal His-tag into *C. sporogenes xdhAC* using CRISPR. (**E**) Uric acid consumption by wild-type, His-tag engineered *xdhAC*, and mutant *xdhAC* strains of *C. sporogenes*. (**F**) SDS-PAGE of C-terminal His-tagged XdhAC purified natively from *C. sporogenes*. (**G**) Enzyme activity of purified XdhAC in the reverse reaction, catalyzing the oxidation of 2,8-dioxopurine to uric acid. (**H**) Testing different electron donors for *C. sporogenes* XdhAC. XdhAC was incubated with uric acid (1 mM) in the presence of dithionite (10 mM) and methylviologen (1 mM) or nicotinamides (10 mM each) and conversion of uric acid to 2,8-dioxopurine was followed by Poroshell positive LC-MS. (**I**) Domain organization of *E. coli* YgfK and overview of expression and anaerobic purification. (**J**) SDS-PAGE of anaerobically purified *E. coli* YgfK. (**K**) Co-factor dependence of *E. coli* YgfK using *C. sporogenes ygfK* mutant supernatants that contain UA-2. YgfK was incubated with various compounds (nicotinamides, 4 mM; flavins, 0.5 mM; selenium species, 2 mM) and UA-2 conversion to UA-3 after 1 hour was monitored by HILIC LC-MS. Note that the cell suspension supernatants used as the substrate likely also contains co-factors that cannot be removed. (**L**) Impact of selenate (Se(VI)) on *E. coli* YgfK activity using xanthine isomer mix. The three xanthine isomers (1 mM each) were dissolved in KPi buffer with NADPH (4 mM) and incubated with YgfK and sodium selenate (2 mM) as indicated. After 1 hour, xanthine isomers were quantified by Poroshell positive LC-MS and UA-3 was detected by HILIC LC-MS. For **K**, results are shown from a single replicate. For **E, G, H** and **L**, data represent means ± standard deviations from three replicates.

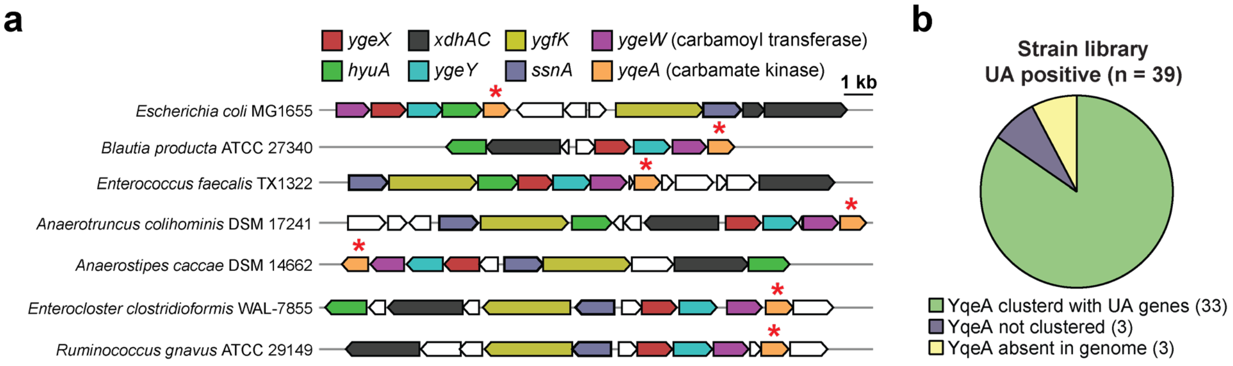

**Extended Data Figure 7. Carbamate kinase is linked to the uric acid gene cluster.** (**A**) Representative gene clusters from microbes in our strain library that have a carbamate kinase gene (*yqeA*), frequently adjacent to the carbamoyl transferase (*ygeW*). (**B**) Proportion of bacterial genomes in our strain library that have carbamate kinase (*yqeA*) clustered with uric acid genes, not clustered, or absent from genome.

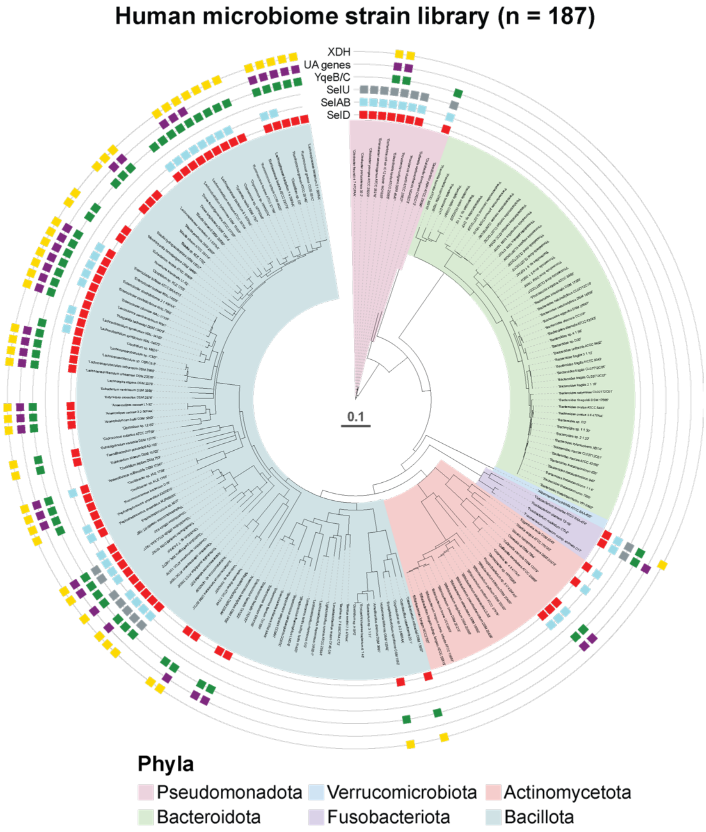

**Extended Data Figure 8. Distribution of selenium utilization traits and uric acid degrading genes across gut bacteria.** Phylogenetic tree of our human gut bacterial strain library showing presence of selenium utilization traits, uric acid degrading genes (UA genes) and xanthine dehydrogenase (XDH). SelD is selenophosphate synthase; SelAB predicts the selenocysteine decoding trait; SelU predicts the selenouridine trait; YqeB/C predicts the selenium-cofactor trait in molybdenum hydroxylases. Marker genes are listed on different tracks and solid filled boxes indicate presence of the corresponding marker genes.

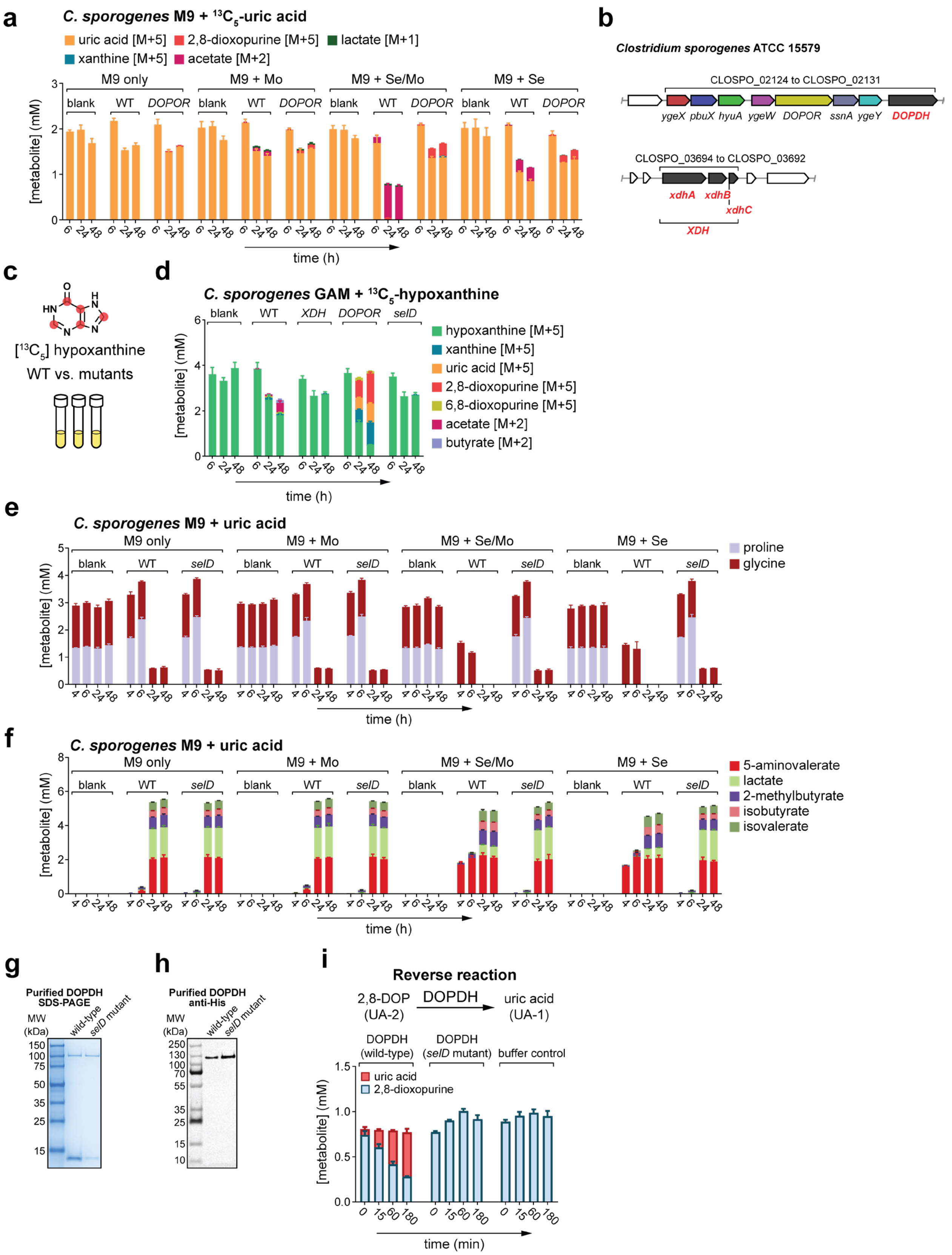
a
C. sporogenes M9 + 13C5-uric acid
uric acid [M+5]
2,8-dioxopurine [M+5]
lactate [M+1]
xanthine [M+5]
acetate [M+2]
M9 only
M9 + Mo
M9 + Se/Mo
M9 + Se
blank
WT
DOPOR
[metabolite] (mM)
time (h)
b
Clostridium sporogenes ATCC 15579
CLOSPO_02124 to CLOSPO_02131
ygeX pbuX hyuA ygeW DOPOR ssnA ygeY DOPDH
CLOSPO_03694 to CLOSPO_03692
xdhA xdhB xdhC
XDH
c
HN
N
H
O
[13C5] hypoxanthine
WT vs. mutants
d
C. sporogenes GAM + 13C5-hypoxanthine
blank
WT
XDH
DOPOR
selD
hypoxanthine [M+5]
xanthine [M+5]
uric acid [M+5]
2,8-dioxopurine [M+5]
6,8-dioxopurine [M+5]
acetate [M+2]
butyrate [M+2]
[metabolite] (mM)
time (h)
e
C. sporogenes M9 + uric acid
M9 only
M9 + Mo
M9 + Se/Mo
M9 + Se
blank
WT
selD
proline
glycine
[metabolite] (mM)
time (h)
f
C. sporogenes M9 + uric acid
M9 only
M9 + Mo
M9 + Se/Mo
M9 + Se
blank
WT
selD
5-aminovalerate
lactate
2-methylbutyrate
isobutyrate
isovalerate
[metabolite] (mM)
time (h)
g
Purified DOPDH
SDS-PAGE
MW (kDa)
wild-type
selD mutant
150
100
75
50
35
25
15
h
Purified DOPDH
anti-His
MW (kDa)
wild-type
selD mutant
250
130
100
70
55
35
25
15
10
i
Reverse reaction
2,8-DOP (UA-2)
DOPDH
uric acid (UA-1)
DOPDH (wild-type)
DOPDH (selD mutant)
buffer control
uric acid
2,8-dioxopurine
[metabolite] (mM)
time (min)

**Extended Data Figure 9. Data on XDH, DOPDH, and the *selD* mutant related to Figure 4.** (**A**) Stable isotope tracing in wild-type and mutant strains of *C. sporogenes* during growth in defined M9 media with or without the trace elements, molybdenum and selenium. (**B**) *C. sporogenes* genomic context for the 2,8-dioxopurine dehydrogenase (DOPDH) and xanthine dehydrogenase (XDH). (**C-D**) Stable isotope tracing during growth in Gifu anaerobic medium (GAM) with [$^{13}C_5$]-labeled hypoxanthine in wild-type and mutant strains of *C. sporogenes*. (**E-F**) Metabolic profiling of amino acids (**E**) and fatty acids (**F**) in wild-type and mutant *C. sporogenes* during growth in defined M9 media with or without the trace elements, molybdenum and selenium. Only metabolites that show significant differences between wild-type and *selD* mutant *C. sporogenes* are shown. All other metabolite data are presented in **Supplementary Table 7**. (**G**) SDS-PAGE and (**H**) western blot analysis of C-terminal His-tagged DOPDH purified natively from wild-type or *selD* mutant *C. sporogenes*. (**I**) Enzyme activity of purified DOPDH from wild-type or *selD* mutant in the reverse reaction, catalyzing the oxidation of 2,8-dioxopurine to uric acid. Metabolites were quantified by poroshell (**A, D, I**), NPH (**A, D, F**), or DNS (**E**) LC-MS and data represent means ± standard deviations from three replicates.

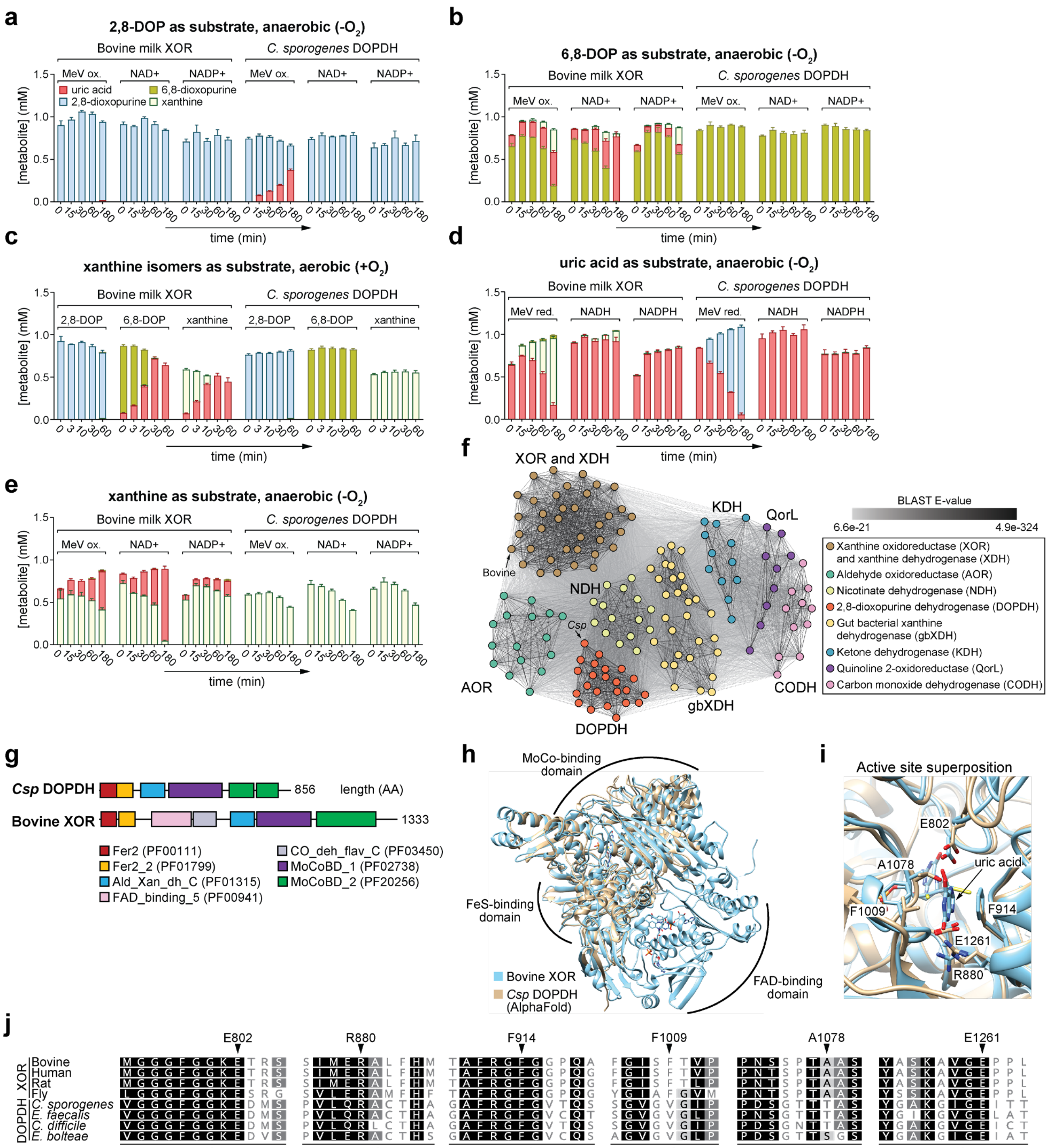

**Extended Data Figure 10. Biochemical and structural comparison of DOPDH and bovine XOR.** (**A-E**) Biochemical assays for DOPDH and bovine XOR with different redox cofactors and under anaerobic vs. aerobic conditions. Assays were initiated by the addition of substrate and aliquots taken at indicated timepoints were quenched in organic solvent (3:1 MeOH/ACN) and stored prior to analysis by porshell LC-MS. Note that bovine milk XOR has both xanthine dehydrogenase (XDH) activity and xanthine oxidase (XO) activity[3]. In (**C**), assays did not include supplemental redox co-factors, so oxygen present in these aerobic conditions represents the sole electron acceptor. (**F**)

Phylogenetic distribution of molybdenum-binding domains (PFAM PF02738 and PF20256) from well-characterized molybdenum hydroxylase families. All-against-all BLAST searches were performed on the corresponding sequences (**Supplementary Table 17**) using the CLANS web-utility in the MPI Bioinformatics Toolkit[4,5]. Two-dimensional force-directed layouts were generated with ~50,000 iterations. Dots represent individual protein sequences colored according to their protein family, and lines represent BLAST E-values with values indicated by line shading according to the legend. XOR, xanthine oxidoreductase; XDH, xanthine dehydrogenase (related to *Rhodobacter capsulatus*); AOR, aldehyde oxidoreductase; DOPDH, 2,8-dioxopurine dehydrogenase; NDH, nicotinate dehydrogenase; gbXDH, gut bacterial xanthine dehydrogenase (related to *Clostridium sporogenes*); KDH, ketone dehydrogenase; QorL, quinoline 2-oxidoreductase; CODH, carbon monoxide dehydrogenase. (**G**) Domain organization of *C. sporogenes* DOPDH and bovine XOR. (**H**) Structural alignment of bovine xanthine dehydrogenase and *C. sporogenes* DOPDH. The crystal structure for bovine XOR complexed with uric acid (PDB 3AMZ) and the AlphaFold model for *C. sporogenes* DOPDH (AF-J7T3J4-F1-v4.pdb) were aligned in UCSF Chimera using the MatchMaker feature. (**I**) Active site superposition of the structural alignment from **H**. Numbering is according to bovine XOR. (**J**) Amino acid sequence motifs containing active site residues. Amino acid sequences for molybdenum-binding domains from mammalian XORs and bacterial DOPDHs were aligned using MUSCLE. Active site residues corresponding to **I** are shown with arrows with numbering based on bovine XOR. For **A-E**, data represent means ± standard deviations from three replicates.

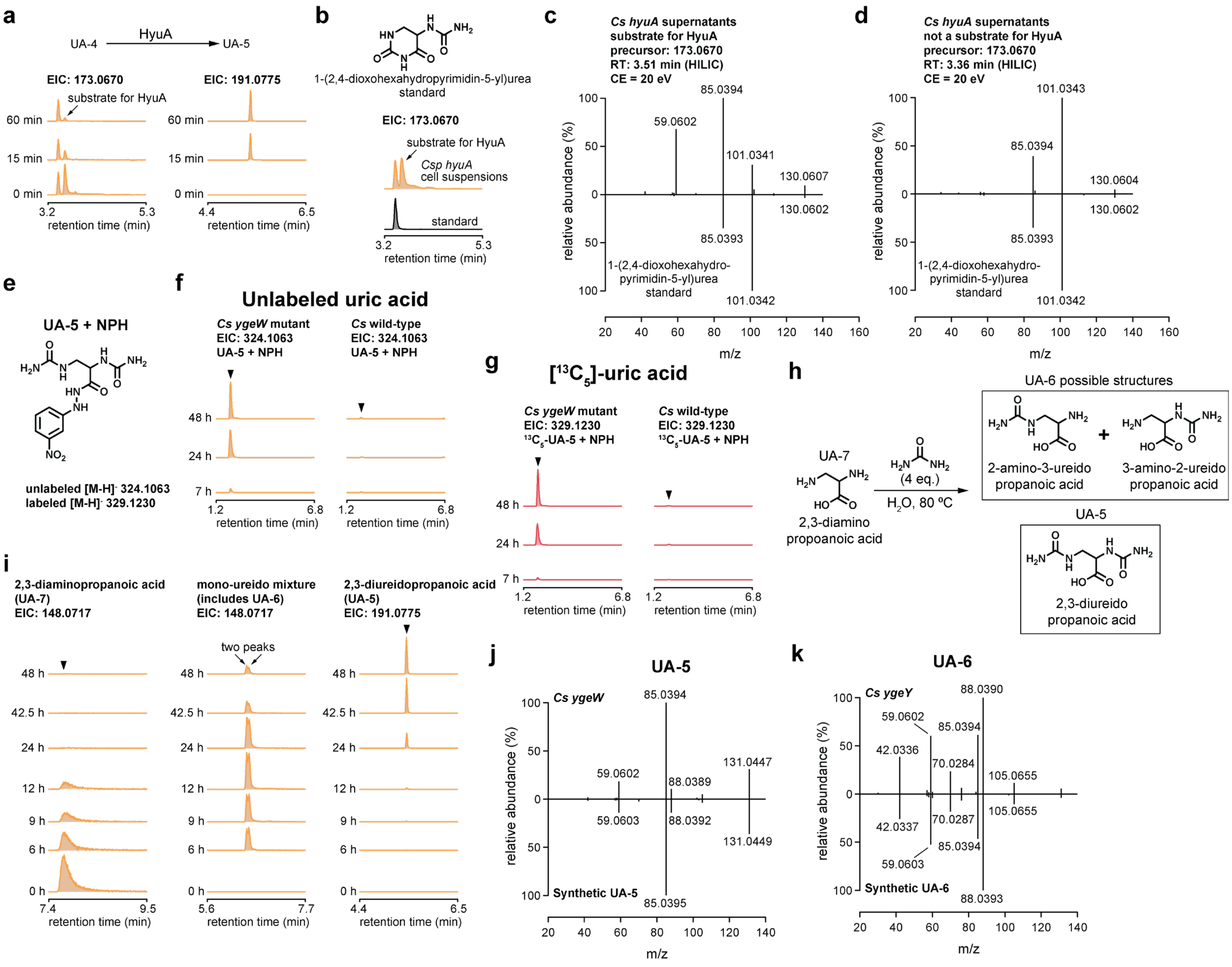
a
UA-4
HyuA
UA-5
EIC: 173.0670
EIC: 191.0775
substrate for HyuA
60 min
15 min
0 min
retention time (min)
b
1-(2,4-dioxohexahydropyrimidin-5-yl)urea standard
EIC: 173.0670
substrate for HyuA
Csp hyuA cell suspensions
standard
retention time (min)
c
Cs hyuA supernatants
substrate for HyuA
precursor: 173.0670
RT: 3.51 min (HILIC)
CE = 20 eV
85.0394
59.0602
101.0341
130.0607
130.0602
85.0393
101.0342
1-(2,4-dioxohexahydro-pyrimidin-5-yl)urea standard
relative abundance (%)
m/z
d
Cs hyuA supernatants
not a substrate for HyuA
precursor: 173.0670
RT: 3.36 min (HILIC)
CE = 20 eV
101.0343
85.0394
130.0604
130.0602
85.0393
101.0342
1-(2,4-dioxohexahydro-pyrimidin-5-yl)urea standard
relative abundance (%)
m/z
e
UA-5 + NPH
unlabeled [M-H]- 324.1063
labeled [M-H]- 329.1230
f
Unlabeled uric acid
Cs ygeW mutant
EIC: 324.1063
UA-5 + NPH
Cs wild-type
EIC: 324.1063
UA-5 + NPH
48 h
24 h
7 h
retention time (min)
g
[13C5]-uric acid
Cs ygeW mutant
EIC: 329.1230
13C5-UA-5 + NPH
Cs wild-type
EIC: 329.1230
13C5-UA-5 + NPH
retention time (min)
h
UA-7
2,3-diamino propoanoic acid
(4 eq.)
H2O, 80 °C
UA-6 possible structures
2-amino-3-ureido propanoic acid
3-amino-2-ureido propanoic acid
UA-5
2,3-diureido propanoic acid
i
2,3-diaminopropanoic acid (UA-7)
EIC: 148.0717
mono-ureido mixture (includes UA-6)
EIC: 148.0717
2,3-diureidopropanoic acid (UA-5)
EIC: 191.0775
two peaks
48 h
42.5 h
24 h
12 h
9 h
6 h
0 h
retention time (min)
j
UA-5
Cs ygeW
85.0394
59.0602
88.0389
131.0447
59.0603
88.0392
131.0449
85.0395
Synthetic UA-5
relative abundance (%)
m/z
k
UA-6
Cs ygeY
88.0390
59.0602
85.0394
42.0336
70.0284
105.0655
105.0655
42.0337
70.0287
85.0394
59.0603
88.0393
Synthetic UA-6
relative abundance (%)
m/z

**Supplementary Figure 1. Data supporting structural assignments of UA-4, UA-5, and UA-6.** (**A**) Extracted ion chromatograms from HILIC LC-MS for UA-4 and UA-5 in *C. sporogenes hyuA* mutant concentrated cell suspension supernatants incubated with *E. coli* HyuA. (**B**) Comparison of retention times by HILIC LC-MS between *C. sporogenes hyuA* mutant concentrated cell suspension supernatants and authentic 1-(2,4-dioxohexahydropyrimidin-5-yl)urea standard. (**C-D**) Tandem mass spectrometry comparing the mass spectra for the HyuA substrate (**C**) or the isomer that is not a substrate for HyuA (**D**). Targeted MS/MS was performed using an Agilent LC-Q-TOF with collision energy of 20 eV. (**E**) Predicted structure of UA-5 after derivatization with NPH, with expected mass-to-charge ratios for the unlabeled and [$^{13}C_5$]-labeled derivatives. (**F**) Extracted ion chromatograms from NPH LC-MS for unlabeled UA-5 NPH derivative in culture supernatants of the *C. sporogenes* wild-type and *ygeW* mutant strains during growth with unlabeled uric acid. (**G**) Extracted ion chromatograms from NPH LC-MS for $^{13}C$-labeled UA-5 NPH derivative in culture supernatants of the *C. sporogenes* wild-type and *ygeW* mutant strains during growth with [$^{13}C_5$]-labeled uric acid. (**H**) Overview of carbamation reaction. 2,3-Diaminopropanoic acid (DPA) was incubated with a 4-fold molar excess of urea in water at 80 °C. This yields a mixture of three compounds: two monoureido products which are candidates for UA-6 and the diureido product, likely to be UA-5. (**I**) Carbamation reaction progress monitored over time by HILIC LC-MS. Extracted ion chromatograms (EICs) are shown for the reactant (DPA) and the monoureido and diureido products. Note that two closely eluting peaks were detected for the monoureido EIC, suggesting a mixture of two isomers. (**J**) Targeted MS/MS was performed on UA-5 from *C. sporogenes ygeW* mutant culture supernatants and synthetic diureido compound from the carbamation reaction. (**K**) Targeted MS/MS was performed on UA-6 from *C. sporogenes ygeY* mutant culture supernatants and synthetic monoureido compound from the carbamation reaction. Note that we were unable to chromatographically separate the two monoureido compounds, so the MS/MS spectrum of the synthetic compound likely represents a mixture of the two isomers.

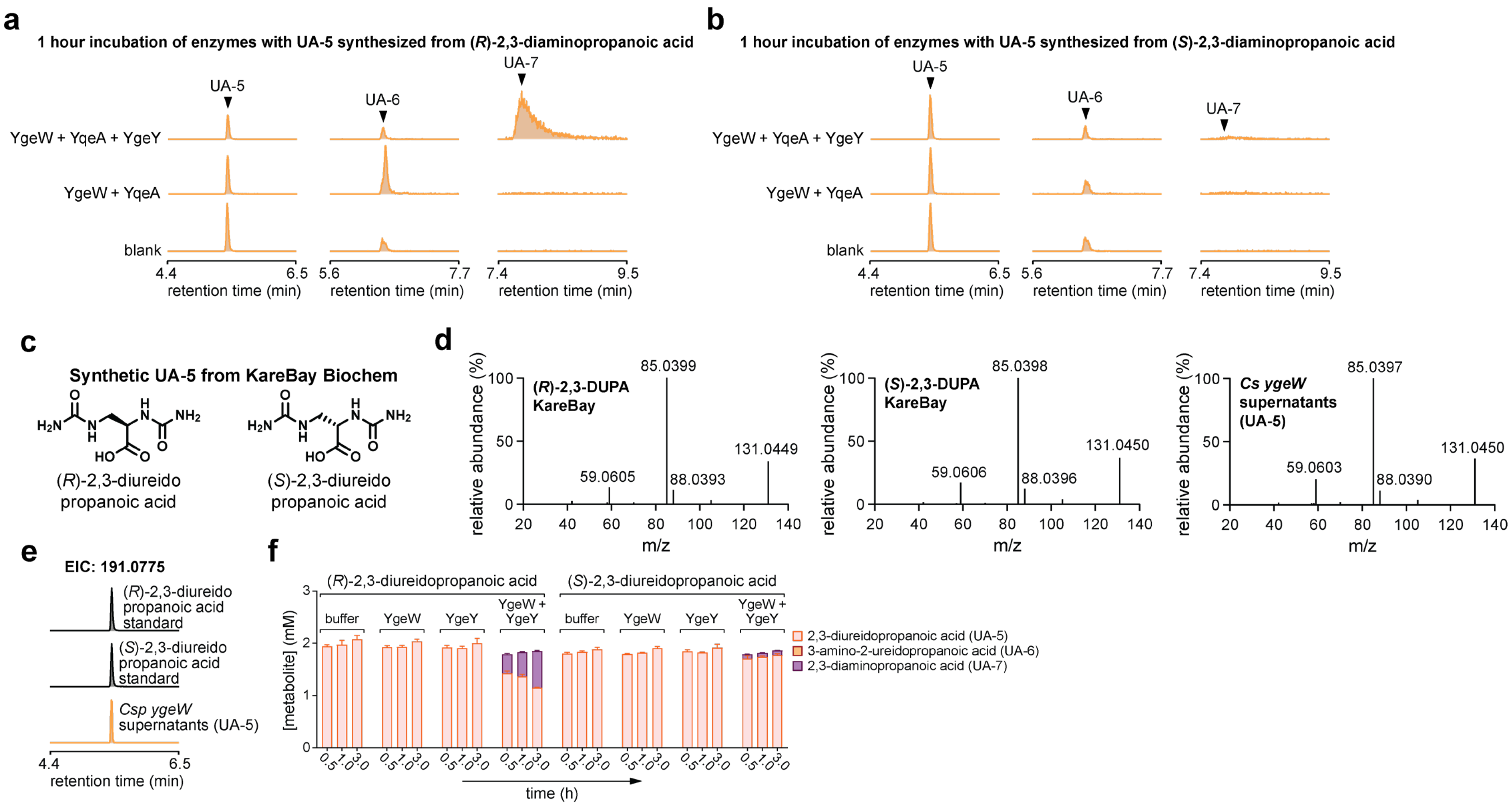

**Supplementary Figure 2. Data supporting structural assignment of UA-5.** (**A-B**) UA-5 was synthesized by carbamation of either (*R*)-2,3-diaminopropanoic acid (**A**) or (*S*)-2,3-diaminopropanoic acid (**B**). The 48-hour reaction product, where UA-5 was the dominant product (**Supplementary Figure 1i**) was incubated with the indicated enzymes for 1 hour and UA-5, UA-6, and UA-7 were detected by HILIC LC-MS. (**C**) Structures of two enantiomers of UA-5 synthesized by KareBay Biochem (datasheets in **Supplementary Data File 2**). (**D**) Targeted MS/MS was performed on authentic (*R*)-2,3-diureidopropanoic acid ((*R*)-2,3-DUPA), (*S*)-2,3-diureidopropanoic acid ((*S*)-2,3-DUPA) and UA-5 from *C. sporogenes ygeW* mutant culture supernatants. Spectra were obtained using an Agilent LC-Q-TOF with precursor mass selection of 191.0775 and collision energy of 20 eV. (**E**) Extracted ion chromatograms by HILIC LC-MS for authentic (*R*)-2,3-diureidopropanoic acid, (*S*)-2,3-diureidopropanoic acid, and UA-5 from *C. sporogenes ygeW* mutant culture supernatants. (**F**) YgeW and YgeY enzymes were incubated with (*R*)-2,3-diureidopropanoic acid or (*S*)-2,3-diureidopropanoic acid and UA-5, UA-6 were quantified by HILIC and UA-7 by DNS LC-MS. Data represent means ± standard deviations from three replicates.

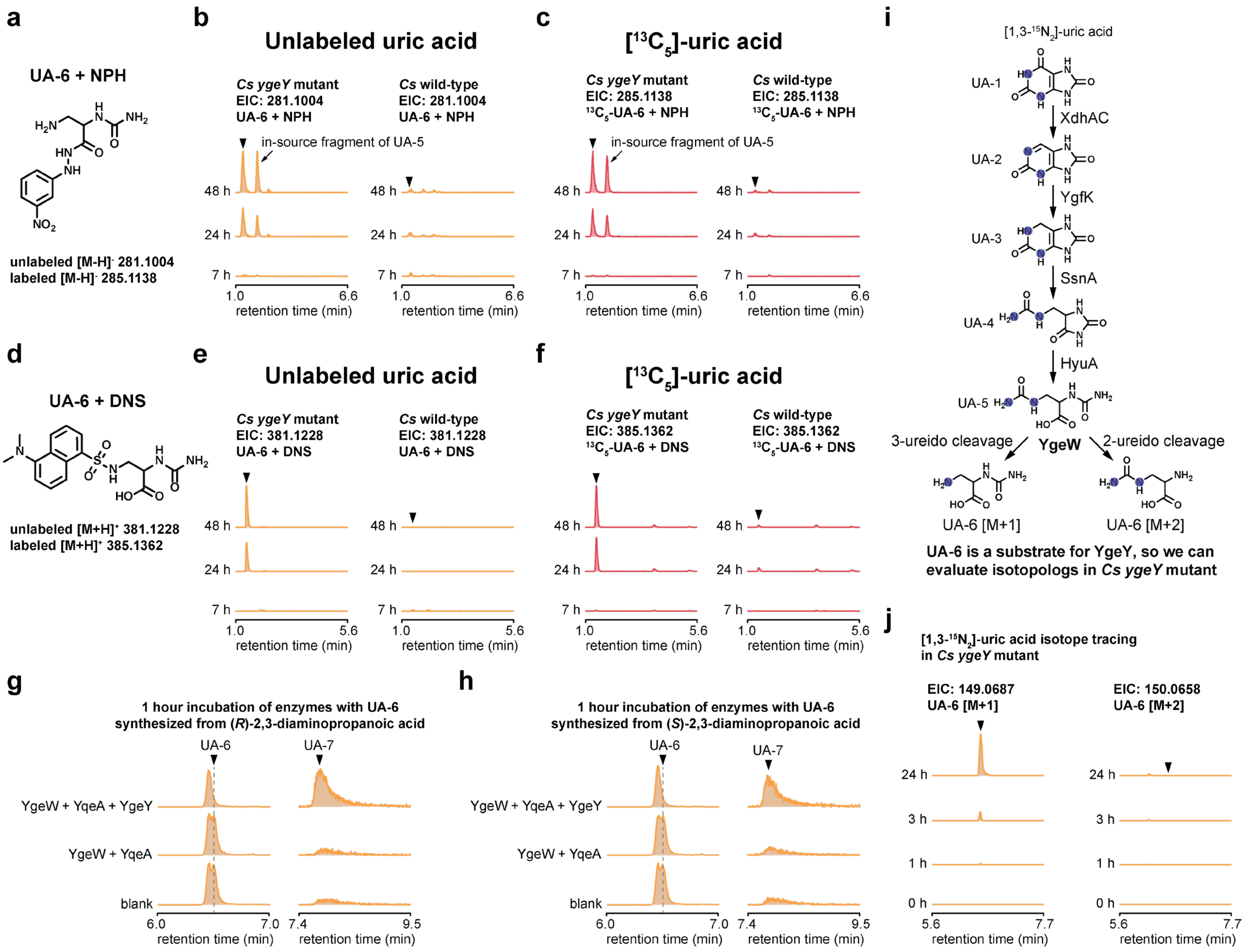

a
UA-6 + NPH
unlabeled [M-H]- 281.1004
labeled [M-H]- 285.1138
b
Unlabeled uric acid
Cs ygeY mutant
EIC: 281.1004
UA-6 + NPH
Cs wild-type
EIC: 281.1004
UA-6 + NPH
in-source fragment of UA-5
retention time (min)
c
[13C5]-uric acid
Cs ygeY mutant
EIC: 285.1138
13C5-UA-6 + NPH
Cs wild-type
EIC: 285.1138
13C5-UA-6 + NPH
in-source fragment of UA-5
d
UA-6 + DNS
unlabeled [M+H]+ 381.1228
labeled [M+H]+ 385.1362
e
Unlabeled uric acid
Cs ygeY mutant
EIC: 381.1228
UA-6 + DNS
Cs wild-type
EIC: 381.1228
UA-6 + DNS
f
[13C5]-uric acid
Cs ygeY mutant
EIC: 385.1362
13C5-UA-6 + DNS
Cs wild-type
EIC: 385.1362
13C5-UA-6 + DNS
i
[1,3-15N2]-uric acid
UA-1
XdhAC
UA-2
YgfK
UA-3
SsnA
UA-4
HyuA
UA-5
3-ureido cleavage
YgeW
2-ureido cleavage
UA-6 [M+1]
UA-6 [M+2]
UA-6 is a substrate for YgeY, so we can evaluate isotopologs in Cs ygeY mutant
g
1 hour incubation of enzymes with UA-6 synthesized from (R)-2,3-diaminopropanoic acid
UA-6
UA-7
YgeW + YqeA + YgeY
YgeW + YqeA
blank
h
1 hour incubation of enzymes with UA-6 synthesized from (S)-2,3-diaminopropanoic acid
j
[1,3-15N2]-uric acid isotope tracing in Cs ygeY mutant
EIC: 149.0687
UA-6 [M+1]
EIC: 150.0658
UA-6 [M+2]
retention time (min)

389

**Supplementary Figure 3. Data supporting structural assignment of UA-6.** (**A**) Predicted structure of UA-6 after derivatization with NPH, with expected mass-to-charge ratios for the unlabeled and $^{13}C$-labeled derivatives. (**B**) Extracted ion chromatograms from NPH LC-MS for unlabeled UA-6 NPH derivative in culture supernatants of the *C. sporogenes* wild-type and *ygeY* mutant strains during growth with unlabeled uric acid. (**C**) Extracted ion chromatograms from NPH LC-MS for $^{13}C$-labeled UA-6 NPH derivative in culture supernatants of the *C. sporogenes* wild-type and *ygeY* mutant strains during growth with [$^{13}C_5$]-labeled uric acid. Note that we identified a second peak as an in-source fragment of UA-5 which also accumulates in the supernatants of the *C. sporogenes ygeY* mutant (**Figure 1b**). (**D**) Predicted structure of UA-6 after derivatization with DNS, with expected mass-to-charge ratios for the unlabeled and $^{13}C$-labeled derivatives. (**E**) Extracted ion chromatograms from DNS LC-MS for unlabeled UA-6 DNS derivative in culture supernatants of the *C. sporogenes* wild-type and *ygeY* mutant strains during growth with unlabeled uric acid. (**F**) Extracted ion chromatograms from DNS LC-MS for $^{13}C$-labeled UA-6 DNS derivative in culture supernatants of the *C. sporogenes* wild-type and *ygeY* mutant strains during growth with [$^{13}C_5$]-labeled uric acid. (**G-H**) UA-6 was synthesized by carbamation of either (*R*)-2,3-diaminopropanoic acid (**G**) or (*S*)-2,3-diaminopropanoic acid (**H**). The 12-hour reaction product, where UA-6 was the dominant product (**Supplementary Figure 1i**) was incubated with the indicated enzymes for 1 hour and UA-5, UA-6, and UA-7 were detected by HILIC LC-MS. Note that the second (right-most) peak for the UA-6 synthetic product disappeared when YgeY was added to the reactions as indicated by the dashed line. (**I**) Overview of the uric acid degradation pathway showing the positions of the two labeled nitrogen atoms. If YgeW cleaves the 3-ureido group, it will result in a singly labeled product, whereas if it cleaves the 2-ureido group, the product will have two labels. (**J**) Extracted ion chromatograms from HILIC LC-MS for the singly or doubly labeled UA-6 product that accumulated in concentrated cell suspension supernatants of the *C. sporogenes ygeY* mutant.

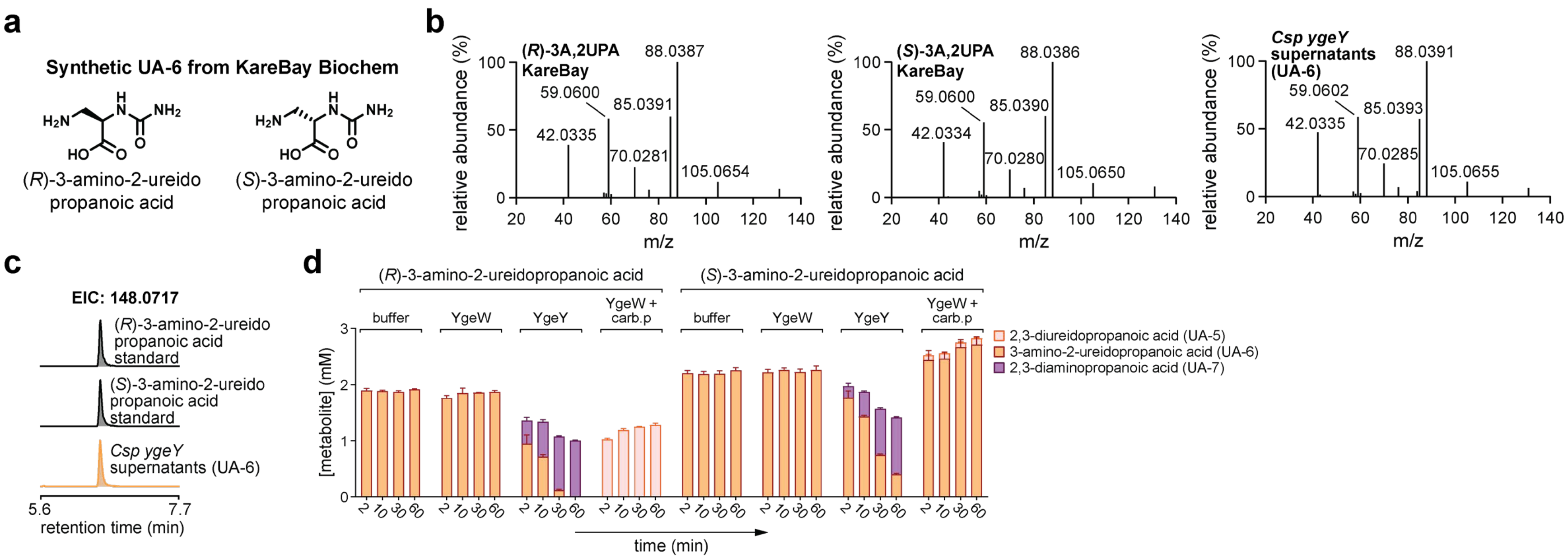

**Supplementary Figure 4. Data supporting structural assignment of UA-6.** (**A**) Structures of two enantiomers of UA-6 synthesized by KareBay Biochem (datasheets in **Supplementary Data File 3**). (**B**) Targeted MS/MS was performed on authentic (*R*)-3-amino-2-ureidopropanoic acid ((*R*)-3A,2UPA), (*S*)-3-amino-2-ureidopropanoic acid ((*S*)-3A,2UPA), and UA-6 from *C. sporogenes ygeY* mutant culture supernatants. Spectra were obtained using an Agilent LC-Q-TOF with precursor mass selection of 191.0775 and collision energy of 20 eV. (**C**) Extracted ion chromatograms by HILIC LC-MS for authentic (*R*)-3-amino-2-ureidopropanoic acid, (*S*)-3-amino-2-ureidopropanoic acid, and UA-6 from *C. sporogenes ygeY* mutant culture supernatants. (**D**) YgeW and YgeY enzymes were incubated with (*R*)-3-amino-2-ureidopropanoic acid or (*S*)-3-amino-2-ureidopropanoic acid and UA-5, UA-6 were quantified by HILIC and UA-7 by DNS LC-MS. Carbamoyl phosphate (carb.p) was added to YgeW enzyme incubations to drive synthesis of UA-5 in the reverse reaction. Data represent means ± standard deviations from three replicates.

*Teklad Custom Diet*

# TD.220052

**3% OA, 2% UA Diet (VI, Fiber, RS)**

#### Formula

| Formula | g/Kg |
|---|---|
| Casein | 218.0 |
| L-Cystine | 3.0 |
| Sucrose | 228.78 |
| Corn Starch | 150.0 |
| Maltodextrin | 80.0 |
| High Amylose Corn Starch | 35.0 |
| Inulin | 35.0 |
| Pectin | 35.0 |
| Soybean Oil | 80.0 |
| Cellulose | 25.0 |
| Mineral Mix, AIN-93M-MX (94049) | 35.0 |
| Calcium Phosphate, monobasic, monohydrate | 2.0 |
| Calcium Carbonate | 1.0 |
| Vitamin Mix, AIN-93-VX (94047) | 19.5 |
| Choline Bitartrate | 2.7 |
| TBHQ, antioxidant | 0.02 |
| Oxonic Acid, potassium salt, customer supplied | 30.0 |
| Uric Acid, customer supplied | 20.0 |

#### Footnote

Modification of TD.210629 to include 3% oxonic acid and 2% uric acid. Contains 3.5% each high amylose corn starch, pectin, and inulin. Vitamins are increased for irradiation.

#### Selected Nutrient Information[1]

| | % by weight | % kcal from |
|---|---|---|
| **Protein** | 19.3 | 21.9 |
| **Carbohydrate** | 50.1 | 57.0 |
| **Fat** | 8.2 | 21.0 |
| **Kcal/g** | **3.5** | |

[1] Values are calculated from ingredient analysis or manufacturer data

#### Speak With A Nutritionist

+ **(800) 483·5523**
+ ****

*Teklad diets are designed & manufactured for research purposes only.*

#### Key Features

+ **Purified Diet**
+ **Resistant Starch, Pectin, and Inulin**
+ **Oxonic Acid and Uric Acid**
+ **Suitable For Irradiation**

#### Key Planning Information

+ *Products are made fresh to order*
+ Store product at 4°C or lower
+ Use within 6 months (applicable to most diets)
+ Box labeled with product name, manufacturing date, and lot number
+ Replace diet at minimum once per week
  *More frequent replacement may be advised*
+ Lead time:
  · 2 weeks non-irradiated
  · 4 weeks irradiated

#### Product Specific Information

+ 1/2" Pellet or Powder (free flowing)
+ Minimum order 3 Kg
+ Irradiation available upon request

#### Options (fees will apply)

+ Rush order (pending availability)
+ Irradiation (see Product Specific Information)
+ Vacuum packaging (1 and 2 Kg)

#### Contact Us

*Obtain pricing · Check order status*

+ ****
+ **(800) 483·5523**

#### International Inquiry *(outside USA or Canada)*

+ ****

#### Place Your Order (USA & Canada)

*Please Choose One*

+ **www.envigo.com/teklad-orders**
+ ****
+ **(800) 483·5523**

#N/A

© 2015 Envigo

01/18/22

### Certificate of Analysis

**Compound Name:** (R)-2,3-diureidopropanoic acid

**Physical Description** White powder

**CAS** N/A

**Chemical Structure**

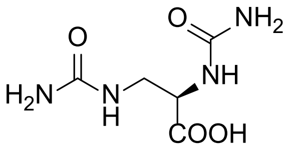

**Molecular Weight** 190

**Chemical Formula** C5H10N4O4

**Purity** 95 % (214nm)

**Lot No.** KB240712-1

#### Storage Conditions

lndividual variation in chemical stability profiles, do occur and for this reason we recommend adherence to the minimum recommended storage conditions. Because most compounds are custom made and shipped upon completion, long-term stability data is not available in most cases. If data is available, a specific recommendation will be provided below. Chemist cannot provide a guarantee of the long-term chemical stability of any compound.

Mlnimum Recommended Storage Conditions:

- Samples should be stored in an air tight vial
- Samples should be stored at≤ 0° Celsius when not in use for a prolonged period of time
- Samples should be stored in the dark or in an amber vial.

#### Caution

This information is provided as an indication of the quality of the underlying material when examined by a specific technique. The reported values are subject to normal experimental error and should be treated as estimates. The absence of undetected impurities cannot be guaranteed by this or any other general approach and this certificate does not certify the absence of such substances in the sample.

#### Intended Use

This product is intended for investigational use only and should not be used in humans. It is pharmaceutically unrefined, may contain uncharacterized toxic impurities, and is not intended for use in humans. Responsibility for its use and compliance with all federal laws rests solely with the purchaser.

### HPLC (214nm)

Sample Information

| | |
|---|---|
| Sample Name | : ENBB240469-001-A1 |
| Sample ID | : ENBB240469-001-A1 |
| Tray# | : 1 |
| Vail# | : 20 |
| Injection Volume | : 2 uL |
| Data Filename | : ENBB240469-001-A1_1.lcd |
| Method Filename | : AB-0-60-100-13MIN-35C.lcm |
| Date Acquired | : 2024/7/2 14:16:50 |
| Description | : Method: 0-60 |

Column: Gemini C18 4.6*150mm 5um
Mobile phase:
H2O(0.05%TFA)-ACN(0.05%TFA)
ACN from 0% to 60% over 7.5 minutes
7.5-8min , ACN from 60% to 100%, hold 2 minutes
Oven: 35C
Flow rate: 1.2mL/min
Instrument: SHIMADZU LC-2050C

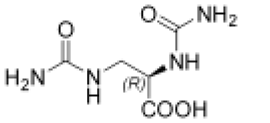

**KB13-R**

Chemical Formula: $C_5H_{10}N_4O_4$

Molecular Weight: 190.16

<Chromatogram>

ENBB240469-001-A1 D:\...\DATA\2024\202407\2\LC-7\ENBB240469-001-A1_1.lcd

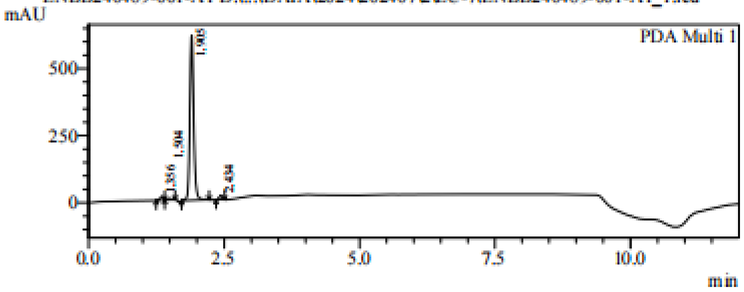

1 PDA Multi 1 / 214nm,4nm

<Result>

PeakTable

PDA Ch1 214nm

| Peak# | Ret. Time | Area | Height | Area % |
|---|---|---|---|---|
| 1 | 1.356 | 65581 | 14334 | 2.049 |
| 2 | 1.504 | 26859 | 4953 | 0.839 |
| 3 | 1.905 | 3041407 | 615252 | 95.010 |
| 4 | 2.434 | 67290 | 16576 | 2.102 |
| Total | | 3201137 | 651115 | 100.000 |

### HPLC (254nm)

Sample Information

Sample Name : ENBB240469-001-A1
Sample ID : ENBB240469-001-A1
Tray# : 1
Vail# : 20
Injection Volume : 2 uL
Data Filename : ENBB240469-001-A1_1.lcd
Method Filename : AB-0-60-100-13MIN-35C.lcm
Date Acquired : 2024/7/2 14:16:50
Description : Method: 0-60
Column: Gemini C18 4.6*150mm 5um
Mobile phase:
H2O(0.05%TFA)-ACN(0.05%TFA)
ACN from 0% to 60% over 7.5 minutes
7.5-8min , ACN from 60% to 100%, hold 2 minutes
Oven: 35C
Flow rate: 1.2mL/min
Instrument: SHIMADZU LC-2050C

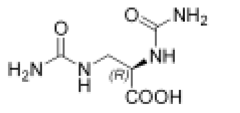

**KB13-R**

Chemical Formula: $C_5H_{10}N_4O_4$

Molecular Weight: 190.16

<Chromatogram>

ENBB240469-001-A1 D:\...\DATA\2024\20240702\LC-7\ENBB240469-001-A1_1.lcd

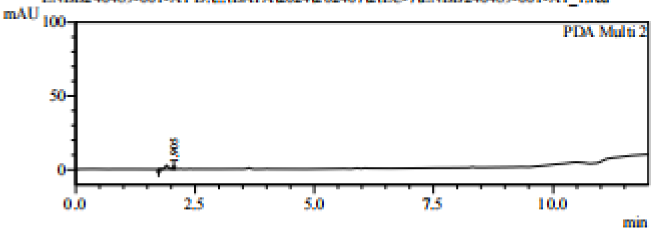

1 PDA Multi 2 / 254nm,4nm

<Result>

PeakTable

PDA Ch2 254nm

| Peak# | Ret. Time | Area | Height | Area % |
|---|---|---|---|---|
| 1 | 1.905 | 12190 | 2467 | 100.000 |
| Total | | 12190 | 2467 | 100.000 |

### LCMS

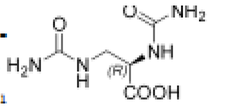

**KB13-R**

Chemical Formula: $C_5H_{10}N_4O_4$

Molecular Weight: 190.16

```
Acq. Operator   : ADMIN
Acq. Instrument : LCMS-B                              Location : P1-B-06
Injection Date  : 7/2/2024 9:07:32 AM                      Inj :    1
                                                    Inj Volume : 1.000 µl
Acq. Method     : D:\CHEMSTATION\1\METHODS\P50-600.M
Last changed    : 6/29/2024 4:38:26 PM by ADMIN
Analysis Method : C:\CHEM32\1\METHODS\DEF_LC.M
Last changed    : 11/20/2006 6:14:44 PM
Additional Info : Peak(s) manually integrated
```

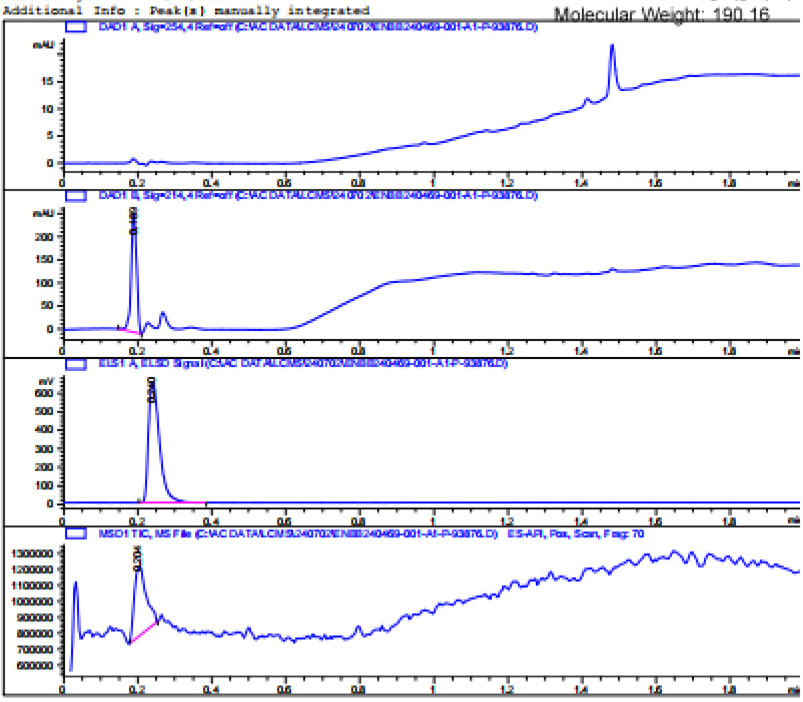

```
ata File C:\AC DATA\LCMS\240702\ENBB240469-001-A1-P-93876.D
ample Name: ENBB240469-001-A1-P

=====================================================================
                         Area Percent Report
=====================================================================

Sorted By             :      Signal
Multiplier:                  :      1.0000
Dilution:                    :      1.0000
Use Multiplier & Dilution Factor with ISTDs

Signal 1: DAD1 A, Sig=254,4 Ref=off

Signal 2: DAD1 B, Sig=214,4 Ref=off

Peak RetTime Type  Width     Area      Height     Area
  #   [min]        [min]   [mAU*s]     [mAU]        %
----|-------|----|-------|----------|----------|--------|
   1   0.189 BB    0.0165  264.69244  256.61115 100.0000

Totals :                   264.69244  256.61115

Signal 3: ELS1 A, ELSD Signal

Peak RetTime Type  Width     Area      Height     Area
  #   [min]        [min]   [mV*s]      [mV]         %
----|-------|----|-------|----------|----------|--------|
   1   0.240 BB    0.0327 1348.84143  650.74426 100.0000

Totals :                  1348.84143  650.74426

Signal 4: MSD1 TIC, MS File

Peak RetTime Type  Width     Area      Height     Area
  #   [min]        [min]                            %
----|-------|----|-------|----------|----------|--------|
   1   0.204 BB    0.0327 9.74948e5  4.38686e5  100.0000

Totals :                  9.74948e5  4.38686e5

=====================================================================
                         *** End of Report ***
```

MS Spectrum

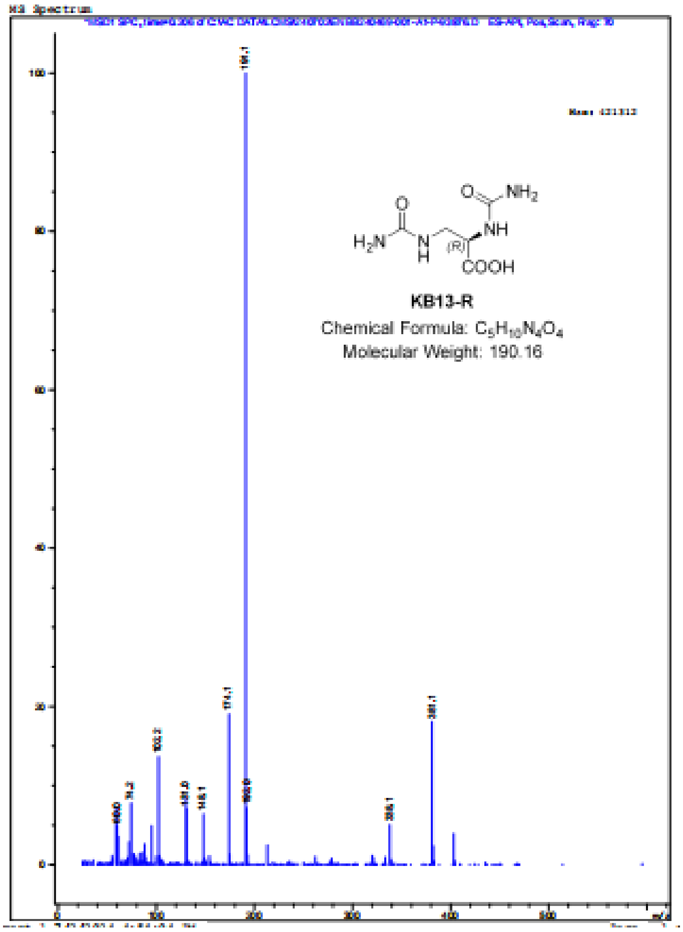

KB13-R
Chemical Formula: $C_5H_{10}N_4O_4$
Molecular Weight: 190.16
191.1
174.1
381.1
103.2

### HNMR

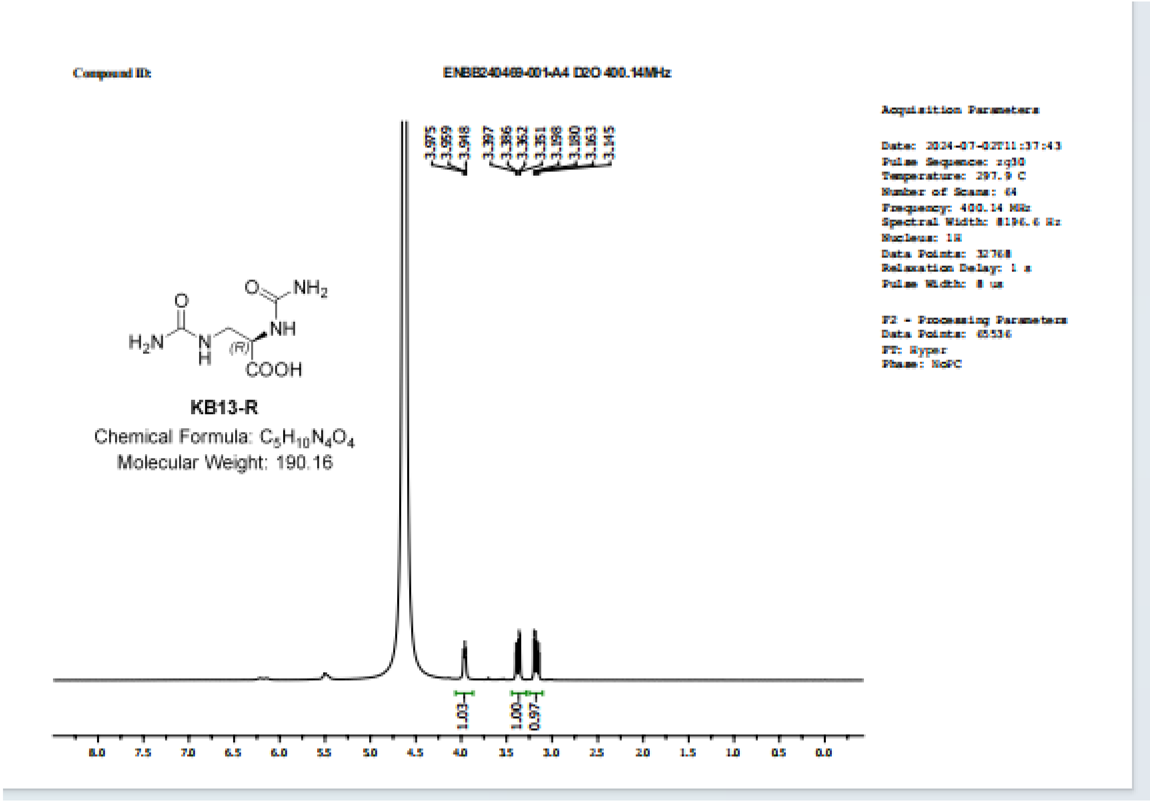

Compound ID:
ENBB240469-001-A4 D2O 400.14MHz
3.975
3.959
3.948
3.397
3.386
3.362
3.351
3.198
3.180
3.163
3.145
KB13-R
Chemical Formula: C5H10N4O4
Molecular Weight: 190.16
1.03
1.00
0.97
8.0 7.5 7.0 6.5 6.0 5.5 5.0 4.5 4.0 3.5 3.0 2.5 2.0 1.5 1.0 0.5 0.0
Acquisition Parameters
Date: 2024-07-02T11:37:43
Pulse Sequence: zg30
Temperature: 297.9 C
Number of Scans: 64
Frequency: 400.14 MHz
Spectral Width: 8196.6 Hz
Nucleus: 1H
Data Points: 32768
Relaxation Delay: 1 s
Pulse Width: 8 us
F2 - Processing Parameters
Data Points: 65536
FT: Hyper
Phase: NoPC

### Certificate of Analysis

**Compound Name:** (S)-2,3-diureidopropanoic acid

**Physical Description** White powder

**CAS** N/A

**Chemical Structure**

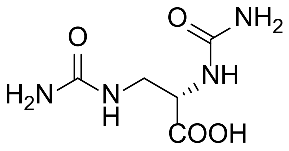

**Molecular Weight** 190

**Chemical Formula** C5H10N4O4

**Purity** 100 % (214nm)

**Lot No.** KB240712-2

#### Storage Conditions

lndividual variation in chemical stability profiles, do occur and for this reason we recommend adherence to the minimum recommended storage conditions. Because most compounds are custom made and shipped upon completion, long-term stability data is not available in most cases. If data is available, a specific recommendation will be provided below. Chemist cannot provide a guarantee of the long-term chemical stability of any compound.

Mlnimum Recommended Storage Conditions:

- Samples should be stored in an air tight vial
- Samples should be stored at⩽ 0° Celsius when not in use for a prolonged period of time
- Samples should be stored in the dark or in an amber vial.

#### Caution

This information is provided as an indication of the quality of the underlying material when examined by a specific technique. The reported values are subject to normal experimental error and should be treated as estimates. The absence of undetected impurities cannot be guaranteed by this or any other general approach and this certificate does not certify the absence of such substances in the sample.

#### Intended Use

This product is intended for investigational use only and should not be used in humans. It is pharmaceutically unrefined, may contain uncharacterized toxic impurities, and is not intended for use in humans. Responsibility for its use and compliance with all federal laws rests solely with the purchaser.

### HPLC (214nm)

Sample Information

Sample Name : ENBB240469-002-A1-HPLC
Sample ID : ENBB240469-002-A1-HPLC
Tray# : 1
Vail# : 21
Injection Volume : 3 uL
Data Filename : ENBB240469-002-A1-HPLC_1.lcd
Method Filename : AB-0-60-100-13MIN-35C.lcm
Date Acquired : 7/9/2024 1:47:22 PM
Description : Method: 0-60
Column: Gemini C18 4.6*150mm 5um
Mobile phase:
H2O(0.05%TFA)-ACN(0.05%TFA)
ACN from 0% to 60% over 7.5 minutes
7.5-8min, ACN from 60% to 100%, hold 2 minutes
Oven: 35C
Flow rate: 1.2mL/min
Instrument: SHIMADZU LC-2030C

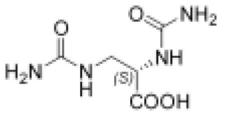

KB13-S
Chemical Formula: $C_5H_{10}N_4O_4$
Molecular Weight: 190.16

<Chromatogram>

ENBB240469-002-A1-HPLC D:\acdata\2024\07\09\LC-6\ENBB240469-002-A1-HPLC_1.lcd

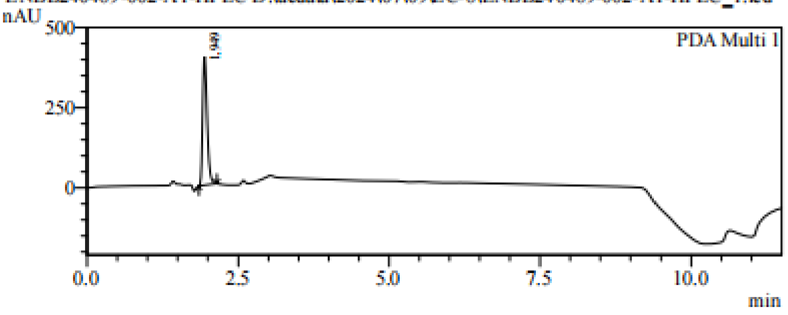

PDA Multi 1 / 214nm,4nm

<Result>

PeakTable

PDA Ch1 214nm

| Peak# | Ret. Time | Area | Height | Area % |
|---|---|---|---|---|
| 1 | 1.949 | 2057953 | 399711 | 100.000 |
| Total | | 2057953 | 399711 | 100.000 |

### HPLC (254nm)

Sample Information

Sample Name : ENBB240469-002-A1-HPLC
Sample ID : ENBB240469-002-A1-HPLC
Tray# : 1
Vail# : 21
Injection Volume : 3 uL
Data Filename : ENBB240469-002-A1-HPLC_1.lcd
Method Filename : AB-0-60-100-13MIN-35C.lcm
Date Acquired : 7/9/2024 1:47:22 PM
Description : Method: 0-60
Column: Gemini C18 4.6*150mm 5um
Mobile phase:
H2O(0.05%TFA)-ACN(0.05%TFA)
ACN from 0% to 60% over 7.5 minutes
7.5-8min , ACN from 60% to 100%, hold 2 minutes
Oven: 35C
Flow rate: 1.2mL/min
Instrument: SHIMADZU LC-2030C

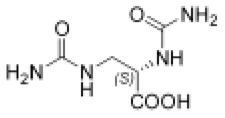

**KB13-S**

Chemical Formula: $C_5H_{10}N_4O_4$

Molecular Weight: 190.16

<Chromatogram>

ENBB240469-002-A1-HPLC D:\acdata\2024\0709\LC-6\ENBB240469-002-A1-HPLC_1.lcd

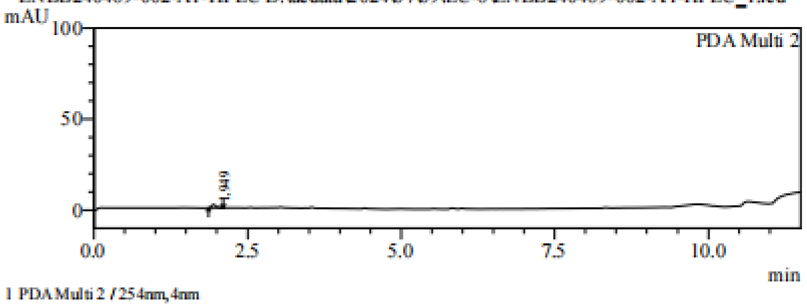

1 PDA Multi 2 / 254nm,4nm

<Result>

PeakTable

PDA Ch2 254nm

| Peak# | Ret. Time | Area | Height | Area % |
|---|---|---|---|---|
| 1 | 1.949 | 8709 | 1705 | 100.000 |
| Total | | 8709 | 1705 | 100.000 |

### LCMS

```
Acq. Operator   : ADMIN
Acq. Instrument : LCMS-K                              Location : P1-B-05
Injection Date  : 7/8/2024 2:32:20 PM                      Inj :    1
                                                    Inj Volume : 1.000 µl
Acq. Method     : D:\CHEMSTATION\1\METHODS\P50-600.M
Last changed    : 6/29/2024 11:19:46 AM by ADMIN
Analysis Method : C:\CHEM32\1\METHODS\DEF_LC.M
Last changed    : 11/20/2006 6:14:44 PM
Additional Info : Peak(s) manually integrated
```

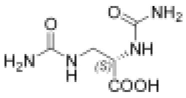

KB13-S

Chemical Formula: $C_5H_{10}N_4O_4$

Molecular Weight: 190.16

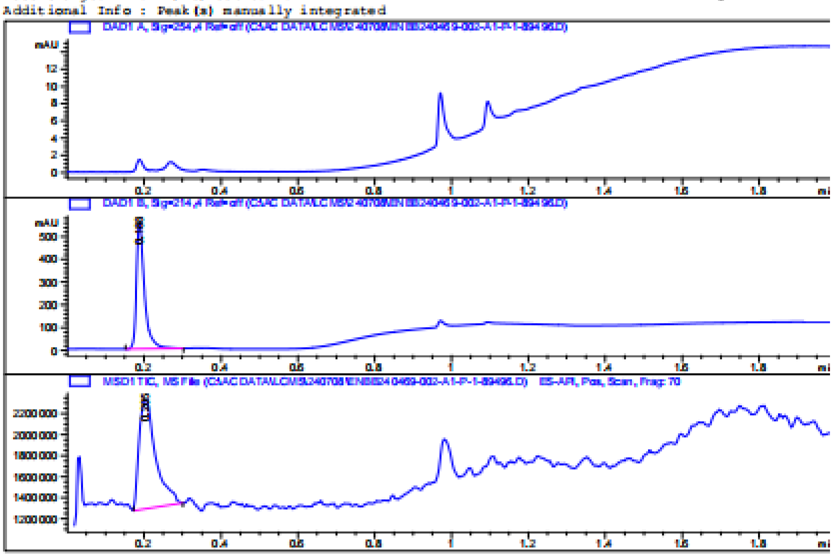

Area Percent Report

```
Sorted By             :      Signal
Multiplier:                  :      1.0000
Dilution:                    :      1.0000
Use Multiplier & Dilution Factor with ISTDs
```

Signal 1: DAD1 A, Sig=254,4 Ref=off

Signal 2: DAD1 B, Sig=214,4 Ref=off

| Peak # | RetTime [min] | Type | Width [min] | Area [mAU*s] | Height [mAU] | Area % |
|---|---|---|---|---|---|---|
| 1 | 0.188 | BB | 0.0219 | 806.34711 | 555.85394 | 100.0000 |
| Totals : | | | | 806.34711 | 555.85394 | |

Signal 3: MSD1 TIC, MS File

| Peak # | RetTime [min] | Type | Width [min] | Area | Height | Area % |
|---|---|---|---|---|---|---|
| 1 | 0.205 | BB | 0.0456 | 3.14280e6 | 1.04461e6 | 100.0000 |
| Totals : | | | | 3.14280e6 | 1.04461e6 | |

*** End of Report ***

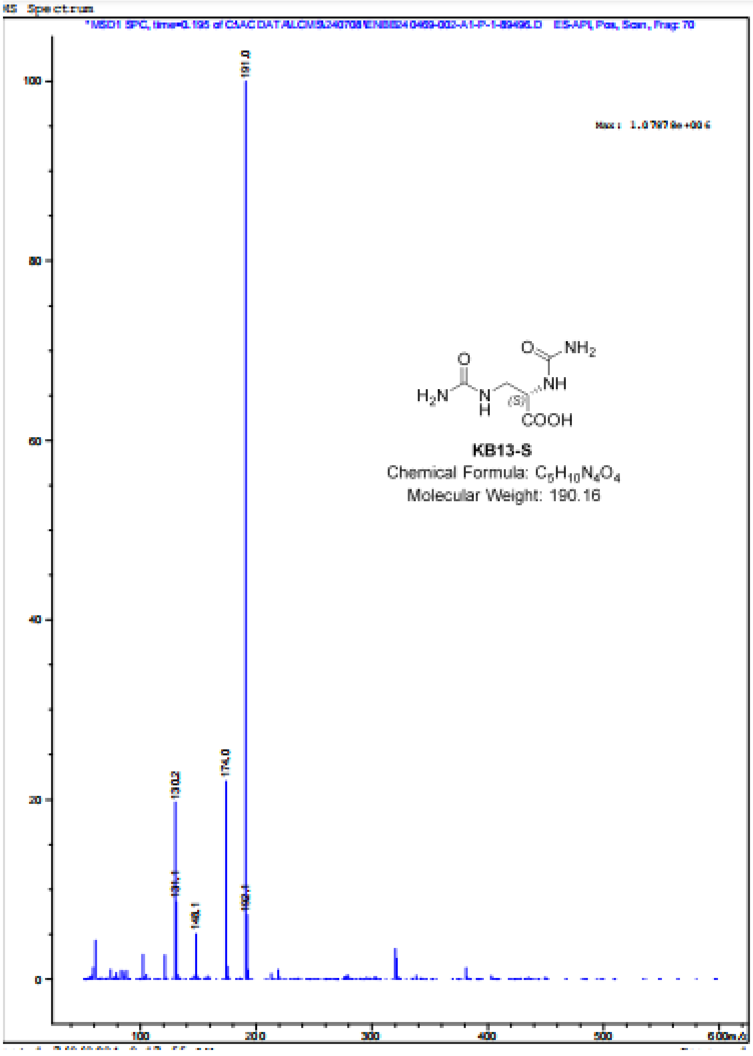
MS Spectrum
ES-API, Pos, Scan, Frag: 70
191.0
174.0
130.2
131.1
148.1
192.1
KB13-S
Chemical Formula: C5H10N4O4
Molecular Weight: 190.16
COOH
NH
NH2
H2N
(S)
100
80
60
40
20
0
100
200
300
400
500
600

### HNMR

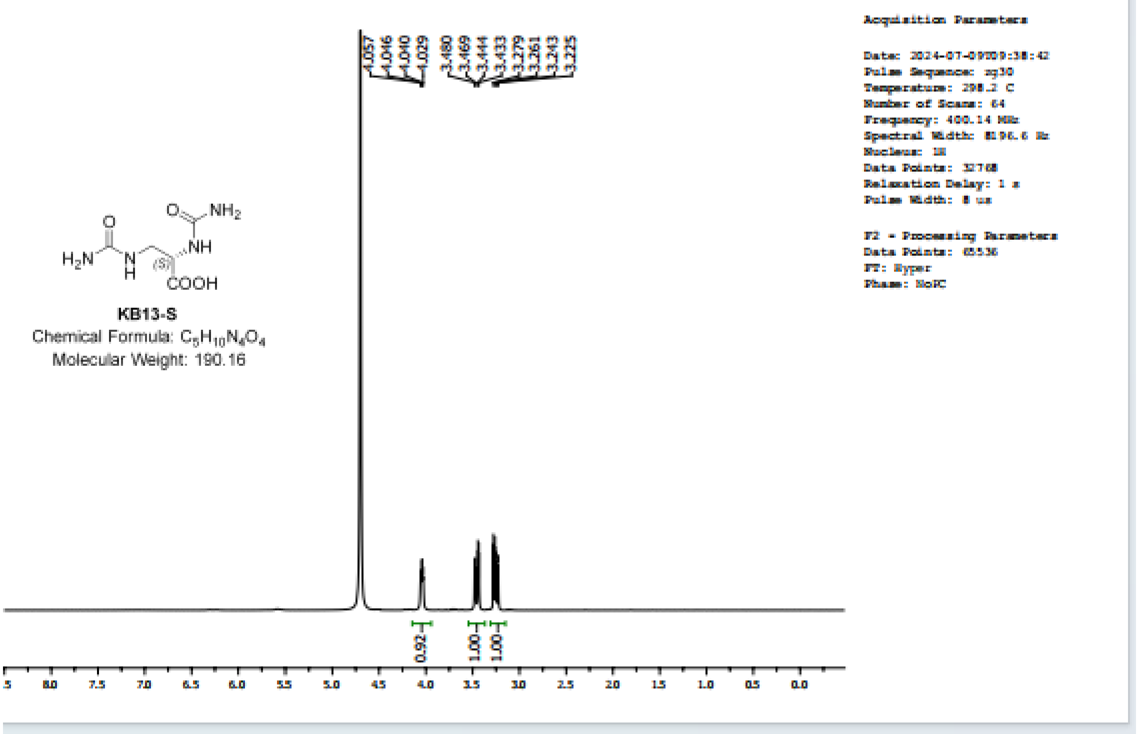
4.057
4.046
4.040
4.029
3.480
3.469
3.444
3.433
3.279
3.261
3.243
3.225
KB13-S
Chemical Formula: $C_5H_{10}N_4O_4$
Molecular Weight: 190.16
0.92
1.00
1.00
8.0
7.5
7.0
6.5
6.0
5.5
5.0
4.5
4.0
3.5
3.0
2.5
2.0
1.5
1.0
0.5
0.0
Acquisition Parameters
Date: 2024-07-09T09:38:42
Pulse Sequence: zg30
Temperature: 298.2 C
Number of Scans: 64
Frequency: 400.14 MHz
Spectral Width: 8196.6 Hz
Nucleus: 1H
Data Points: 32768
Relaxation Delay: 1 s
Pulse Width: 8 us
F2 - Processing Parameters
Data Points: 65536
FT: Hyper
Phase: NoPC

#### Certificate of Analysis

**Compound Name:** (R)-3-amino-2-ureidopropanoic acid TFA salt

**Physical Description** gray solid

**Chemical Structure**

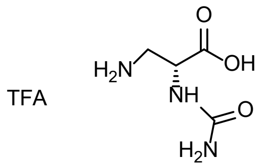

**Empirical Formula** $C_6H_{10}F_3N_3O_5$

**Molecular Weight** 261.06

**Purity** >95%

**HNMR** Confirmed

**Batch No.** KB20240626XRWR

#### Storage Conditions

lndividual variation in chemical stability profiles, do occur and for this reason we recommend adherence to the minimum recommended storage conditions. Because most compounds are custom made and shipped upon completion, long-term stability data is not available in most cases. If data is available, a specific recommendation will be provided below. Chemist cannot provide a guarantee of the long-term chemical stability of any compound.

Mlnimum Recommended Storage Conditions:

- Samples should be stored in an air tight vial
- Samples should be stored at≤ 20° Celsius when not in use for a prolonged period of time
- Samples should be stored in the dark or in an amber vial.

#### Caution

This information is provided as an indication of the quality of the underlying material when examined by a specific technique. The reported values are subject to normal experimental error and should be treated as estimates. The absence of undetected impurities cannot be guaranteed by this or any other general approach and this certificate does not certify the absence of such substances in the sample.

#### lntended Use

This product is intended for investigational use only and should not be used in humans. It is pharmaceutically unrefined, may contain uncharacterized toxic impurities, and is not intended for use in humans. Responsibility for its use and compliance with all federal laws rests solely with the purchaser.

# MS

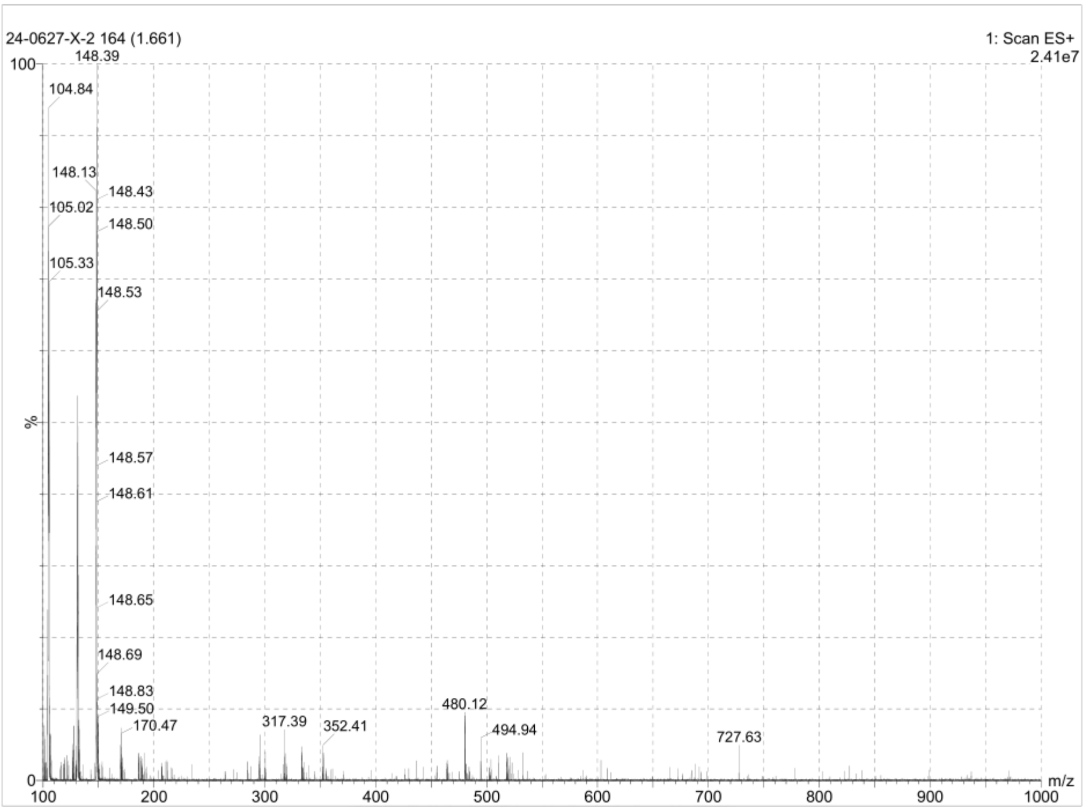
24-0627-X-2 164 (1.661)
1: Scan ES+
2.41e7
148.39
104.84
148.13
148.43
105.02
148.50
105.33
148.53
148.57
148.61
148.65
148.69
148.83
149.50
170.47
317.39
352.41
480.12
494.94
727.63
%
m/z
100
0
100
200
300
400
500
600
700
800
900
1000

### HNMR

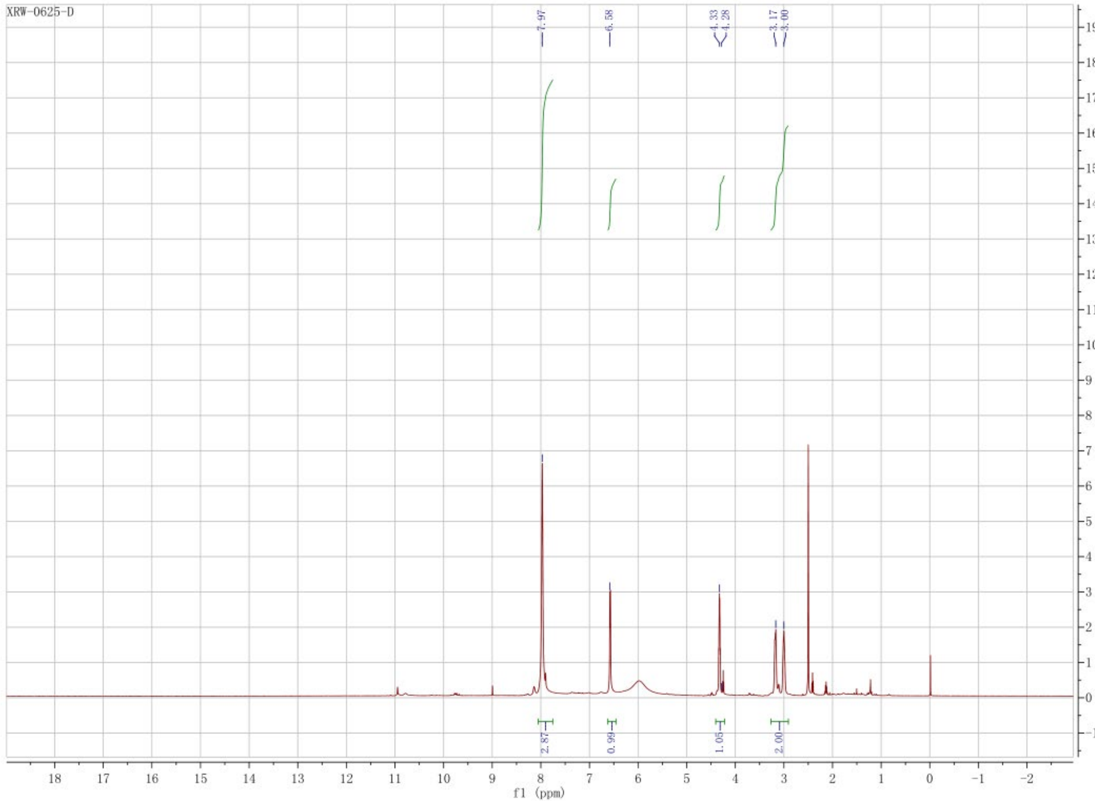

### Certificate of Analysis

**Compound Name：** (S)-3-amino-2-ureidopropanoic acid TFA salt

**Physical Description** gray solid

**Chemical Structure**

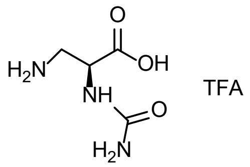

**Empirical Formula** $C_6H_{10}F_3N_3O_5$

**Molecular Weight** 261.06

**Purity** >95%

**HNMR** Confirmed

**Batch No.** KB20240626XRWS

#### Storage Conditions

lndividual variation in chemical stability profiles, do occur and for this reason we recommend adherence to the minimum recommended storage conditions. Because most compounds are custom made and shipped upon completion, long-term stability data is not available in most cases. If data is available, a specific recommendation will be provided below. Chemist cannot provide a guarantee of the long-term chemical stability of any compound.

Mlnimum Recommended Storage Conditions:

- Samples should be stored in an air tight vial
- Samples should be stored at≤ 20° Celsius when not in use for a prolonged period of time
- Samples should be stored in the dark or in an amber vial.

#### Caution

This information is provided as an indication of the quality of the underlying material when examined by a specific technique. The reported values are subject to normal experimental error and should be treated as estimates. The absence of undetected impurities cannot be guaranteed by this or any other general approach and this certificate does not certify the absence of such substances in the sample.

#### lntended Use

This product is intended for investigational use only and should not be used in humans. It is pharmaceutically unrefined, may contain uncharacterized toxic impurities, and is not intended for use in humans. Responsibility for its use and compliance with all federal laws rests solely with the purchaser.

# MS

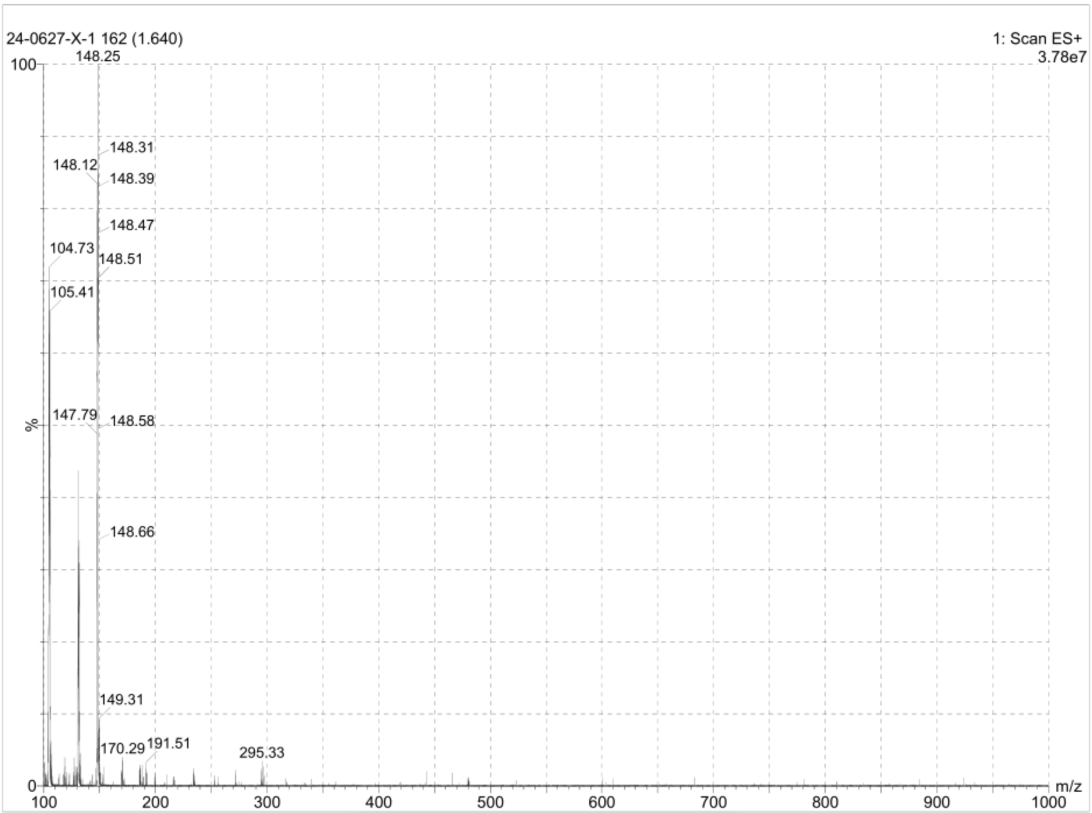
24-0627-X-1 162 (1.640)
1: Scan ES+
3.78e7
148.25
148.31
148.12
148.39
148.47
104.73
148.51
105.41
147.79
148.58
148.66
149.31
170.29
191.51
295.33
%
m/z
100
0
100
200
300
400
500
600
700
800
900
1000

### HNMR

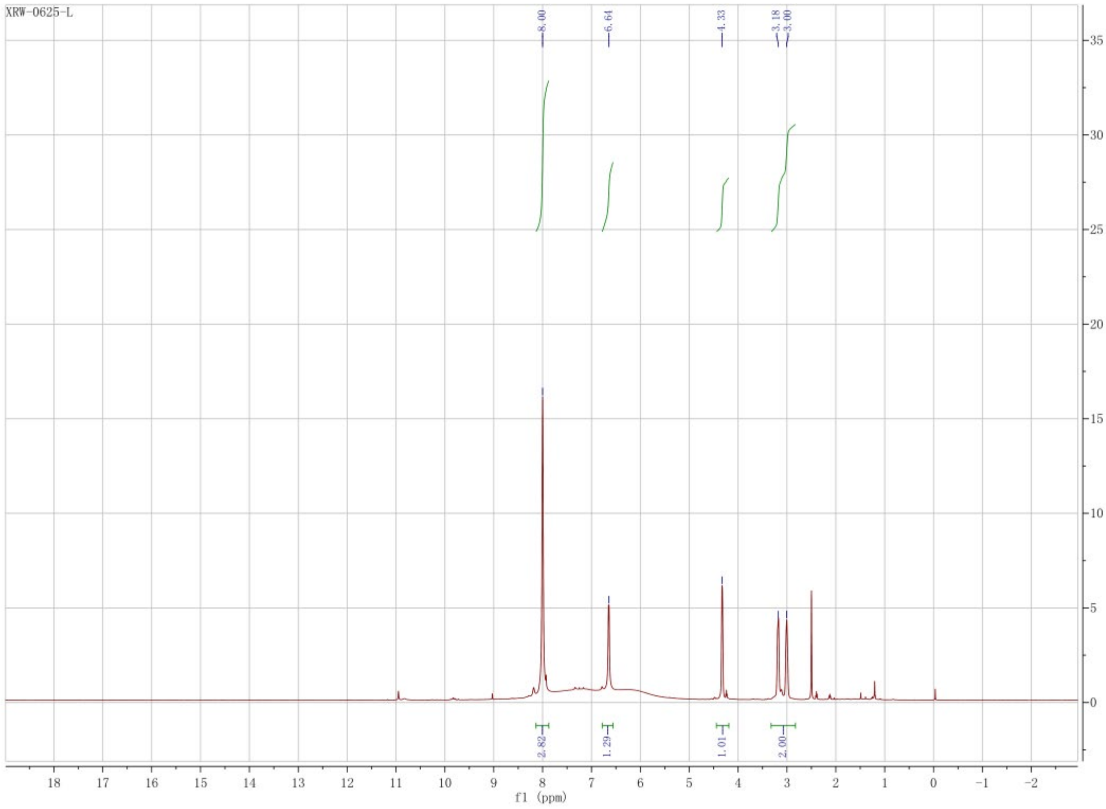
